# Cathepsin K Mediates the Formation of Potential Rheumatoid Arthritis-Relevant Cis- and Trans-Spliced Peptides Compatible With HLA-DR4 Presentation

**DOI:** 10.64898/2026.09.06.749279

**Authors:** Olivier Hinse, S. Yasin Tabatabaei Dakhili, Lee Freiburger, Eliot Mar, Jason Rogalski, Sriram Subramaniam, Eddie A. James, Leonard J. Foster, Dieter Brömme

## Abstract

Rheumatoid arthritis (RA) is characterized by a loss of immunological tolerance to synovial self-proteins, yet the initial triggers generating novel neo-antigens remain incompletely defined. Here, we demonstrate that human cathepsin K (hCatK), a key cysteine protease driving joint degradation in RA, catalyzes covalent cis- and trans-splicing of peptides from major RA-associated self-proteins and foreign antigens, including type II collagen, fibrinogen, and SARS-CoV-2 Spike protein. Using high-resolution LC-MS/MS and database-assisted *de novo* sequencing, we identified over 90 unique spliced peptides. Splicing efficiency peaked at near-neutral pH (6.5-7.5), contrasting with classic hydrolytic profiles. Biochemical profiling revealed strong subsite selectivity, with a striking enrichment for small, aliphatic and/or hydroxyl-containing residues (Gly, Thr, Ser) at the P1 position. Furthermore, splicing preferentially targeted flexible, intrinsically disordered protein regions, with 81% of fibrinogen splicing events clustering within its αC domain. *In silico* binding predictions for the RA-susceptibility allele HLA-DRB1*04:01 harboring the shared epitope revealed that numerous hCatK- generated spliced peptides exhibit predicted affinities exceeding those of established immunogenic and genomic sequences, uncovering protease-mediated transpeptidation as a novel post-translational modification capable of generating potent MHC class II autoantigens in RA.

## INTRODUCTION

Rheumatoid arthritis (RA) is a chronic autoimmune disease characterized by severe joint inflammation and destruction (1, 2). Several hypotheses have been proposed to explain how the immune system breaches immunological tolerance against self-proteins within the synovium, including molecular mimicry, post-translational modifications (PTMs) such as citrullination, and epitope spreading via neo-antigens generated during progressive joint erosion (3, 4). Genetic susceptibility further supports a central role for antigen presentation in RA pathogenesis, evidenced by the strong association between disease risk and the shared epitope alleles of HLA-DRB1 (5). Among the self-proteins targeted by the autoimmune response in RA, including vimentin, fibrinogen, type II collagen (TIIC), and α-enolase, citrullinated forms in particular are linked to the induction of autoantibody and autoreactive T cells (4, 6–8). The HLA-DRB1 shared epitope is strongly associated with the presence of anti-citrullinated protein antibodies (ACPAs) in patients (9, 10). However, ACPAs often precede clinical symptoms by years or even decades, leaving open the question of whether yet-unidentified PTMs act as the initial trigger for disease induction (11).

Cysteine cathepsins are key mediators of antigenic peptide processing in antigen-presenting cells (APCs), which in RA include not only classical dendritic cells and B-cells but also synovial fibroblasts, macrophages, and osteoclasts (12–15). Beyond their canonical role in endo-lysosomal peptide generation, cathepsins are also active extracellularly during matrix degradation (16–18). Peptides generated in the extracellular space of the joint remain viable for MHC class II presentation via peptide exchange, provided they have sufficient affinity (19). Among cysteine cathepsins, human cathepsin K (hCatK) is a major driver of joint pathology and disease progression in RA (20–22). Within the joint, hCatK is expressed by osteoclasts, synovial macrophages, and synovial fibroblasts, where it degrades cartilage and bone tissue matrix components such as collagens and aggrecans (23). TIIC is a well-established substrate of hCatK in the synovium, and fibrinogen/fibrin has similarly been proposed as a natural substrate (24, 25). The SARS-CoV-2 Spike protein has separately been implicated as a contributing factor in RA (26), and we show here that it is cleaved by hCatK. hCatK is thus uniquely positioned at the interface of the intra- and extracellular spaces relevant to RA.

We recently demonstrated that cysteine cathepsins, including hCatK, can generate cis- and trans-spliced peptides (27). We define cis-spliced peptides as products resulting from the fusion of two non-contiguous sequences from the same protein substrate, either from the same molecule or two distinct ones, and trans-spliced peptides as those resulting from the fusion of sequences from two distinct protein substrates including foreign components such as the SARS-CoV-2 Spike protein. We therefore hypothesized that cathepsin-mediated peptide splicing could contribute to the generation of novel autoantigens.

The primary objective of this study is to extend this peptide splicing mechanism to RA pathology and to comprehensively characterize the specificity of hCatK in mediating this PTM.

## RESULTS

### Transpeptidation between small RA-related peptides by hCatK

To investigate the *in vitro* transpeptidation capability of hCatK, the fluorogenic FRET substrate Ac-AE(EDANS)PGKAGEGGK-Dabcyl (TIIC1) was co-incubated with the nucleophilic donor peptide TSTCitGGK-Biotin (V1) in the presence of 500 nM hCatK at pH 6.5. The FRET substrate incorporates a human TIIC-derived hCatK cleavage site (GEPG↓KAGE; residues 612–619 of the TIIC α1 chain) predicted by the PACMANS software (28). The V1 peptide encompasses residues 33–36 of human vimentin and features a citrullinated residue; this sequence is predicted to be generated via hCatK cleavage at the SYVT↓TSTR motif. A citrullinated variant of this vimentin sequence was previously identified as an immunogenic peptide capable of inducing CD4^+^ T-cell proliferation in anti-citrullinated protein antibody-positive (ACPA^+^) and shared epitope-positive (SE^+^) patients (29).

Following co-incubation at pH 6.5 with hCatK, LC-MS/MS and *de novo* sequencing identified several cleavage and spliced peptides, including 10 trans-spliced products and two V1-derived cis-spliced products (**Figure 1A**). Notably, the primary trans-spliced peptide T1 (Ac-AE(EDANS)PGTSTCitGGK-Biotin) underwent secondary hydrolysis to yield trans-spliced peptide T2 (Ac-AE(EDANS)PGTSTCit). Formation of the subsequent fusions with full-length V1 (T3–T5) required that the parental FRET substrate be cleaved at two additional sites before undergoing successive secondary hydrolysis. Furthermore, hCatK mediated the N-terminal hydrolytic truncation of V1 by one to four amino acids, yielding trans-spliced peptides T6–T10 via one or two distinct FRET substrate-derived thioacyl intermediates and subsequent secondary hydrolysis. Two cis-spliced products derived from V1 (C1 and C2) were also detected (**Figure 1A**). These results provided the first confirmation that hCatK could catalyze the synthesis of many spliced peptide products from a very limited number of original small peptide substrates.

**Figure 1.**
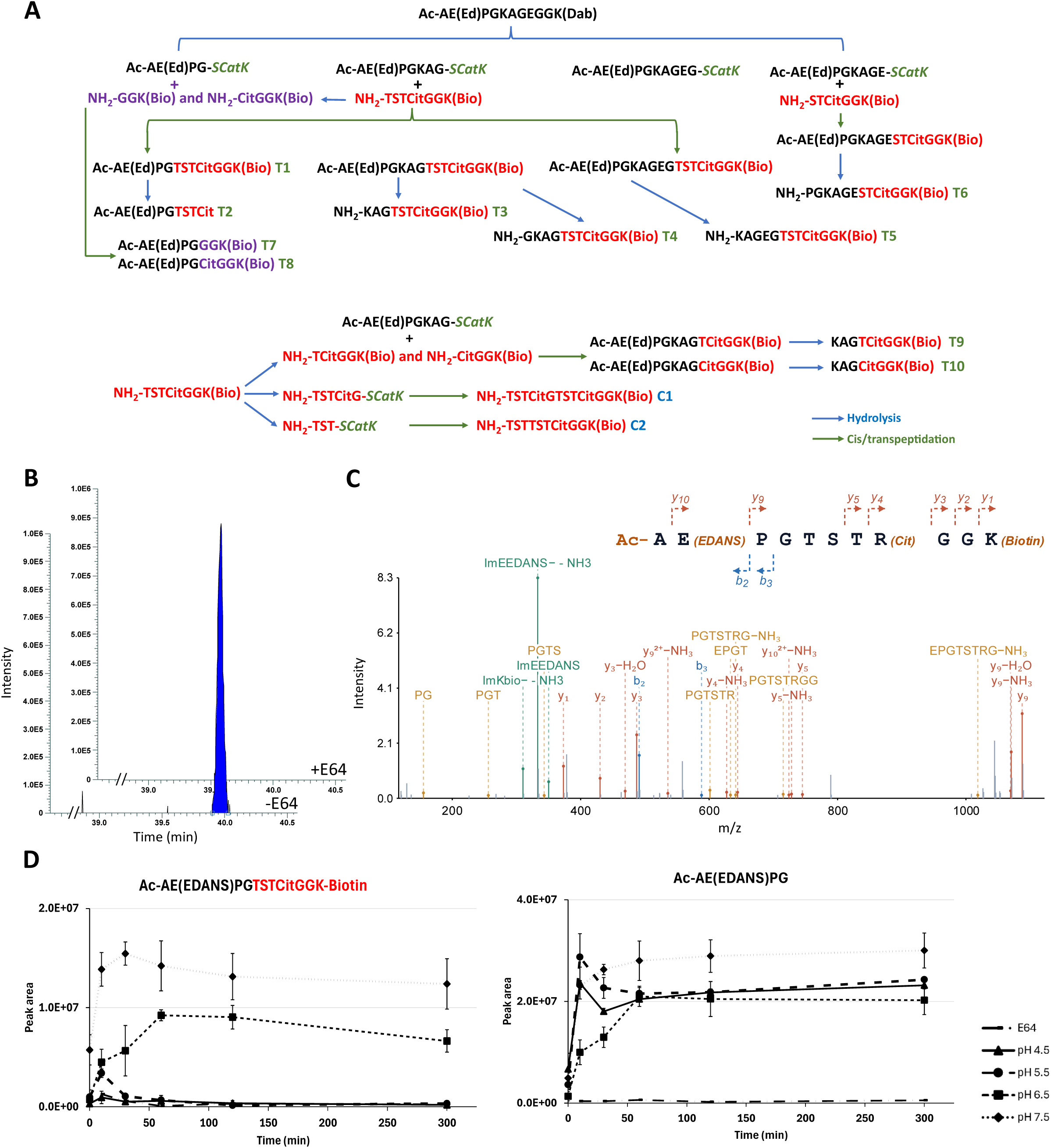
Characteristics of transpeptidation between two RA-related small peptide sequences. **A)** Schematic illustration of a mechanistically plausible pathway for the formation of the 12 spliced peptides generated between TIIC1 and V1 during coincubation with 500 nM hCatK. **B)** Extracted ion chromatograms of the trans-spliced peptide AcAE(EDANS)PGTSTCitGGK(Biotin) in the presence and absence of E64. **C)** Annotated tandem mass spectrum of AcAE(EDANS)PGTSTCitGGK(Biotin). **D)** Peak area over time for the transpeptidation product AE(EDANS)PGTSTCitGGK(Biotin) formed between TIIC1 and V1, and for the cleavage product AE(EDANS)PG derived from TIIC1, measured across four pH conditions. Values represent three independent experiments; error bars indicate SD.

Product identification and validation were confirmed by tracking diagnostic signature fragment ions derived from the EDANS and Biotin tags in the tandem mass spectra (**Figures 1C and S1**). In the presence of the pan-cysteine cathepsin inhibitor E64, all hydrolysis and fusion activities were abolished (**Figure 1B**).

Given that pH varies across physiological compartments, from acidic lysosomes to slightly alkaline extracellular matrix (ECM) environments, and this influences splicing efficiency, we quantified the amounts of the major initial cleavage product Ac- AE(EDANS)PG and the T1 spliced product (Ac-AE(EDANS)PGTSTCitGGK-Biotin) across a pH range from 4.5 to 7.5. Minimal spliced product was formed at pH 4.5 and 5.5; however, its synthesis was very noticeable at pH 6.5 and 7.5 (**Figure 1D**). The levels of all identified cleavage and spliced peptides observed across pH values are presented in **Figure S2**. Overall, the amounts of spliced peptides were undetectable or very low at pH 4.5-5.5, but reached very high detection levels at pH 6.5-7.5, while the levels of the cleavage products identified remained similar across the whole pH gradient, except for PGKAGE which presented lower levels at pH 4.5. These results indicate that splicing is favored at a neutral pH over acidic conditions.

These elevated rates of peptide splicing were further corroborated by evaluating the k_cat_/K_m_ values of TIIC1 hydrolysis across increasing concentrations of the nucleophilic V1 peptide and elevated pH values. Because the nucleophilic donor directly competes with water for the covalent acyl-enzyme acceptor intermediate, transpeptidation in the presence of excess biotinylated donor peptide should suppress fluorophore re-quenching by preventing the partial reformation of the initial FRET substrate. Consistent with this mechanism, a 1.9-fold increase in the apparent k_cat_/K_m_ was captured at pH 7.5, when compared to pH 5.5, in the presence of 200 μM V1 (**Table 1**).

**Table 1.**
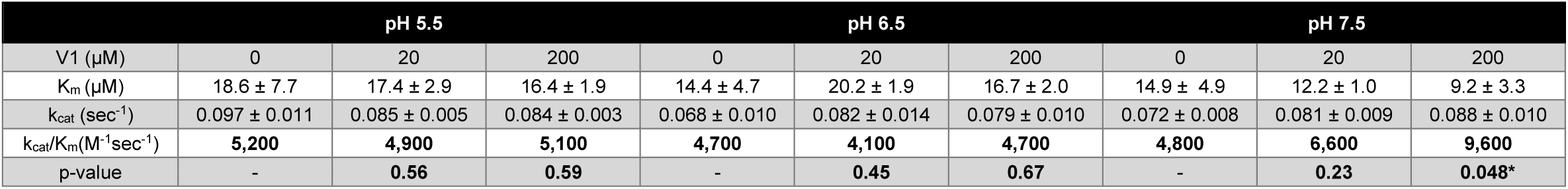
Michaelis-Menten calculations obtained for hCatK in differing pH and V1-containing environments. Kinetic parameters (k_cat_, K_M_, and k_cat_/K_M_) for hCatK digestion of TIIC1 under three pH conditions and three concentrations of V1. Values are reported as mean ± SD from three independent biological triplicates. P-values were calculated relative to the no- V1 condition (0 µM V1) at each pH.

These results, obtained with a simplified peptide model, confirm that hCatK acts as a transpeptidase and that splicing is favored at pH 6.5 and 7.5, leading us to examine this in a more complex system.

### Peptide splicing between full-length RA-related proteins and a small vimentin-derived peptide

Building on the RA-related model peptides experiments, we expanded our experimental framework to evaluate full-length, native RA-associated protein substrates in combination with the V1 donor peptide. Full-length human fibrinogen and denatured bovine TIIC (DTIIC) were co-incubated with hCatK across different pH values in the presence of the citrullinated and biotinylated vimentin-derived donor peptide, V1.

First, we rigorously verified whether the observed trans-spliced products bearing the V1 peptide were indeed true covalent adducts. No fusion products were detected either in the absence of hCatK or when the protease was blocked by the pan-cathepsin inhibitor E64. Similarly, incubating pre-digested DTIIC or fibrinogen with V1, in the presence of inactivated hCatK, yielded no hybrid products. Biotinylated fusion products emerged only when the active acceptor protein was co-incubated with V1 in the presence of catalytically active hCatK (**Figure 2A**). This strict dependence on active hCatK, coupled with the complete absence of products upon enzyme inhibition or prior substrate pre-digestion, unambiguously validates that these hybrid molecules form via a *bona fide* covalent acyl-enzyme intermediate rather than through non-enzymatic peptide association. These proof-of-concept experiments were performed at pH 6.5.

**Figure 2.**
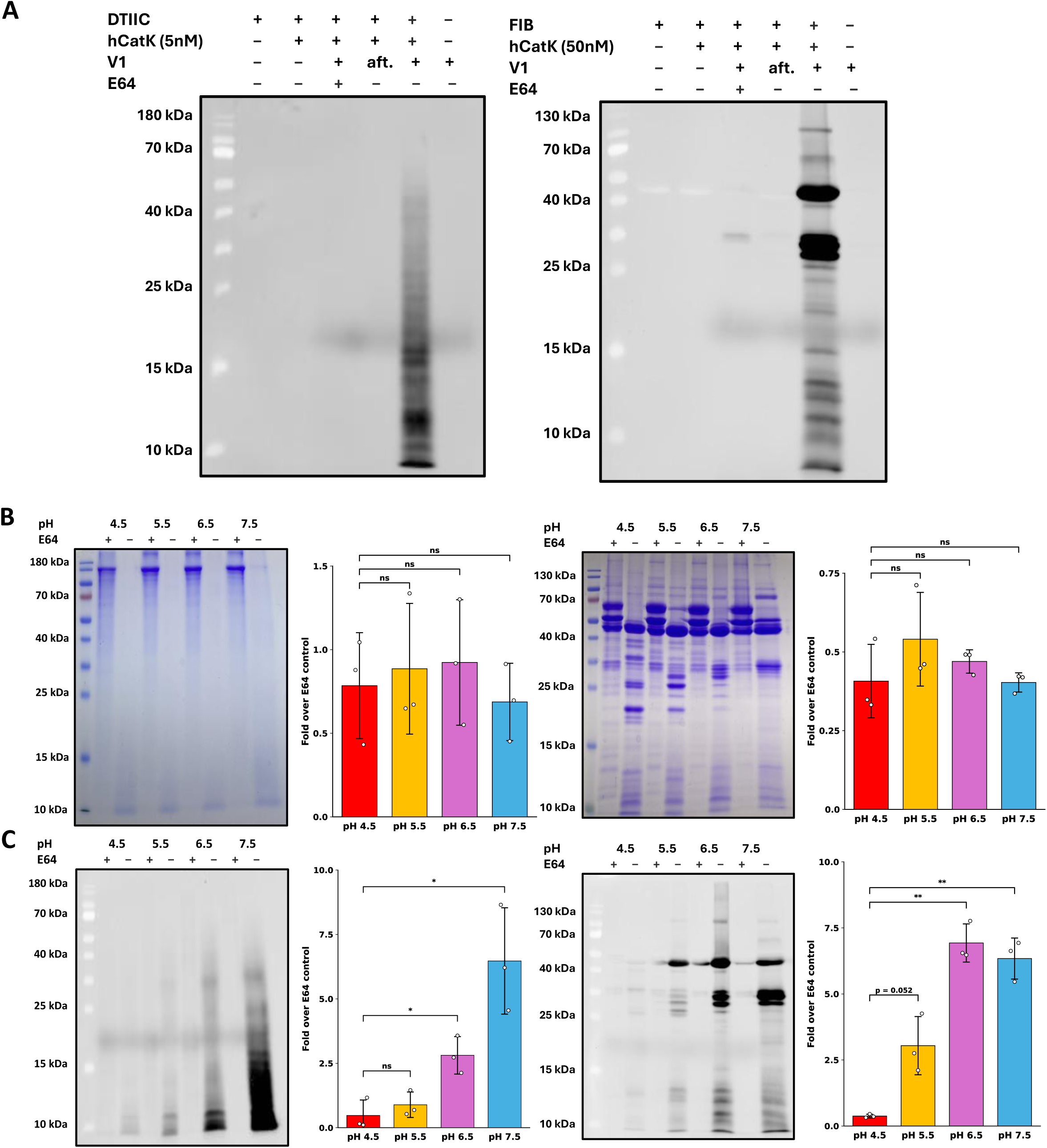
Transpeptidation between full-length RA-related protein substrates and a biotinylated vimentin-derived peptide (V1). **A)** Schematic illustration of the formation of biotinylated hybrid products, inferred from avidin-FITC–stained bands observed for four substrates (from left to right: DTIIC, fibrinogen, Spike S1, and Spike S2) following co-incubation with V1 and hCatK. **B-C)** Quantification of DTIIC (left) and fibrinogen (right) degradation (**B**) and transpeptidation (**C**) by hCatK across a pH gradient. Band intensities from Coomassie-stained gels (**B**) and avidin-FITC-stained membranes (**C**) were quantified using ImageJ and normalized to the E64 control at each pH; aft. (after digestion). Values represent three independent experiments; error bars indicate SD. Statistical comparisons were performed against pH 4.5 using Welch’s t-test (ns, not significant; *p < 0.05; **p < 0.01; ***p < 0.001).

To capture the physiological pH gradient hCatK encounters as it transits from the slightly alkaline ECM to the acidic lysosome, we next monitored degradation and splicing patterns of DTIIC and fibrinogen, each with V1, across a pH range of 4.5 to 7.5 using SDS-PAGE and avidin-FITC blotting. Cleavage efficacy, visualized by Coomassie blue staining, trended with the known pH profile of hCatK, exhibiting optimal hydrolytic potency between pH 5.5 and 6.5, although pairwise comparisons across the pH range did not show significant differences (**Figure 2B**) (30). Intriguingly, the formation of trans-spliced peptides generated between fragments of the full-length proteins and the V1 peptide showed an increase with rising pH values across the range tested (**Figure 2C**). This trend indicates that deprotonation of the α-amino group at higher pH values strongly favors the transpeptidation pathway over hydrolysis. Notably, transpeptidation also occurred efficiently with high molecular weight fragments of the acceptor protein.

To evaluate donor sequence constraints, we tested a second biotinylated peptide, V2 (ETNLGGK-Biotin), which encompasses an alternative antigenic epitope within the native human vimentin sequence (residues 425–428) (29). Interestingly, V2 was two to three times less efficient at splicing compared to V1, suggesting sequence preferences for donor peptides (**Figure S3**).

To confirm the identities of these spliced molecules, trans-spliced peptides generated between the full-length proteins and V1 were rigorously validated using high-resolution LC-MS/MS coupled with *de novo* peptide sequencing. For the DTIIC–V1 system, a total of 12 unique trans-spliced peptides were confidently assigned (**Table 2**, **Figures S4–S5 & S18A**). Also, we mapped 64 unique trans-spliced peptides formed between fibrinogen fragments and V1 (**Table 3**, **Figures S6–S7 & S18B–T**).

**Table 2.**
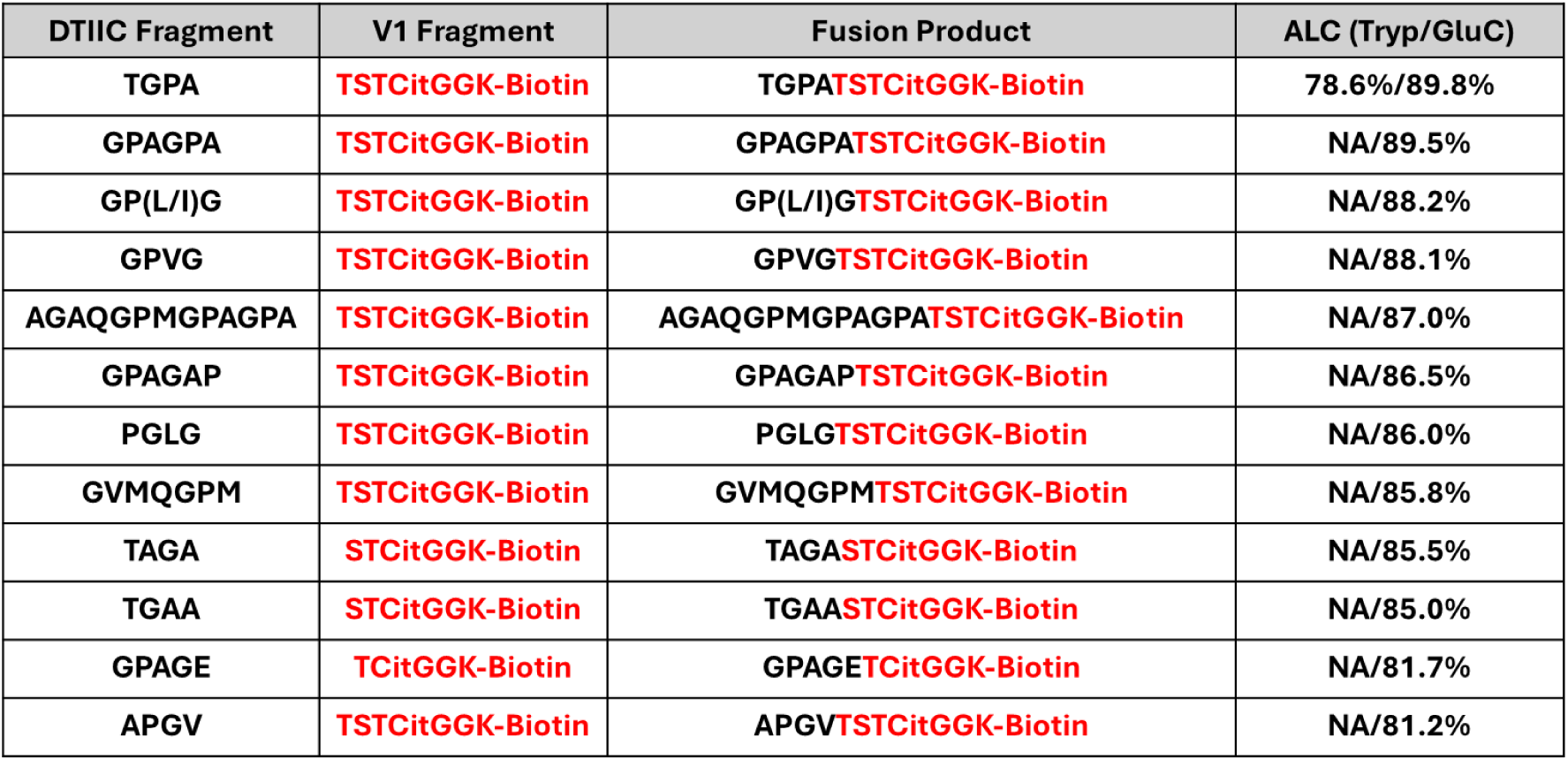
Trans-spliced peptides catalyzed by hCatK between DTIIC & V1 fragments. Trans-spliced peptides were identified by regular de novo peptide sequencing in PEAKS Studio 12.5 with ALC ≥ 80% and orthogonal verification by diagnostic biotin fragment ions (m/z 227.0849 and 310.1580), and were absent from all E64-inhibited controls. Columns list: acceptor sequence (from DTIIC, black), donor sequence (from V1, red), full trans-spliced peptide sequence, ALC (%) obtained for each monoplicate (Trypsin and GluC, NA indicates the peptide was not detected in that digestion condition). Cit = citrullination; L/I indicates leucine/isoleucine ambiguity inherent to de novo sequencing.

**Table 3.**
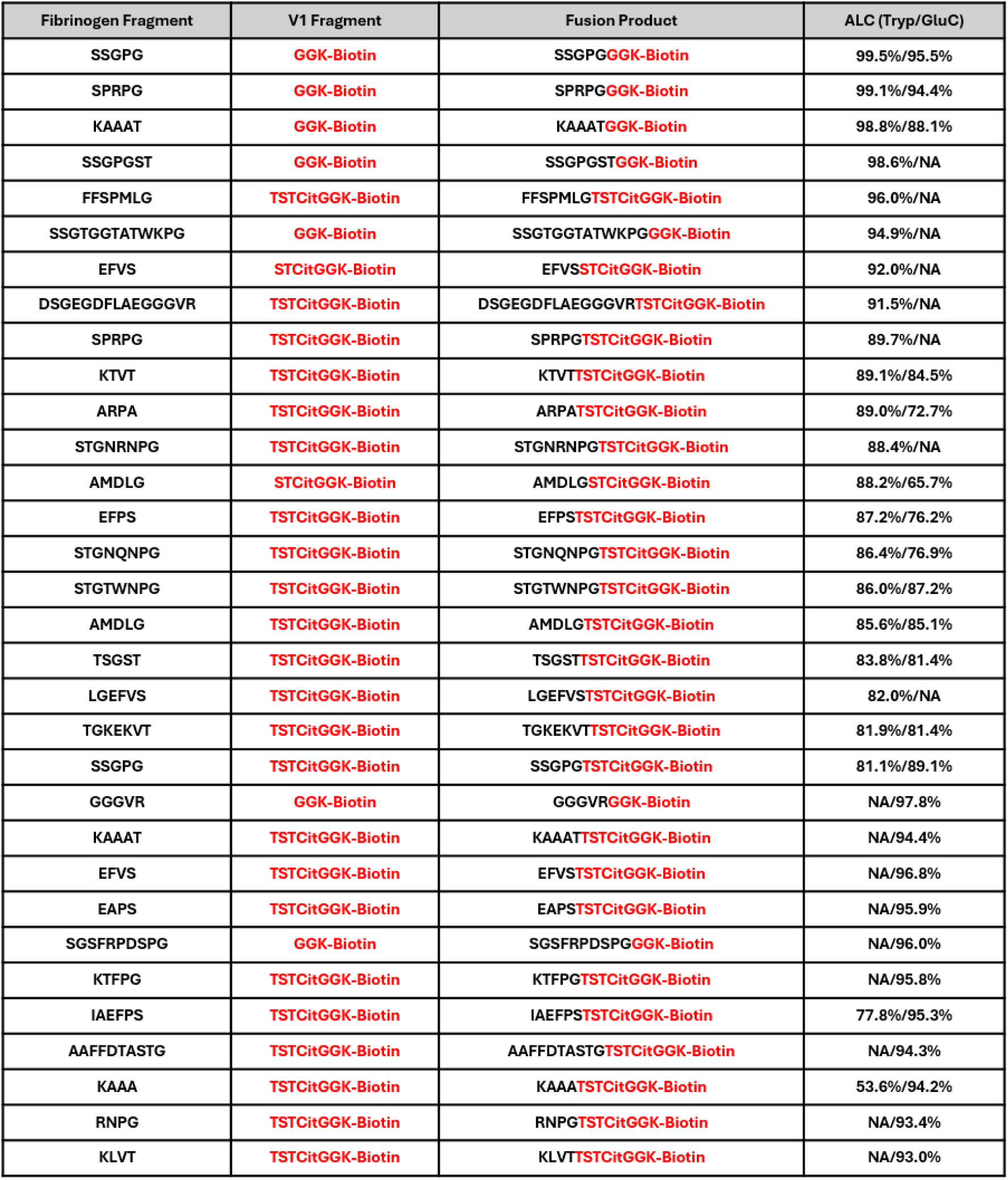
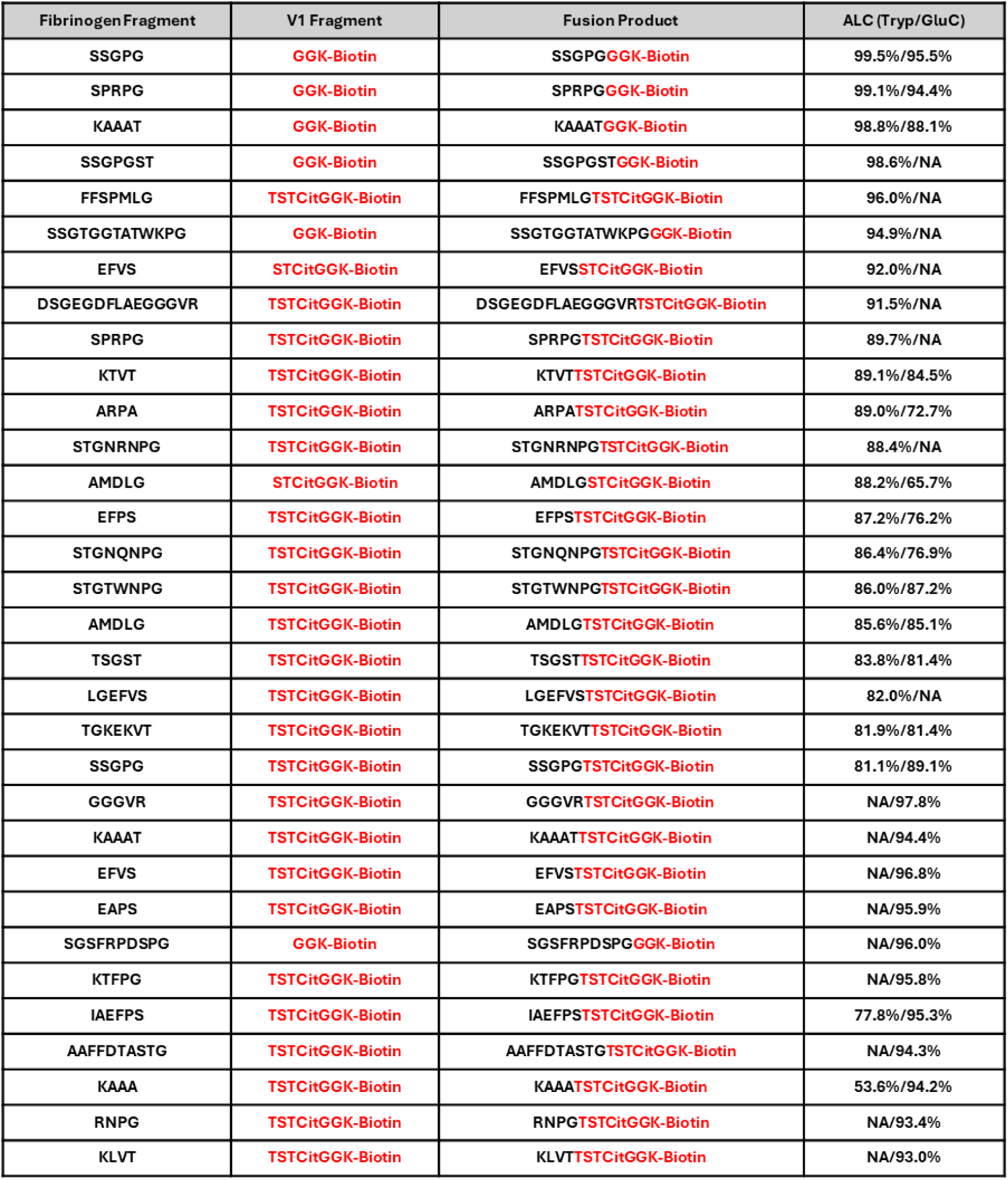
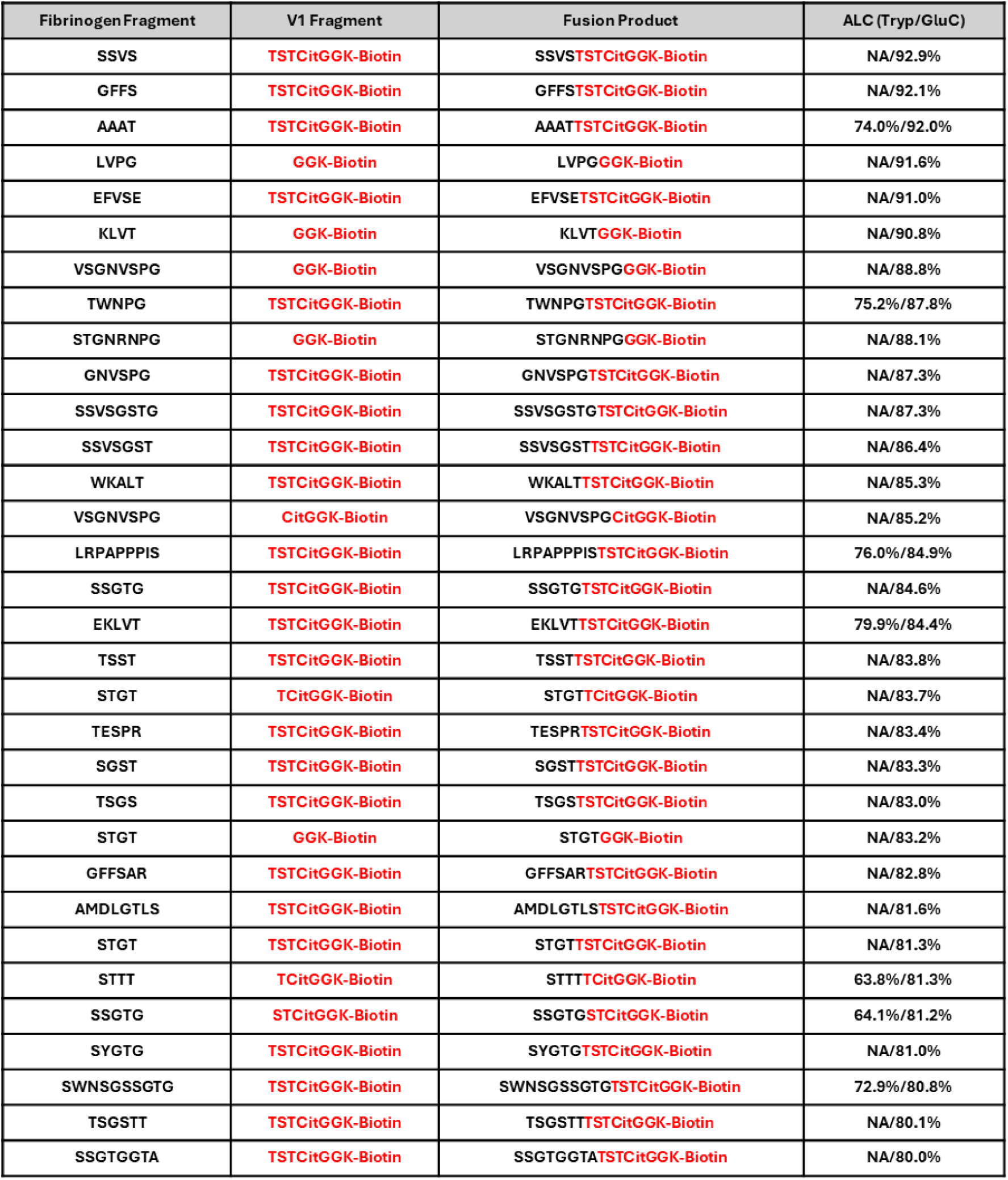
Trans-spliced peptides catalyzed by hCatK between fibrinogen & V1 fragments. Trans-spliced peptides were identified by regular de novo peptide sequencing in PEAKS Studio 12.5 with ALC ≥ 80% and orthogonal verification by diagnostic biotin fragment ions (m/z 227.0849 and 310.1580), and were absent from all E64-inhibited controls. Columns list: acceptor sequence (from fibrinogen, black), donor sequence (from V1, red), full trans-spliced peptide sequence, ALC (%) obtained for each monoplicate (Trypsin and GluC, NA indicates the peptide was not detected in that digestion condition). Cit = citrullination.

Collectively, these results build on the simplified peptide model by demonstrating covalent splicing between RA-associated protein fragments and V1, while confirming the splicing reaction is favored at near-neutral pH, unlike hydrolysis.

### Peptide splicing between full-length RA-related proteins and subsite specificity

Because physiologically relevant splicing is unlikely to always depend on a pre- formed exogenous donor peptide, we sought to demonstrate the autonomous formation of cis- and trans-spliced peptides between fragments from full-length, RA-associated proteins (fibrinogen and DTIIC). Following the incubation of human fibrinogen alone (comprising the FGA, FGB, and FGG chains) with hCatK, we discovered 69 unique cis-spliced peptides. These events were partitioned into 50 intra-FGA, 1 intra-FGG, and 18 inter-subunit splicing events (**Table 6 & Figures S12–S13**). Analogous digests yielded 17 unique cis-spliced peptides for DTIIC (**Table 5 & Figures S10–S11**). A subset of these spliced peptides underwent manual spectral validation (summarized in **Figure S19**). These events are presented in **Figure 3A**.

**Figure 3.**
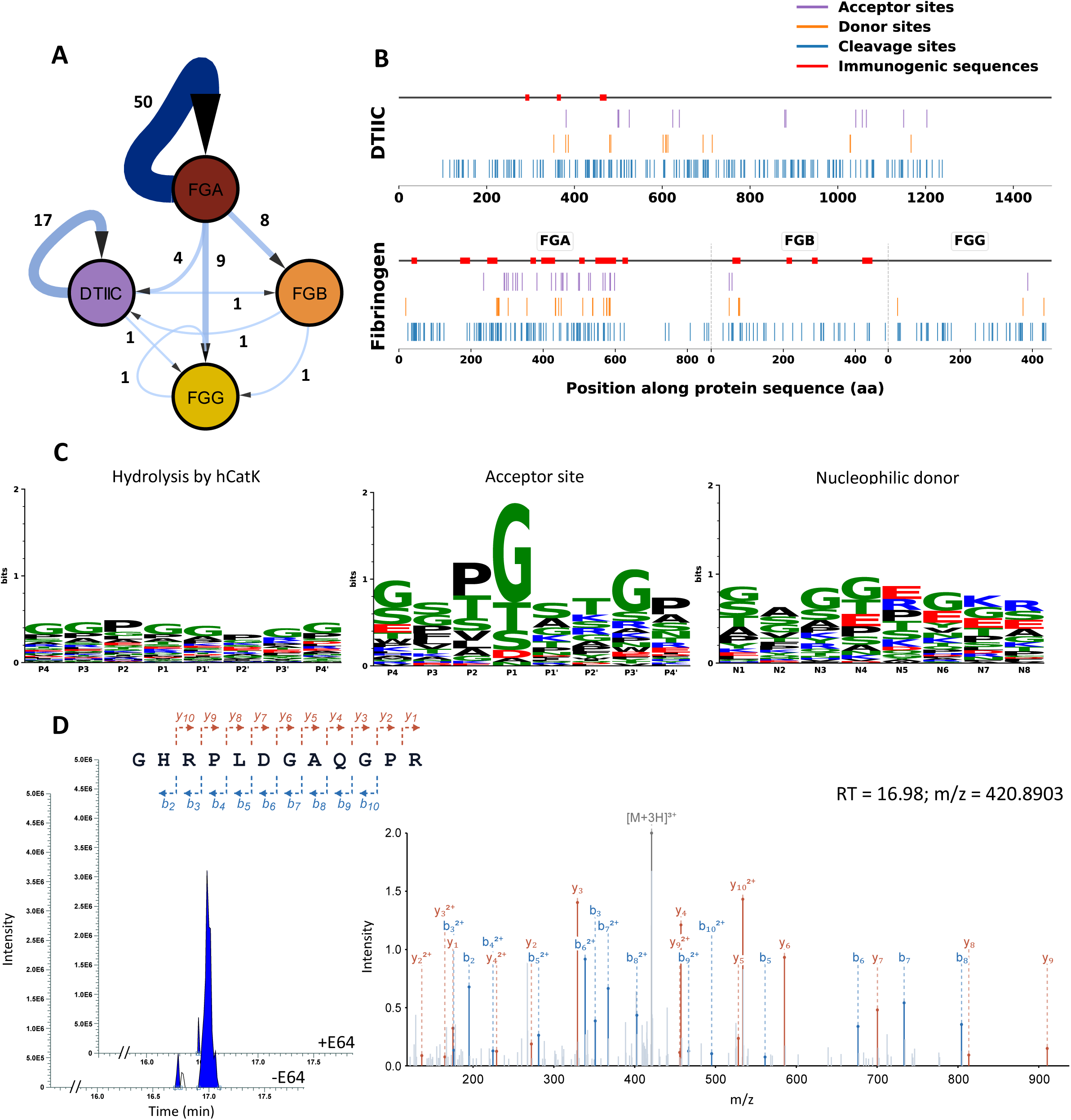
Cis- and transpeptidation patterns observed between full-length RA-related protein substrates b tandem mass spectrometr . **A)** Network quantitatively illustrating the amount of splicing events between two RA-related substrates: DTIIC, fibrinogen (FGA, FGB, and FGG). **B)** Map illustrating the sites of cleavage and splicing (acceptor and donor/nucleophilic sites) by hCatK in RA-related substrates. **C)** Cleavage and splicing patterns observed following hCatK activity on DTIIC and fibrinogen. **D)** Integrated ion chromatogram and annotated tandem mass spectrum of G RPLDGAQGPR, a trans-spliced peptide identified between a fibrinogen fragment and a DTIIC fragment, following hCatK activity (50nM of enzyme). The trans-spliced peptide is not observed in the presence of E64, an hCatK inhibitor.

To push the boundaries of multi-protein splicing dynamics, fibrinogen and DTIIC substrates were co-incubated in the presence of hCatK. This dual-substrate system yielded 7 high-confidence interprotein trans-spliced peptides with an Average Local Confidence score (ALC) > 90%. Structurally, five of these hybrid products featured a nucleophilic donor fragment from DTIIC fused to a corresponding acceptor fragment from fibrinogen. The remaining two products resulted from a fibrinogen-derived donor fragment directly attacking a DTIIC-derived acyl-enzyme intermediate (**Table 8**, **Figures S16–S17 & S20**). The events are schematically presented in **Figure 3A**. These results show hCatK can splice peptide products autonomously within and between native RA-associated proteins.

The tandem mass spectra of two cis-spliced and one trans-spliced products were further validated by comparison to the spectra of synthetic versions of the same products. Comparison revealed identical precursor ion m/z, RT values, and fragmentation patterns, with cosine similarity scores of 0.80 (cis-spliced in fibrinogen), 0.77 (cis-spliced in DTIIC), and 0.74 (trans-spliced between DTIIC and fibrinogen). The cosine scores are reduced by non-relevant background peaks, with the most abundant features matching with high fidelity (**Figure S21**).

Mapping these intra-protein targeting coordinates back to their primary sequences provided structural insights. While hCatK-mediated hydrolysis occurred indiscriminately across most domains, certain segments remained highly resistant to cleavage, such as the C-terminal domain of DTIIC (**Figure 3B**, in blue). Splicing acceptor (purple) and donor (yellow) loci appeared sporadically across the two model proteins; this lower density of splicing events likely stems from the rapid, competing degradation of the substrates. However, within the FGA chain, the fibrinogen subunit yielding the highest density of splicing events, the acceptor sites clustered tightly within a localized region in the middle of the subunit (**Figure 3B**, in orange). Structural profiling mapped this “hyper-splicing hotspot” directly to an intrinsically disordered region (IDR): specifically, the highly flexible αC-connector domain that tethers the αC-domain to the core protein (31). A similar preference for flexible segments was observed, though to a lesser degree, within the disordered N-terminal region of the FGB chain (**Figure 3B**). **Figure 3D** depicts a tandem mass spectrum of a representative hCatK-generated trans-spliced peptide between a fibrinogen fragment and a DTIIC fragment. Validating the enzymatic nature of this mechanism, this trans-spliced peptide was completely absent in the presence of E64. Overall, these findings suggest that splicing by hCatK is favored in disordered, flexible regions of fibrinogen.

To delineate the structural and sequence constraints governing hCatK-mediated splicing, the biochemical features of all captured events (intra-protein cis-splicing and inter-protein trans-splicing) were pooled and analyzed. The sequence conservation patterns surrounding sites targeted for cis–transpeptidation versus hydrolytic cleavage reveal a striking contrast (**Figure 3C**). As anticipated, hydrolysis by hCatK displayed high variability and limited sequence specificity across the different substrates, reflecting its broad substrate specificity (as depicted in the MEROPS database; **Figure 3C**). Indeed, DTIIC and fibrinogen exhibited rather broad cleavage profiles lacking any globally shared motifs (**Figures S5 & S7**).

In sharp contrast, the cis–transpeptidation dataset unveiled a highly conserved target sequence preference. This pathway was characterized by a dramatic enrichment of small aliphatic or hydroxyl-containing residues (Gly, Thr, Ser) at the P1 position in the acceptor site (**Figure 3C**). Small polar (Thr, Ser) and hydrophobic (Pro, Leu, Val) residues were similarly recurrent at the P2 position. The donor sequences supplying the incoming amine nucleophile (positions N1–N4) to attack the acyl-enzyme intermediate also showed a distinct preference for smaller residues, such as Gly, Ser, Thr, and Ala, though to a lesser degree (**Figure 3C**). Small residues were not favored at the more distal N5-N8 positions to the same extent as at N1-N4.

### SARS-CoV-2 Spike protein as donor and acceptor in hCatK-catalyzed cis- and transpeptidation

The Spike protein of SARS-CoV-2 has been implicated as a trigger of RA after viral infection (26). Therefore, we evaluated the possibility of forming trans-spliced peptides between Spike and the V1 peptide, as well as with DTIIC and fibrinogen as typical RA-associated endogenous antigens. We incubated the Spike subunits S1 and S2 separately with V1 and noticed the formation of trans-spliced products (**Figure 4A**). Altogether, we identified 26 distinct splicing events between S1/S2 fragments and V1 (**Table 4**, **Figures S8–S9 & S18U–AI**) where 20 were derived from S1 and 6 from S2. Incubating the combined S1/S2 with full-length DTIIC or fibrinogen interestingly resulted in no trans-spliced peptides with an ALC over 90% and only two, between Spike and DTIIC, with an ALC over 80% (**Table 8**, **Figures 4C-D**, **S16-S17 & S20D-E**). Furthermore, two cis-spliced peptides with an ALC over 90% and specific to the S1 and S2 subunits were also observed and manually confirmed (**Table 7 & Figures 4B, S14–S15**). These results indicate that splicing by hCatK can generate neo-sequences from SARS-CoV-2 Spike, which offers a potential biochemical rationale for post-viral RA.

**Figure 4.**
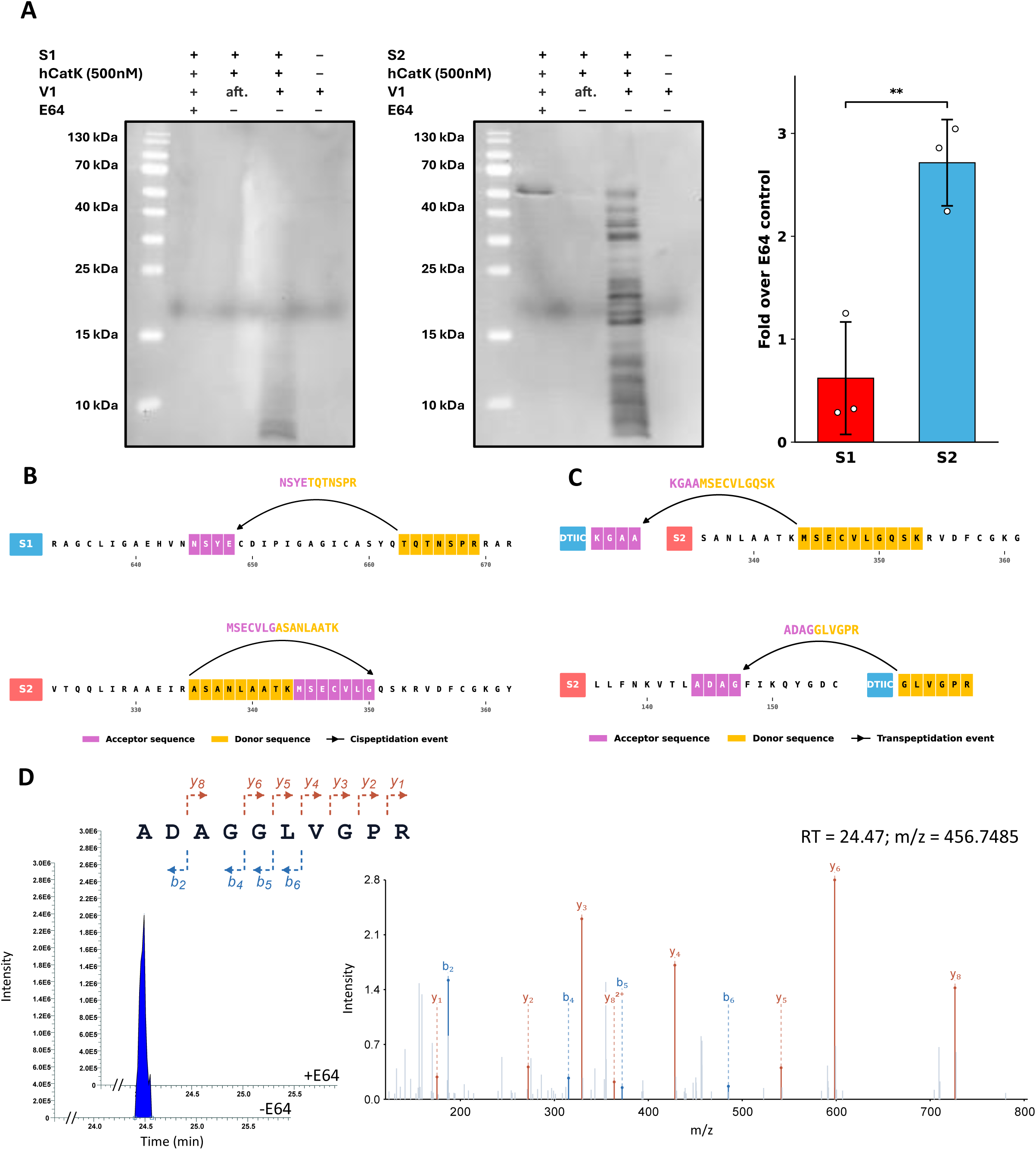
Cis- and transpeptidation with the SARS-CoV-2 Spike protein. **A)** Comparison and quantification of transpeptidation patterns observed for Spike subunits (S1 and S2) co-incubated with V1. Band intensities from avidin-FITC-stained membranes were quantified using ImageJ; aft. (after digestion). Values represent three independent experiments; error bars indicate SD. Statistical comparisons were performed against the E64 control using Welch’s t-test (ns, not significant; *p < 0.05; **p < 0.01; ***p < 0.001). **B-C)** Schematic illustration of the formation of cis- (**B**) and trans-spliced (**C**) peptides with Spike subunits. (**D**) Integrated ion chromatogram and annotated tandem mass spectrum of **ADAGGLVGPR**, a trans-spliced peptide identified between a Spike fragment and a DTIIC fragment, following hCatK activity (500nM of enzyme). The trans-spliced peptide is not observed in the presence of E64, an hCatK inhibitor.

**Table 4.**
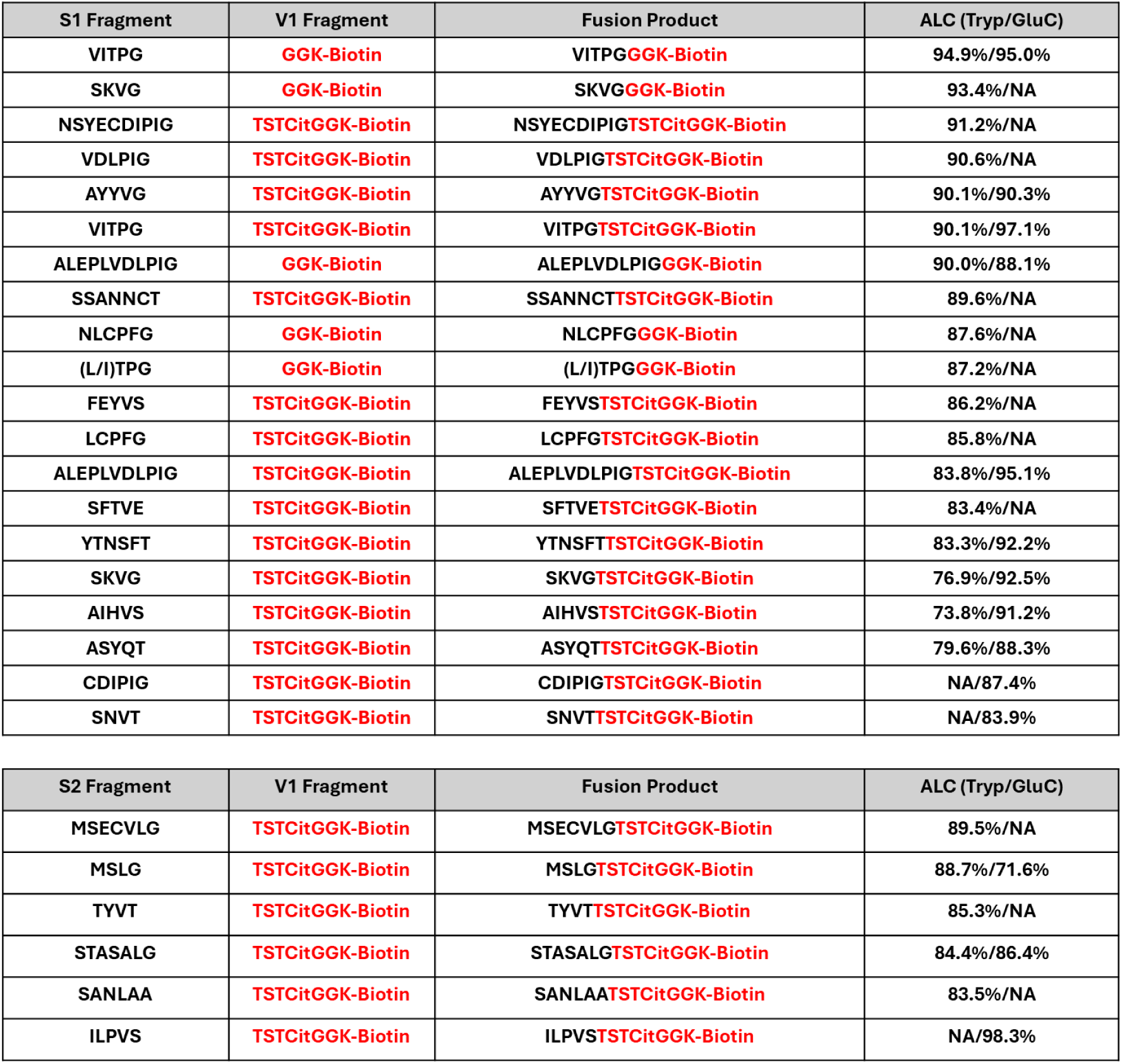
Trans-spliced peptides catalyzed by hCatK between Spike & V1 fragments. Trans-spliced peptides were identified by regular de novo peptide sequencing in PEAKS Studio 12.5 with ALC ≥ 80% and orthogonal verification by diagnostic biotin fragment ions (m/z 227.0849 and 310.1580), and were absent from all E64-inhibited controls. Columns list: acceptor sequence (from Spike, black), donor sequence (from V1, red), full trans-spliced peptide sequence, ALC (%) obtained for each monoplicate (Trypsin and GluC, NA indicates the peptide was not detected in that digestion condition). Cit = citrullination; L/I indicates leucine/isoleucine ambiguity inherent to de novo sequencing.

**Table 5.** Cis-spliced peptides catalyzed by hCatK between DTIIC fragments. Cis-spliced peptides were identified by database-assisted de novo peptide sequencing in PEAKS Studio 12.5, excluding all matches to reference proteomes, with ALC ≥ 90%, detected in at least 2 of 3 replicates, and absent from all E64- inhibited controls. Columns list: acceptor sequence (black), donor sequence (red), full cis-spliced peptide sequence, average ALC score across replicates for trypsin and GluC digestions (Tryp/GluC; NA indicates the peptide was not detected in that digestion condition, %), standard deviation of ALC (SD ALC, %), average retention time (RT, min), standard deviation of RT (SD RT, min), average peak area (arbitrary units), standard deviation of peak area (SD Peak Area, arbitrary units), and number of replicates in which the peptide was detected (N). Protein abbreviation: DTIIC = denatured bovine type II collagen.

| DTIIC Fragment 1 | DTIIC Fragment 2 | Fusion Product | Average ALC<br>(Tryp/GluC) | SD (ALC) | Average<br>RT | SD (RT) | Average<br>Peak Area | SD (Peak<br>Area) | Replicates<br>(N) |
| --- | --- | --- | --- | --- | --- | --- | --- | --- | --- |
| GPSGD | PGKAGE | GPSGDPGKAGE | NA/99.0% | 0.3% | 10.18 | 2.15 | 4.55x10<sup>5</sup> | 1.83x10<sup>5</sup> | 3 |
| AGEKGLPG | GANGEPGK | AGEKGLPGGANGEPGK | 98.4%/NA | 0.4% | 17.16 | 0.25 | 4.61x10<sup>5</sup> | 2.29x10<sup>5</sup> | 2 |
| EGSPG | GPAGEPGR | EGSPGGPAGEPGR | 97.4%/NA | 1.7% | 13.41 | 0.10 | 9.78x10<sup>6</sup> | 5.33x10<sup>6</sup> | 3 |
| KGLPG | GANGEPGKAGE | KGLPGGANGEPGKAGE | NA/96.2% | 0.0% | 17.71 | 0.04 | 5.42x10<sup>5</sup> | 3.09x10<sup>4</sup> | 2 |
| SGETGPAGPPG | GPPGPVGPSGK | SGETGPAGPPGGPPGPVGPSGK | 95.5%/NA | 2.1% | 24.83 | 0.05 | 6.91x10<sup>6</sup> | 3.74x10<sup>6</sup> | 3 |
| GFPGQDGLAG | APGPAGEEGK | GFPGQDGLAGAPGPAGEEGK | 94.6%/NA | 3.0% | 31.00 | 0.03 | 2.56x10<sup>6</sup> | 5.05x10<sup>4</sup> | 2 |
| GPTG | GAQGPR | GPTGGAQGPR | 94.2%/NA | 0.9% | 13.50 | 0.02 | 1.49x10<sup>5</sup> | 1.44x10<sup>4</sup> | 2 |
| DGETG | LVGPR | DGETGLVGPR | 94.0%/NA | 1.3% | 26.64 | 0.01 | 5.27x10<sup>5</sup> | 1.30x10<sup>5</sup> | 3 |
| GLPG | GFPGPK | GLPGGFPGPK | 93.7%/NA | 0.1% | 32.52 | 0.01 | 5.45x10<sup>5</sup> | 4.31x10<sup>4</sup> | 2 |
| GEPGG | ANGEPGK | GEPGGANGEPGK | 93.6%/NA | 0.0% | 18.25 | 0.03 | 7.96x10<sup>4</sup> | 1.40x10<sup>4</sup> | 2 |
| SGETGPAGPPG | PVGPSGK | SGETGPAGPPGPVGPSGK | 92.3%/NA | 1.0% | 25.56 | 0.03 | 2.14x10<sup>6</sup> | 7.47x10<sup>4</sup> | 2 |
| DGAAGVKG | PAGEPGR | DGAAGVKGPAGEPGR | 91.5%/NA | 4.2% | 17.21 | 0.01 | 1.84x10<sup>6</sup> | 9.79x10<sup>5</sup> | 3 |
| GPTGVTG | LQGAR | GPTGVTGLQGAR | 90.6%/NA | 0.8% | 15.44 | 0.02 | 7.00x10<sup>6</sup> | 3.40x10<sup>6</sup> | 3 |
| TGAVG | GPAGEPGRE | TGAVGGPAGEPGRE | NA/90.3% | 3.8% | 18.11 | 0.05 | 2.96x10<sup>6</sup> | 1.44x10<sup>6</sup> | 3 |
| GPSGLAG | APGPAGEEGKR | GPSGLAGAPGPAGEEGKR | 90.2%/NA | 1.4% | 21.96 | 0.05 | 1.20x10<sup>6</sup> | 5.89x10<sup>5</sup> | 3 |
| GEPGGAG | PAGEEGKR | GEPGGAGPAGEEGKR | 90.2%/NA | 5.2% | 12.60 | 1.00 | 1.40x10<sup>7</sup> | 1.70x10<sup>7</sup> | 3 |
| GPPG | GPPGPVGPSGK | GPPGGPPGPVGPSGK | 90.2%/NA | 4.1% | 20.94 | 0.09 | 2.18x10<sup>7</sup> | 1.03x10<sup>7</sup> | 3 |

### Database-assisted *de novo* sequencing demonstrates robust spliced peptide identification capabilities

In order to validate the splicing events, our laboratory has established an analytical workflow optimized to screen for and identify cis- and trans-spliced peptides generated in low-complexity digestions containing discrete substrate-protease mixtures (27). In the present study, we expanded this computational architecture to chart peptide splicing driven by hCatK (**Figure 5A**). ALC score distributions were compared between the complete raw peptide dataset with a mean ALC ≥ 90% (n = 27,073) and the final population of validated spliced peptides with a mean ALC ≥ 90% (n = 95; **Figure 5B**). The spliced peptide population exhibited a significantly higher mean ALC profile relative to the baseline background peptide pool (Mann-Whitney p = 3.92 x 10^-20^, **Figure 5B**). A distribution analysis of all sequenced peptides identified by PEAKS Studio and our downstream analysis as potential spliced peptide candidates demonstrated that 90.8% of these raw *de novo* assignments scored below our 90% ALC filter threshold and were filtered out as unconfirmed (**Figure 5C**). Only 9.2% (144 unique candidates) successfully cleared this confidence threshold; among these, 95 unique hybrid sequences were reproducibly detected in ≥ 2 experimental replicates to establish the final pool of confirmed spliced peptides (**Figure 5C-D**). None of these peptides were found in the presence of the E64 inhibitor. It should be mentioned that the two identified Spike/DTIIC trans-spliced peptides had an ALC score below 90% and were not considered for downstream analysis. This suggests that our filtering steps reliably identify true spliced peptides, albeit conservatively, and likely underestimate the true extent of splicing.

**Figure 5.**
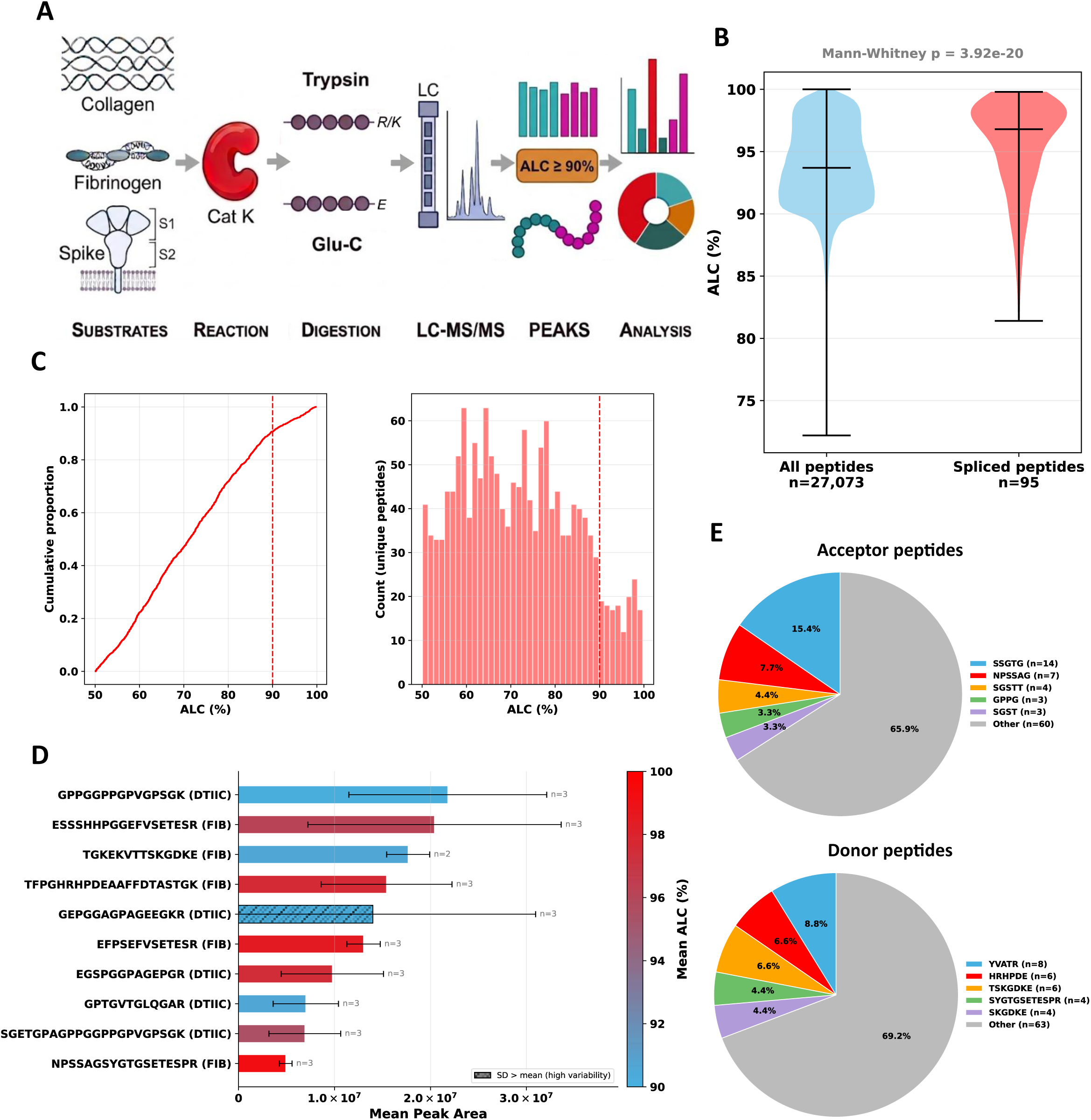
Characterization, workflow, and quality metrics of spliced peptides identification by LC-MS/MS analysis. **A)** Simplified workflow used to identify spliced peptides, for more details see Materials & Methods. **B)** Distribution of mean ALC scores for all peptides (blue) and confirmed spliced peptides (red) with a mean ALC ≥ 90%. **C)** Empirical cumulative distribution function (left) and histogram (right) depicting the distribution of spliced peptide candidates identified by database-assisted *de novo* sequencing in PEAKS. Candidates with a mean ALC below 90% (dashed line) were excluded from subsequent analyses as potential false positives; only those above this threshold were considered. **D)** Bar plot illustrating the top 10 unique spliced peptides presenting the highest mean peak area by LC-MS/MS, hatched boxes indicate peptides presenting high variability (SD > mean). **E)** Pie chart depicting the most common acceptor and donor peptides observed in unique spliced peptides across all substrates tested.

Among the ten confirmed spliced peptides with the highest mean peak areas across experimental replicates, nine were generated via tryptic digestion, while a single product was recovered following Glu-C endoproteinase digestion (**Figure 5D**). Globally, the median integrated peak area calculated across all identified peptides with a mean ALC ≥ 90% was 3.91 x 10^5^, which closely matched the median peak area determined for the confirmed spliced peptide population (4.27 x 10^5^). Globally, spliced peptides are generated at abundances comparable to conventional peptide products.

An alignment frequency analysis of the component fragments making up these verified spliced peptides demonstrated that three discrete acceptor sequences, SSGTG (n=14), NPSSAG (n=7), and SGSTT (n=4), accounted for 27.5% of all acceptor events. Concurrently, three discrete donor sequences, YVATR (n=8), HRHPDE (n=6), and TSKGDKE (n=6), comprised 22.0% of all nucleophilic donor events (**Figure 5E**). These localized pooling dynamics further indicate that hCatK-mediated splicing is a site-directed biochemical event favoring specific hotspots across self-proteins. The most highly represented acceptor sequences consist of threonine/serine/glycine-rich motifs displaying either a Gly or Thr residue at the P1 position, aligning with the preference patterns described previously.

However, a data-acquisition bias favoring peptides with more readily ionizable C-terminal sequences cannot be excluded.

### Transpeptidation could potentially lead to the creation of immunogenic spliced peptides

To draw inference about the immunogenic potential of these newly mapped hCatK- derived spliced products, we conducted *in silico* binding predictions targeting the RA-susceptible HLA-DRB1*04:01 allele, which pairs with the HLA-DRA*01:01 chain to present peptides to CD4+ T cells and has a well characterized binding motif. High-confidence hCatK-spliced peptides identified from tryptic or Glu-C digestions were extended into standardized 20-mer peptide models centered directly on the fusion junction (10 amino acids from the acceptor block fused to 10 amino acids from the donor block). This modeling architecture was selected to preserve the local structural and stereochemical context flanking the fusion site while standardizing peptide lengths for comparative docking. Binding predictions were performed using a well-defined *in silico* prediction matrix for the HLA-DRB1*04:01 receptor, previously published by our collaborators (32, 33).

As comparative controls, simulations were additionally performed using peptides derived from TIIC, fibrinogen, and vimentin that have been previously described immunodominant or capable of inducing T-cell proliferation in human or murine models (e.g., CIA mice) (34–37). These reference peptides comprised: citrullinated FGB_69-81_(Cit 74), a major immunodominant T-cell epitope in RA previously shown to be presented by HLA-DR4 (34); CII_259-273_ which harbors a well-characterized RA-related immunodominant T-cell epitope (35, 36); and VIM_65-77_(Cit 70), a high-affinity binder to the HLA-DR4 receptor (37). Among these controls, one of three reference peptides exhibited good predicted affinity for HLA-DR4, expressed as a relative binding affinity score (RBA): VIM_65-77_(Cit 70) (predicted RBA = 0.325), whereas the two others had lower predicted affinities, CII_259-273_ (predicted RBA = 0.093) and FGB_69-81_(Cit 74) (predicted RBA = 0.092), that are still in the same range as previously published epitopes (32, 38).

Strikingly, a large proportion of the confirmed cis- and trans-spliced peptides exhibited higher predicted binding affinity for the HLA-DR4 receptor than any of these established immunogenic controls, and 67 (59 cis-, 8 trans-) passed the RBA threshold at 0.4 above which peptides are considered very good predicted binders (**Figure 6A**). In total, 57 of these predicted good binders were fibrinogen-derived cis-spliced products, with two cis-spliced products from Spike, and none from collagen **(Figure 6B**). The top-ranked binders were AAFFDTASTGSGSTTTTRRS (predicted RBA = 4.229), SYSKQFTSSTEFVSETESRG (predicted RBA = 3.566), and AAFFDTASTGHRHPDEAAFF (predicted RBA = 2.795), each representing fibrinogen cis-spliced 20-mer products.

**Figure 6.**
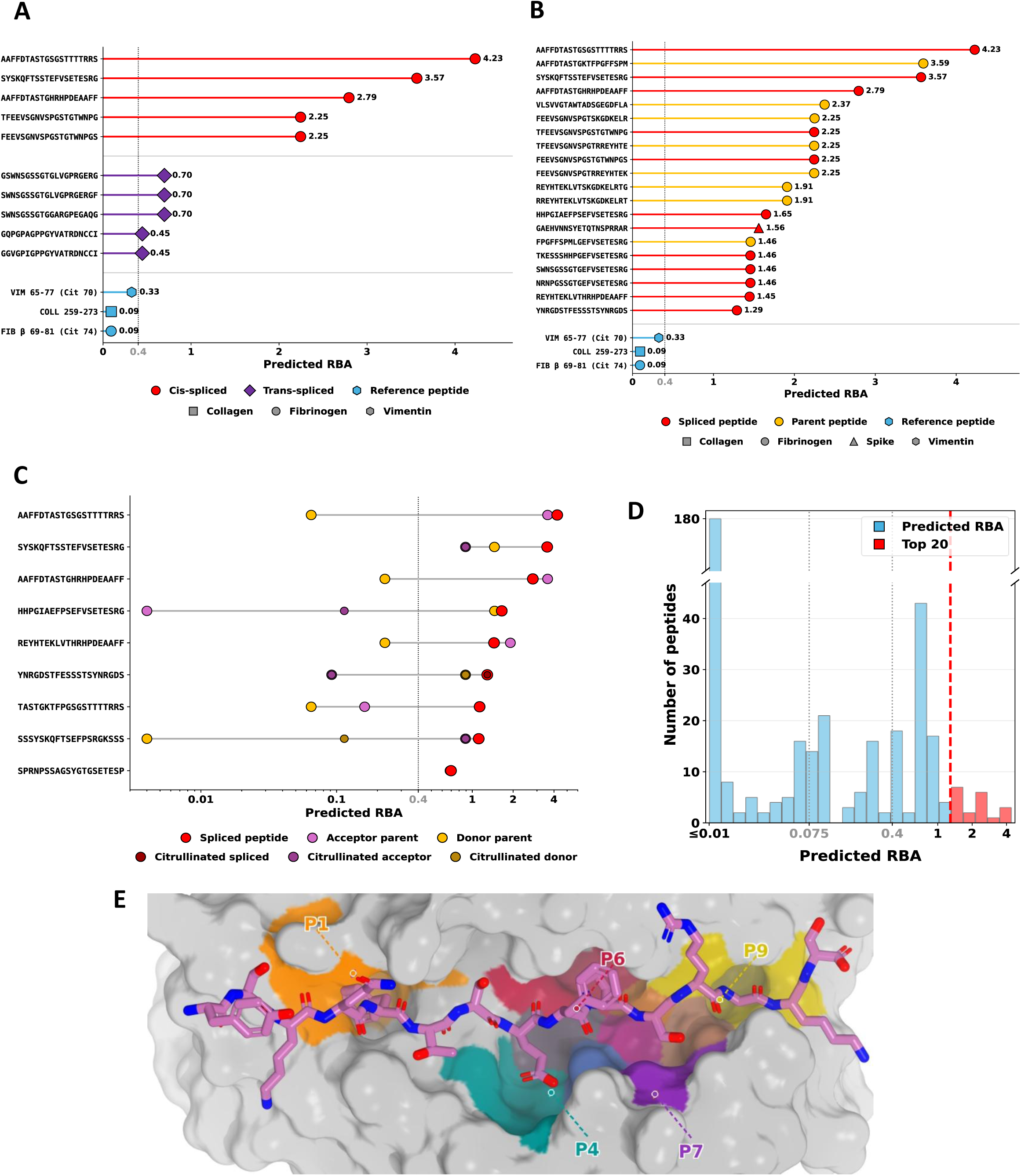
Predicted binding affinity of cis- and trans-spliced 20-mers to HLA-DR4 compared to known high-affinity binders and genomic 20-mers. **A)** Top 5 cis- and trans-spliced peptide sequences and **B)** top 20 peptide sequences (spliced and genomic) presenting the best predicted RBA to HLA-DRB1*04:01, compared to the predicted binding affinity of known immunological binders. **C)** Predicted affinity of top spliced peptide sequences to the receptor compared to their acceptor/donor genomic parent sequences and citrullinated versions of these sequences (if applicable). **D)** Distribution of RBA scores for all peptide sequences with the top 20 binders highlighted (red). **E)** Predicted binding conformation obtained for **YSKQFTSEFPSRGKS** in the HLA-DR4 binding groove, an enzymatically obtained cis-spliced 15-mers. Binding simulations were performed using Boltz-2, and binding affinity was calculated using PyRosetta. Positional coloring: P1 (orange), P4 (teal), P6 (crimson), P7 (purple), and P9 (gold).

Across the broader dataset, 12 spliced products (11 from fibrinogen, one from Spike) ranked within the top 20 predicted binders when assessed alongside acceptor/donor genomic 20-mer sequences (**Figure 6B-D**), and a total of 89 spliced peptides (81 cis-, 8 trans-) exceeded the RBA threshold of 0.075 below which peptides are not predicted to bind with measurable affinity (**Table S6**). In total, 190 out of 279 spliced peptides were not considered for further analysis because their predicted RBA was below a threshold value of 0.075 (**Table S6 & Figure 6D**). All three immunogenic controls passed this RBA threshold.

**Table 6.**
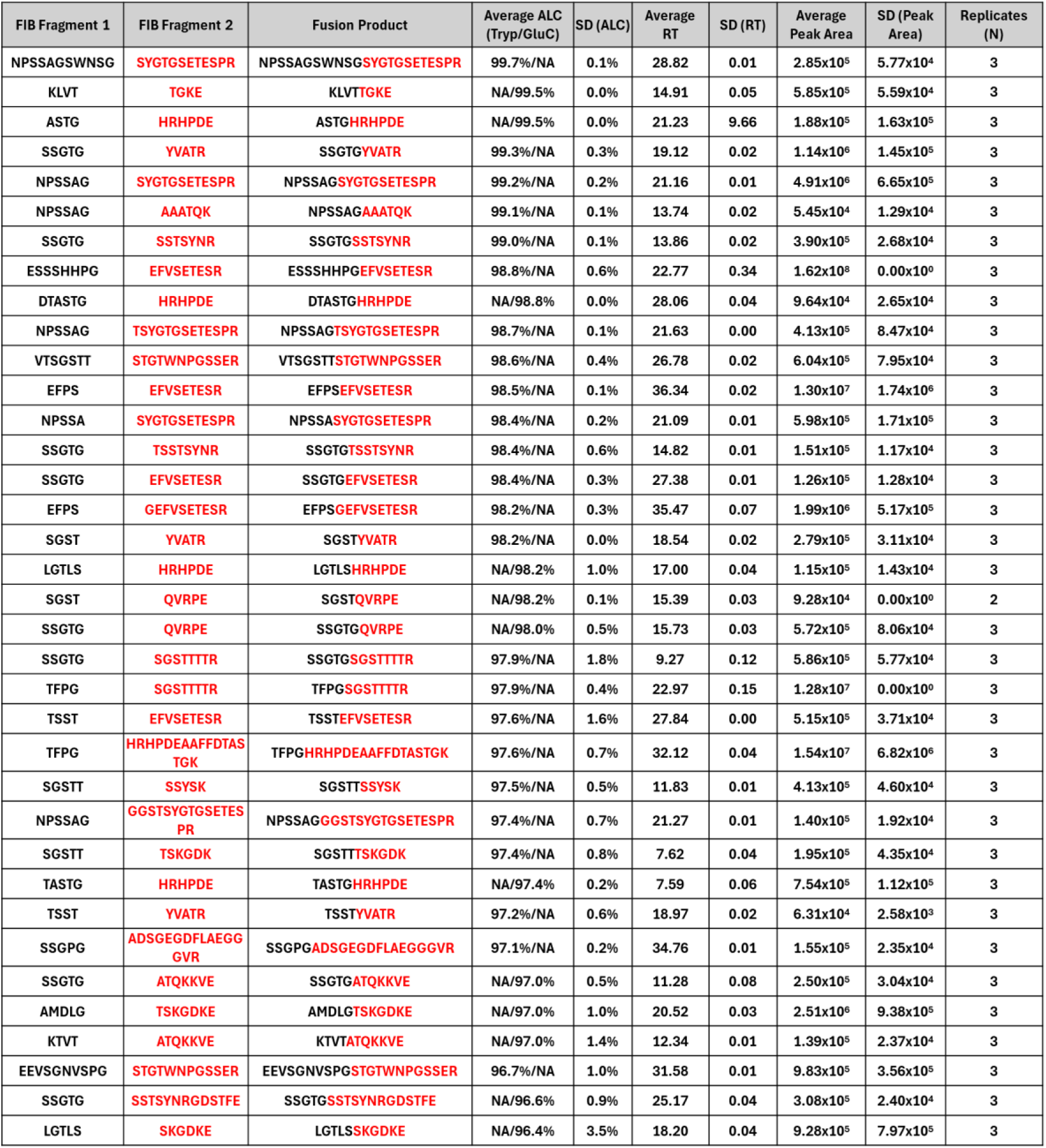
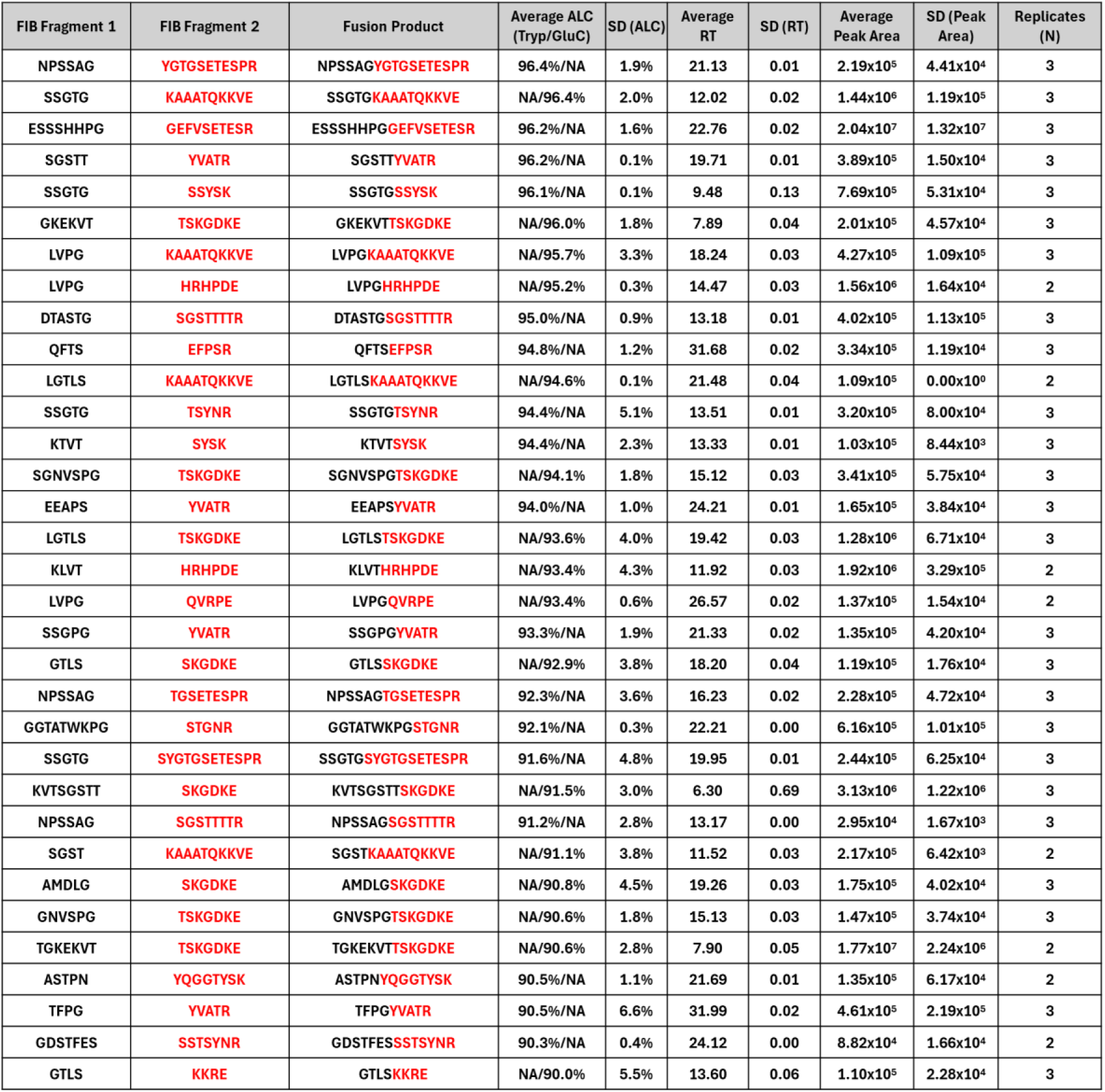
Cis-spliced peptides catalyzed by hCatK between fibrinogen fragments. Cis-spliced peptides were identified by database-assisted de novo peptide sequencing in PEAKS Studio 12.5, excluding all matches to reference proteomes, with ALC ≥ 90%, detected in at least 2 of 3 replicates, and absent from all E64-inhibited controls. Columns list: acceptor sequence (black), donor sequence (red), full cis-spliced peptide sequence, average ALC score across replicates for trypsin and GluC digestions (Tryp/GluC; NA indicates the peptide was not detected in that digestion condition, %), standard deviation of ALC (SD ALC, %), average retention time (RT, min), standard deviation of RT (SD RT, min), average peak area (arbitrary units), standard deviation of peak area (SD Peak Area, arbitrary units), and number of replicates in which the peptide was detected (N). Protein abbreviation: FIB = human fibrinogen. Note: An SD for peak area of 0.00 x 10^0^ a.u. indicates that this cis-spliced peptide was detected in ≥ 2 replicates but only quantified in one.

**Table 7.** Cis-spliced peptides catalyzed by hCatK between Spike fragments. Cis-spliced peptides were identified by database-assisted de novo peptide sequencing in PEAKS Studio 12.5, excluding all matches to reference proteomes, with ALC ≥ 90%, detected in at least 2 of 3 replicates, and absent from all E64- inhibited controls. Columns list: acceptor sequence (black), donor sequence (red), full cis-spliced peptide sequence, average ALC score across replicates for trypsin and GluC digestions (Tryp/GluC; NA indicates the peptide was not detected in that digestion condition, %), standard deviation of ALC (SD ALC, %), average retention time (RT, min), standard deviation of RT (SD RT, min), average peak area (arbitrary units), standard deviation of peak area (SD Peak Area, arbitrary units), and number of replicates in which the peptide was detected (N).

| Spike Fragment 1 | Spike Fragment 2 | Fusion Product | Average ALC (Tryp/GluC) | SD (ALC) | Average RT | SD (RT) | Average Peak Area | SD (Peak Area) | Replicates (N) |
| --- | --- | --- | --- | --- | --- | --- | --- | --- | --- |
| NSYE | TQTNSPR | NSYETQTNSPR | 98.6%/NA | 0.4% | 22.60 | 0.02 | 6.34x10<sup>5</sup> | 1.52x10<sup>5</sup> | 3 |
| MSECVLG | ASANLAATK | MSECVLGASANLAATK | 96.9%/NA | 0.4% | 35.43 | 0.01 | 1.34x10<sup>5</sup> | 2.73x10<sup>4</sup> | 3 |

**Table 8.** Trans-spliced peptides catalyzed by hCatK between DTIIC, fibrinogen, and Spike fragments. Trans-spliced peptides were identified by database-assisted de novo peptide sequencing in PEAKS Studio 12.5, excluding all matches to reference proteomes, detected in at least 2 of 3 replicates, and absent from all E64-inhibited controls. Columns list: acceptor sequence (black), donor sequence (red), full trans- spliced peptide sequence, average ALC score across replicates for trypsin and GluC digestions (Tryp/GluC; NA indicates the peptide was not detected in that digestion condition, %), standard deviation of ALC (SD ALC, %), average retention time (RT, min), standard deviation of RT (SD RT, min), average peak area (arbitrary units), standard deviation of peak area (SD Peak Area, arbitrary units), and number of replicates in which the peptide was detected (N). Protein abbreviations: DTIIC = denatured bovine type II collagen; FIB = human fibrinogen; S2 = Subunit 2 of the SARS-CoV-2 Spike protein. Note: An SD for peak area of 0.00 x 10^0^ a.u. indicates that this trans-spliced peptide was detected in ≥ 2 replicates but only quantified in one; * Trans-spliced peptides with ALC < 90%.

| Protein 1 Fragment | Protein 2 Fragment | Fusion Product | Average ALC (Tryp/GluC) | SD (ALC) | Average RT | SD (RT) | Average Peak Area | SD (Peak Area) | Replicates (N) |
| --- | --- | --- | --- | --- | --- | --- | --- | --- | --- |
| SGSTT (FIB) | LVGPR (DTIIC) | SGSTTLVGPR | 99.0%/NA | 0.0% | 24.60 | 0.02 | 7.45x10<sup>5</sup> | 1.33x10<sup>5</sup> | 3 |
| SSGTG (FIB) | LVGPR (DTIIC) | SSGTGLVGPR | 98.5%/NA | 0.4% | 23.85 | 0.02 | 5.12x10<sup>5</sup> | 9.78x10<sup>4</sup> | 3 |
| SSGTG (FIB) | GARGPE (DTIIC) | SSGTGGARGPE | NA/96.3% | 1.6% | 12.88 | 0.00 | 2.49x10<sup>5</sup> | 4.49x10<sup>4</sup> | 2 |
| GPPG (DTIIC) | KAAATQKKVE (FIB) | GPPGKAAATQKKVE | NA/95.5% | 3.3% | 12.83 | 0.01 | 1.64x10<sup>6</sup> | 2.06x10<sup>5</sup> | 3 |
| GHRPLD (FIB) | GAQGPR (DTIIC) | GHRPLDGAQGPR | 94.8%/NA | 2.8% | 16.97 | 0.05 | 3.57x10<sup>5</sup> | 3.40x10<sup>4</sup> | 3 |
| SGST (FIB) | LVGPR (DTIIC) | SGSTLVGPR | 93.8%/NA | 5.7% | 23.27 | 0.02 | 1.11x10<sup>5</sup> | 1.23x10<sup>4</sup> | 3 |
| GPPG (DTIIC) | YVATR (FIB) | GPPGYVATR | 91.7%/NA | 4.0% | 20.43 | 0.01 | 2.72x10<sup>6</sup> | 1.89x10<sup>5</sup> | 3 |
| KGAA (DTIIC) | MSECVLGQSK (S2) | KGAA MSECVLGQSK* | 84.8%/NA | 0.3% | 31.32 | 0.03 | 4.55x10<sup>5</sup> | 9.85x10<sup>3</sup> | 2 |
| ADAG (S2) | GLVGPR (DTIIC) | ADAGGLVGPR* | 80.7%/NA | 6.5% | 24.49 | 0.02 | 2.12x10<sup>5</sup> | 0.00x10<sup>0</sup> | 2 |

Comparative analyses further revealed that many spliced peptides displayed predicted binding affinities markedly exceeding those of the individual acceptor and donor genomic sequences from which they were derived (**Figure 6C**). Notably, a subset of these products also outperformed citrullinated variants of their parent genomic sequences in predicted HLA-DR4 affinity (**Figure 6C**). These results indicate that the act of splicing itself, as a post-translational modification, can substantially remodel the HLA-binding landscape of the resulting peptides. Spike-derived trans-spliced peptides did not show any predicted binding affinity differing from the parental genomic sequences (data not shown).

To solidify our predictive claims further, a comparative *in silico* structural binding prediction to the receptor encoded by HLA-DRB1*04:01 was performed using the open-source Boltz-2 software engine, incorporating multiple sequence alignments against phylogenetically related HLA class II alleles to maximize structural prediction accuracy (39). PyRosetta was used to measure predicted binding affinity by averaging the top 10 predicted peptide-HLA structures for each sequence (40). Previously confirmed spliced products were extended computationally to 15-mers (7 amino acids from the acceptor fragment, and 8 from the nucleophilic donor sequence), restricting this comparison to the confirmed spliced peptide pool given the substantially higher computational cost of structure-based co-folding relative to matrix scoring. Notably, seven of the top 20 predicted spliced binders by the matrix-based method were also within the top 20 predicted spliced binders by Boltz-2/PyRosetta-based prediction, and five of those seven ranked within the Boltz-2/PyRosetta top 10. Because sequence-derived motif scoring and structure-based co-folding rely on largely independent assumptions, this convergence at the top of an orthogonal ranking provides support that a subset of hCatK-generated spliced peptides are genuine high-affinity HLA-DR4 ligands rather than artifacts of a single predictive framework. Given that our matrix-based approach is calibrated against decades of experimentally determined HLA-DRB1*04:01 binding data, it was retained as the primary predictive framework throughout this study. One of those predicted spliced binders is presented in **Figure 6E**.

## DISCUSSION

### Biochemical and structural drivers of hCatK-mediated splicing

Our findings provide the first indication that cysteine protease-mediated splicing can generate RA-relevant neo-peptides with potentially strong binding affinity to the HLA- DRB1*04:01 receptor. This multilayered process is highly likely in endosomal and endolysosomal micro-environments where peptides are in the mM range, and where macromolecular crowding is the norm (16, 41). This study examined only hCatK, though multiple cathepsins act concurrently in physiological environments and would likely generate more complex product profiles.

The effect of pH on hCatK-mediated splicing has direct mechanistic and pathological implications. The shift in kinetic equilibrium favoring splicing events at mildly acidic to neutral pH (pH 6.5–7.5) aligns with the less protonated state of the donor peptide’s N-terminal α-amine, preserving its nucleophilic potency to attack the acyl-enzyme intermediate. Cathepsins, including hCatK, have been shown to be stable in these pH conditions (18, 30, 42). The increase in spliced product levels between pH values of 4.5 and 7.5 underlines that splicing could happen in various pH-dependent proteolytic environments. We have previously shown in a study using hCatS that its splicing activity reached 4.2% of the initial hydrolysis activity at an endosomal pH range closer to 6.5 (27). These rates are sufficient to reach high concentrations of spliced peptides which could competitively bind to disease-prone HLA receptors (43).

In a healthy system, the rapid acidification of the endolysosomal environment (pH∼4.5) would quickly suppress splicing in favor of proteolysis (44). However, the endolysosomal lumen of APCs frequently undergoes pathological alkalinization during chronic inflammation and cellular aging of joints (45, 46). These microenvironmental perturbations disrupt homeostatic proteostasis, creating a window that can favor efficient splicing. Alternatively, these splicing events may occur extracellularly within the neutral pH resorptive zones flanking osteoclasts and synovial macrophages, or within nascent, early endosomal vacuoles immediately following sub-synovial endocytosis. These extracellularly generated spliced peptides could replace MHC class II-bound antigenic peptides if they exhibit sufficient affinity and participate in T-cell epitope spreading (19, 47).

While splicing has been documented for other cysteine cathepsins such as hCatL and hCatS, this study also provides the first systematic biochemical and sequence-level characterization of hCatK-mediated peptide splicing on native, RA-relevant substrates under various physiological pH conditions (27, 48, 49). Spliced neo-peptides have profound pathological implications, as evidenced by the role of hybrid insulin peptides as super-agonistic autoantigens in type 1 diabetes; or the role of the proteasome in the generation of MHC class I-restricted spliced antigenic products (48, 50, 51). Our findings expand this paradigm by establishing that hCatK, a central player in synovial inflammation, joint degradation and antigen processing, shares this intrinsic splicing capacity on critical joint self-antigens like TIIC and fibrinogen, and RA-associated viral proteins such as the SARS-CoV-2 Spike protein (22, 52).

Our biochemical analysis highlights a key distinction between proteolysis and splicing: while hydrolysis by hCatK is mostly driven by substrate accessibility, transpeptidation has a far more conserved sequence filter at the target site. The preference for small amino acids (Gly, Thr, Ser) at the P1 target site indicates that low steric bulk within the enzyme’s S1 subsite is a prerequisite for splicing. This minimal steric footprint likely accelerates the reaction via two non-exclusive mechanisms. Firstly, it minimizes spatial constraints, thereby allowing the acyl-acceptor fragment to remain highly dynamic within the active site, increasing the probability of a productive alignment with the incoming nucleophile. Secondly, a small P1 side chain prevents crowding of the S1 pocket, opening a clear stereochemical trajectory for the incoming donor amine into the adjacent S1’ subsite. Furthermore, the enrichment of polar hydroxyl side chains (Thr, Ser) at P1 points to an additional chemical filter, where hydrogen bonding networks may stabilize the covalent acyl-enzyme intermediate to facilitate splicing.

In contrast to the stringent structural requirements at the P1 position, the nucleophilic donor sequence (positions N1-N4) lacks a rigid sequence conservation although it presents a certain preference for smaller residues. Some nucleophilic sequences also appear to be better at splicing than others. This suggests that local peptide abundance and regional spatial availability dominate the stochastic capture of the donor strand. This is further supported by the clustering of splicing junctions within the IDRs of full-length substrates, such as the αC-domain of fibrinogen. 80.8% of cis- and transpeptidation events in fibrinogen (42 out of the 52 donor/acceptor sites) have been identified in this region. Moreover, another 9.6% of cis- and transpeptidation sites were also found in the flexible N-terminal region of FGB (5 out of 52 donor/acceptor sites) harboring fibrinopeptide B. These sites account for all splicing events identified within the otherwise highly structured β-chain subunit. The enrichment of splicing within such IDRs suggests that structural flexibility and solvent exposure are key determinants of splicing competence by proteases, similar to proteolysis. Additionally, the participation of fibrinopeptides A and B as both donors and acceptors implies that these highly abundant, biologically active fragments, which accumulate at high systemic concentrations during inflammatory and thromboembolic events, serve as direct splicing sites.

Because these spliced products form autonomously from single protein species, autoimmune neo-epitopes could arise during routine intracellular antigen processing without requiring the co-localization or concurrent degradation of multiple distinct proteins. It is also of interest that viral components such as the SARS-CoV-2 Spike protein can be a source of spliced fragments and thus a potential source of neo-antigens.

### Methodological rigor of the workflow for spliced peptide identification

The higher ALC scores observed for confirmed spliced peptides relative to all peptides reflect the additional validation steps and thresholds used to confirm spliced peptides, including manual verification of randomly selected ones and exclusion of spliced peptides found in only one replicate. These findings validate the stringency of the identification workflow and support the high-confidence nature of the confirmed spliced peptides. Additionally, analysis of the distribution of spliced peptide candidates matched to spectra by *de novo* sequencing suggests that many additional spliced peptides could be generated by hCatK but remain undetected due to either insufficient chromatographic separation and ionization efficiency, or the inherent sequencing accuracy limits of *de novo* sequencing. While the stringent thresholds applied here limit the number of candidates considered for further validation, they were necessary to minimize type I errors and misidentifications. Therefore, the confirmed spliced peptides identified here likely represent a conservative lower bound of the total pool generated by hCatK activity.

The similarity in median peak areas calculated for all peptides with a mean ALC ≥ 90% and for confirmed spliced peptides suggests that these products are present at comparable signal intensities to canonical tryptic and Glu-C peptides. The smaller absolute number of spliced products relative to hydrolytic fragments indicates, however, that the transpeptidation reaction is selective rather than indiscriminate, targeting specific sequence contexts. This is consistent with the conserved P1 residue preferences described above.

### Immunological implications and autoimmune initiation

Considering the scale of spliced peptide formation, it is possible they could generate peptides capable of binding to RA-associated HLA receptors and acting as neoantigenic products with superagonist potential. This assumption is supported by our finding that 67 spliced peptides showed *in silico* binding affinity to HLA-DRB1*04:01 using matrix-based binding predictions. Moreover, under inflammatory conditions, where elevated endolysosomal pH simultaneously dampens proteolysis and accelerates transpeptidation, these stable spliced fragments could accumulate. This kinetic pivot could likely increase the density of neo-epitopes displayed via the MHC class II pathway.

The length distribution of the confirmed spliced peptides (13 to 22 amino acids-long) matches the established structural geometry required for presentation by MHC class II molecules. This falls within the range required for hCatK to generate hybrid structures with sufficient physical length and stability to be presented by HLA receptors following antigen processing. Crucially, the matrix-scored *in silico* binding affinities of many of these spliced 20-mers surpassed those of established immunodominant HLA-DR4 ligands, suggesting that enzymatic splicing alone, without citrullination or other post-translational modifications, is sufficient to confer exceptional MHC class II binding competence. The subset of spliced products exhibiting predicted RBA values at or over 0.4, corresponding to very good binding to the receptor, could be expected to outcompete canonical peptides for HLA-DR4 occupancy even when present at limiting concentrations, raising the possibility that these low-abundance enzymatic byproducts could exert disproportionate immunological influence within the antigen-presenting compartment.

### Comparison between *in silico* binding prediction approaches

Matrix-based and Boltz-2/PyRosetta-based predictions rely on fundamentally different underlying methodologies and should not be expected to agree in every respect, nor should residual differences between them be used to discredit *in silico* predictions generally. While matrix-based predictions rely on experimental datasets obtained over many decades for HLA-DRB1*04:01 and sequence motif alignment, biomolecular co-folding relies on structure-aware deep-learning models. Both methods present innate bias and should be carefully compared.

The position-specific scoring matrix for HLA-DRB1*04:01 was calculated using genomic peptide sequences and might exclude the set of spliced peptide sequences which could contain larger sequence diversity in the binding groove. However, HLA- DRB1*04:01 is also known to have very conserved specificity; especially at position P1, P4, P7, P9, which is not factored in by constraint-free biomolecular co-folding as performed here (53). Future studies will need to elucidate the best method to predict binding between peptides and HLA receptors that do not rely only on canonical genomic sequence datasets while preserving specificity intrinsic to receptors of the DRB1 haplotype and others.

Taken together, we regard the matrix-based and *in silico* molecular docking approaches as complementary rather than competing predictors of HLA-DR4 binding. The matrix-based method offers a well-calibrated, high-throughput screen anchored in decades of empirical peptide motif data, whereas structure-aware docking provides an independent assessment less reliant on canonical genomic training sequences. On our spliced peptide dataset, the two methods converged substantially at the top of their rankings. The agreement between the two methods appears to reflect genuine peptide-receptor complementarity.

### Limitations and future directions

Some limitations in this study warrant consideration, including the use of heat-denatured bovine collagen, which models the unfolded states native to intracellular processing but does not replicate hCatK dynamics on highly ordered, native extracellular matrix fibrils. Others include the high micro-molar donor peptide concentrations that were required to clear the detection limit of electrophoretic blotting assays with avidin-FITC, the limitations intrinsic to LC-MS/MS data acquisition, and the non-zero false-positive rate of *de novo* sequencing which remains an inherent challenge. We systematically neutralized this final limitation through strict multi-replicate matching, automated proteome exclusion filters, and tracking diagnostic biotin-derived fragment ions. The other limitations are also addressed in different ways through cross-validation with other substrates/endoproteases and usage of physiological amounts of proteins for full-length protein assays (1-3 µM).

The primary translational gap is the current lack of *in vivo* or *in cellulo* models directly capturing these events within a complex cellular environment. To bridge this gap, our next phase of research will apply high-resolution tandem mass spectrometry to primary synoviocyte and macrophage models of rheumatoid arthritis. This will establish the precise spatial boundaries of cathepsin-mediated splicing and determine whether these hybrid peptides are efficiently processed, loaded, and presented to drive the propagation of autoreactive T-cells. Tetramer-based T-cell proliferation assays using peripheral synovial tissue from RA patients will also enable us to determine the potency of selected spliced peptides to induce T-cell proliferation and activation.

## MATERIAL & METHODS

### Materials

Immunization-grade bovine type II collagen (TIIC) solution was obtained from Chondrex, Inc., WA, USA, Cat. No. 20021, Lot No. 250205. Human research-grade fibrinogen was obtained from Prolytix, VT, USA; SKU. HCI-0150R, Lot No. QQ-0919- 2MG. Recombinant SARS-CoV-2 Spike protein S1 and S2 subunits were obtained from RayBiotech Life Inc., GA, USA; Cat. No. 230-01101-250, Lot No. 06G1324L and Cat. No. 230-01103-250, Lot No. 02G1023L, respectively. All of these were used as protein substrates.

Avidin–fluorescein conjugate was purchased from Invitrogen/Thermo Fisher Scientific, OR, USA; Ref. No. A821, Lot No. 2652967 and Lot No. 2831364, and was employed for biotin detection. Mass spectrometry grade Trypsin-ultra™ was obtained from New England Biolabs, MA, USA; Cat. No. P8101S, Lot No. 10189739. Mass spectrometry-grade Glu-C Endoproteinase was obtained from Thermo Fisher Scientific, OR, USA; Cat. No. 90054, Lot No. AA394154.

The biotinylated peptides TSTCitGGK-Biotin, and ETNLGGK-Biotin and FRET-based substrate Ac-AE(EDANS)PGKAGEGGK-Dabcyl were all custom synthesized to >98% purity by Biomatik Co. (ON, Canada). The synthetic spliced peptides TGAVGGPAGE-Hyp-GRE, NPSSAGAAATQK, and GHRPLDGAQGPR were all custom synthesized to >98% purity by GenScript (NJ, USA). Hyp = Hydroxyproline.

Human cathepsin K was expressed in *Pichia pastoris* and purified as previously described (54). The enzyme was titrated with E64 prior to use as described in (55).

### Enzyme kinetics using the FRET substrate and the biotinylated donor peptide

The peptide Ac-AE(EDANS)PGKAGEGGK-Dabcyl was used as FRET substrate. Assays were performed with 500 nM hCatK at 25°C in four buffers with distinct pH: 0.1 M sodium acetate (pH 4.5), 0.1 M sodium acetate (pH 5.5), 0.1 M MES (pH 6.5), and 0.1 M HEPES (pH 7.5); all also contained 2.5 mM DTT and 2.5 mM EDTA. Substrate concentrations ranged from 0–60 µM. The assays were performed in the presence of 0, 20, or 200 µM of the peptide TSTCitGGK-Biotin. Fluorescence was monitored on a Synergy H1 microplate reader (BioTek; excitation 336 nm, emission 490 nm). Michaelis–Menten parameters (K_m_, *k*_cat_) were determined by nonlinear regression on GraphPad Prism. Assays were performed in triplicate, in three independent experiments. Relative quantification of reaction products was derived from extracted peak areas at each pH/timepoint.

### Enzymatic reactions using the FRET substrate and the biotinylated donor peptide for LC-MS/MS analysis

The workflow was adapted from (27). For time-course experiments, reaction mixtures contained 20 µM of the FRET substrate Ac-AE(EDANS)PGKAGEGGK-Dabcyl and 200 µM of the biotinylated donor peptide TSTCitGGK-Biotin. Assays were performed with 500 nM hCatK in four buffers with distinct pH: 0.1 M sodium acetate (pH 4.5), 0.1 M sodium acetate (pH 5.5), 0.1 M MES (pH 6.5), and 0.1 M HEPES (pH 7.5); all assays also contained 2.5 mM DTT and 2.5 mM EDTA. For controls, the enzyme was pre-incubated with a 10-fold molar excess of E64 for 5 min prior to substrate addition. Reactions were incubated at room temperature in the dark and stopped at 10 min, 30 min, 1 h, 2 h, and 5 h. They were then quenched with E64 (10-fold molar excess relative to enzyme). Human cathepsin K was precipitated by adding methanol (>80% v/v), stored at −20°C for 2 h, and centrifuged (15 min). Supernatants were collected, dried by SpeedVac, reconstituted in 0.5% TFA, and desalted using StageTips prior to LC-MS/MS analysis.

### Enzymatic reactions between protein substrates and the biotinylated donor peptide

Collagen was denatured by heating at 70°C for 15 min in 0.05 M acetic acid buffer solution prior to co-incubation with hCatK. Independent reaction mixtures contained denatured collagen (20 µg), fibrinogen (20 µg), or Spike subunits (5 µg of S1 or S2) combined with 1 mM of the biotinylated peptide TSTCitGGK-Biotin and diluted in <1% DMSO. Reactions were performed with 5 to 500 nM human cathepsin K (optimized for each substrate) in four buffers with distinct pH: 0.1 M sodium acetate (pH 4.5), 0.1 M sodium acetate (pH 5.5), 0.1 M sodium phosphate (pH 6.5), and 0.1 M HEPES (pH 7.5); all also contained 2.5 mM DTT and 2.5 mM EDTA, in triplicates. For controls, the enzyme was pre-incubated with a 10-fold molar excess of E64 for 5 min prior to substrate addition. Reactions were incubated at 28°C (if containing denatured collagen) or room temperature (if containing fibrinogen and Spike subunits) for 4 h in the dark and quenched with E64 (10-fold molar excess relative to enzyme). For LC-MS/MS analysis, proteins were then denatured by adding TFE (20% v/v) and heating at 95°C for 10 min, reduced with 10 mM TCEP and alkylated with 40 mM CAA for 5 min at 70°C, diluted in 50 mM ammonium bicarbonate, and digested with either trypsin or Glu-C endoproteinase overnight in a ratio of 1:50 (protease:protein) at 37°C. Samples were subsequently dried in a SpeedVac, reconstituted in 0.5% TFA, and desalted using StageTips prior to LC–MS/MS analysis.

### SDS-PAGE/Coomassie staining and avidin-FITC blotting assays

Reaction products were diluted in SDS sample buffer to the final concentration of 1.2% SDS, 1% glycerol, 0.001% bromophenol blue, 0.03 M Tris-HCl, pH 6.8, and 2.5% (v/v) 2-mercaptoethanol heated at 100°C for 5 min. Samples were resolved by SDS-PAGE (15-20% polyacrylamide gels; 20 µg protein per well). For Coomassie staining, the resolving gel was washed with dH_2_O, stained in Coomassie Blue staining solution (2.5 g Coomassie Brilliant Blue R-250, 450 mL methanol, 100 mL acetic acid per 1 L) for 1 h, and destained overnight in Coomassie Blue destaining solution (450 mL methanol, 100 mL acetic acid per 1 L). For avidin-FITC blotting assays, the resolving gel was used to transfer proteins onto nitrocellulose membranes (Amersham™ Protran™, GE Healthcare) using a Mini-PROTEAN Tetra Cell (Bio-Rad; 90 mA/gel, 120 min) in ice-cold transfer buffer (2.9 g glycine, 5.8 g Tris, 200 mL methanol per 1 L, pre-chilled at –20 °C for at least 2 h). Membranes were blocked overnight at 4°C in 1% (w/v) skimmed milk in TBST (50 mM Tris, 150 mM NaCl, 0.1% Tween-20, pH 7.5), washed in TBST (4 × 5 min) and PBS (2 × 5 min), and probed with avidin–FITC (25 µg/mL in PBS) for 1 h at room temperature with gentle rocking. Excess probe was removed by washing with PBS + 1% Tween-20 (2 × 5 min, 1 x 15 min, 2 x 5 min). Fluorescence was imaged on an Amersham Typhoon scanner (excitation 488 nm).

### Enzymatic reactions between full-length protein substrates for LC-MS/MS analysis

Enzymatic reactions with the full-length protein substrates were performed and analyzed by LC-MS/MS. Independent reaction mixtures contained denatured collagen (20 µg), fibrinogen (20 µg), or Spike subunits (5 µg each of S1 and S2) individually (reactions *in cis*), or in combination (reactions *in trans*). Reactions were performed with 10 to 500 nM hCatK (optimized for each substrate) in a pH 6.5 buffer containing 0.1 M sodium phosphate, 2.5 mM DTT and 2.5 mM EDTA, in triplicates. For controls, the enzyme was pre-incubated with a 10-fold molar excess of E64 for 5 min prior to substrate addition. Reactions were incubated at 28°C (if containing denatured collagen) or room temperature (if not containing denatured collagen) for 4 h in the dark and quenched with E64 (10-fold molar excess relative to enzyme). For LC-MS/MS analysis, proteins were then denatured by adding TFE (20% v/v) and heating at 95°C for 10 min, reduced with 10 mM TCEP and alkylated with 40 mM CAA for 5 min at 70°C, diluted in 50 mM ammonium bicarbonate, and digested with either trypsin or Glu-C endoproteinase overnight in a ratio of 1:50 (protease:protein) at 37°C. Samples were subsequently dried in a SpeedVac, reconstituted in 0.5% TFA, and desalted using StageTips prior to LC–MS/MS analysis.

### LC-MS/MS analysis (Orbitrap Exploris 480)

The method was adapted from (27). *Liquid Chromatography:* 50 ng of peptides were injected and separated on-line using an Easy-nLC 1200 (Thermo Fisher Scientific) with Aurora Series analytical column, (25 cm x 75 μm 1.6 μm C18; Ion Opticks, Parkville, Victoria, Australia). The analytical column was heated to 40°C using an integrated column oven (PRSO-V2, Sonation, Biberach, Germany). Buffer A consisted of 0.1% aqueous formic acid and 2% acetonitrile in water, and buffer B consisted of 0.1% aqueous formic acid and 80% acetonitrile in water. A standard 60 min gradient was run from 2% B to 20% B over 46 min, then to 32% B over 15 min, then to 50% B from 61 to 66 min, then to 95% B over 5 min, held at 95% B for 8 min, then dropped to 3% B over 2 min, held at 3% B for 6 min. The analysis was performed at 0.25 μL/min flow rate. The Easy-nLC thermostat temperature was set at 7°C. *MS/MS Acquisition Method:* The peptides were analyzed with an Orbitrap Exploris 480 mass spectrometer (Orbitrap ExplorisTM 480, Thermo Fisher Scientific). The Nanospray FlexTM ion source was operated at 1900 V spray voltage and ion transfer tubes were heated to 290°C. During analysis, the Orbitrap Exploris 480 was operated in a data-dependent (DDA) mode with 20 dependent scans. The MS and MS/MS spectra were collected in positive mode. Full MS resolution was set to 60,000 with a normalized automatic gain control (AGC) target of 100%, 50% RF lens and 20 ms maximum injection time. Scan range from m/z 375 Th to m/z 1200 Th, and charge states from 2 to 5 were included. Dynamic exclusion was enabled to exclude after 1 time for 5 s. For MS/MS scans, the resolution was set to 15,000, with normalized AGC target at 50%, normalized HCD collision energy at 28%, and isolation windows at m/z 2 Th.

### LC-MS/MS analysis (timsTOF Pro)

The method was adapted from (56). *Liquid Chromatography:* 50 ng of peptides were injected and separated on-line using a nanoElute 2 UHPLC system (Bruker Daltonics) with an Aurora Series Gen3 (CSI) analytical column, (25 cm x 75 μm 1.7 μm C18 120Å; Ion Opticks, Parkville, Victoria, Australia). The analytical column was heated to 50°C using a column toaster (Bruker Daltonics). Buffer A consisted of 0.1% aqueous formic acid and 0.5% acetonitrile in water, and buffer B consisted of 0.1% aqueous formic acid and 0.5% water in acetonitrile. A gradient was run from 2% B to 12% B over 7.5 min, then to 33% B over the next 7.5 min, then to 95% B over 0.5 min, held at 95% B for 7.72 min. The analysis was performed at 0.3 μL/min flow rate. The nanoElute thermostat temperature was set at 7°C. *MS/MS Acquisition Method:* The peptides were analyzed with a timsTOF Pro mass spectrometer (timsTOF Pro, Bruker Daltonics). The Captive Spray ion source was operated at 1600 V capillary voltage, with 3 L/min drying gas heated to 200°C. During analysis, the timsTOF Pro was operated in data-independent acquisition mode with parallel accumulation-serial fragmentation (DIA-PASEF), with equal TIMS ramp and accumulation times of 100 ms. The MS and MS/MS spectra were collected in positive mode. Scan range from m/z 100 Th to m/z 1700 Th, and ion mobility range (1/K0) from 0.7 to 1.3 V·s/cm², were included. For each TIMS cycle, 7 DIA-PASEF scans were used, with 25 total precursor windows spanning m/z 299.5 to 1200.5 Th and variable isolation widths of 31–61 Th (1 Th overlap between neighboring windows). Collision energy was ramped linearly as a function of mobility, from 20 eV at 1/k0 = 0.6 V·s/cm² to 59 eV at 1/k0 = 1.6 V·s/cm².

### Identification and quantification of small peptide products

Product identification and quantification for the TIIC1–V1 pH and time course experiments, acquired on the Bruker timsTOF Pro in DIA-PASEF mode, were performed in Skyline 24.1 (57). The peptide list source was built directly from expected peptide m/z values obtained from samples run in DDA mode on Orbitrap Exploris 480. Peak areas were extracted from MS1 precursor ion chromatograms within the DIA-PASEF spectra for each pH and time point condition, using a mass tolerance of 10 ppm and a retention time window of ± 2 min, and used to quantify relative abundance of spliced and cleavage peptide products across conditions.

### LC-MS/MS data processing in PEAKS Studio by *de novo* peptide sequencing

The LC-MS/MS data processing workflow is adapted from (27). Raw LC-MS/MS data were processed in PEAKS Studio 12.5 (Bioinformatics Solutions Inc.) (58). Sequences were matched using either database-assisted *de novo* peptide sequencing (for full-length substrate experiments) or regular *de novo* peptide sequencing (for V1 transpeptidation experiments). Following sequencing, peptides were selected for downstream processing under two criteria sets. For transpeptidation analysis with V1: ALC ≥ 80%, orthogonally verified by the presence of diagnostic biotin fragment ions (m/z 227.0849 and 310.1584), absent from all controls, and restricted to sequences ≥4 amino acids. For cis- and transpeptidation analysis between full-length substrates: ALC ≥ 90%, not matching any reference proteome, detected in ≥ 2 of 3 replicates, absent from all controls, and restricted to sequences ≥4 amino acids. The following sequence databases in FASTA format were used: bovine type II collagen α1 chain (Uniprot P02459); human fibrinogen α, β, and γ chains (Uniprot P02671, P02675, P02679); SARS-CoV-2 Spike glycoprotein (Uniprot P0DTC2); and reference proteomes for *H. sapiens* (UP000005640; source of human fibrinogen), *B. taurus* (UP000009136; source of bovine type II collagen), *E. coli* K12 (UP000000625; source of recombinant SARS-CoV-2 Spike glycoprotein), *K. pastoris* (UP000094565; source of recombinant human cathepsin K), *S. scrofa* (UP000008227; source of porcine trypsin), and *S. aureus* (UP000434412; source of bacterial Glu-C endoproteinase). Spectral validation of selected spliced peptides was performed by comparison to synthetic peptide standards analyzed independently by LC-MS/MS. Spectral similarity was quantified by cosine similarity, computed on intensity vectors aligned to a common m/z grid within a 0.02 Da tolerance, with unmatched peaks assigned zero intensity.

### Identification of cleavage and splicing events in protein substrates

For cleavage analysis, a custom Python script was written to identify cleavage sites present under human cathepsin K and trypsin/Glu-C activity in triplicates, but absent in any samples in the triplicate digested with trypsin or Glu-C alone. This analysis was performed, for each protein substrate, on the list of peptides previously obtained by database-assisted *de novo* peptide sequencing and matched to the canonical primary sequence of each protein (type II collagen, fibrinogen, and Spike subunits S1 and S2) using PEAKS Studio. For splicing analysis, a custom Python script was written to identify peptides made up of two distinct non-contiguous sequences of four amino acids or more from either the same protein substrate (cis-spliced peptides) or two distinct ones (trans-spliced peptides) detected in at least 2 of 3 replicates, and absent in any samples in the triplicate containing E64. This analysis was performed, for each experiment, on the list of peptides previously obtained by *de novo* peptide sequencing using PEAKS Studio. To ensure data integrity and minimize false-positive peptide assignments arising from algorithmic misidentification, 3 spliced peptide candidates were randomly selected for each experiment and underwent manual validation. Reanalysis was performed in FreeStyle^TM^ (Thermo Fisher Scientific) by extracting the monoisotopic peptide mass and inspecting the corresponding tandem mass spectra. Validation included extracted ion chromatograms (±5 ppm), and for tagged peptides, diagnostic fragment ions of biotinylated peptides were specifically monitored at m/z 227.0849 and 310.1584.

### Matrix-based *in silico* binding prediction to HLA-DRB1*04:01

The probability that peptide sequences would be bound and presented by HLA- DRB1*04:01 was evaluated based on a previously published prediction method (32, 33). Briefly, the most likely DR0401 motif for each theoretical peptide was deduced by considering each possible combination of nine consecutive residues within each candidate spliced 20-mer. The key binding residues within each 9-mer (positions 1, 4, 6, 7, and 9) were assigned a coefficient from a matrix of values (experimentally determined or inferred based on published data) that reflect the expected binding of every standard amino acid plus citrulline. The highest possible score was assigned as the predicted relative binding affinity score for each spliced 20-mer. Based on prior work, peptides with scores below a threshold value of 0.075 are unlikely to have measurable binding to DRB1*04:01. For spliced peptides with a score over 0.075, we evaluated extended versions of acceptor/donor parent genomic 20-mers, corresponding to the spliced peptide, to assign the most likely motif and a predicted RBA for each parental 20-mer.

### *In silico* molecular docking to HLA-DRB1*04:01

In silico structures of MHC:peptide complexes were generated using Boltz-2 (39). Boltz-2 is an advanced open-source AI model built on a transformer-based architecture similar to AlphaFold3. Multiple sequence alignments (MSAs) are used to help guide Boltz-2 and were pre-generated for the HLA alpha and beta chains using the mmseqs2 server. No MSAs were generated for the peptide fragment due to the minimal structural guidance MSAs provide peptides and the lack of evolutionary information for spliced peptides. The structure predictions were run using 10 recycling steps and 25 diffusion samples. From each prediction the top 10 ranked models selected by Boltz-2 were subsequently relaxed using Rosetta FastRelax from the PyRosetta software suite (59). The Energy score, dSASA and number of hydrogen bonds were subsequently calculated between the HLA complex and the peptide of interest using the InterfaceAnalyzerMover function in the PyRosetta software suite (60). These values were averaged from the top 10 ranked models. PyRosetta is a python-based interface of the Rosetta modeling suite. Rosetta, developed by Baker and coworkers, uses knowledge based and physics-inspired statistical potentials to approximate the complexes’ free energy. The total free energy score is based on a combination of parameters which include attractive, repulsive, solvation, electrostatic, and h-bond energies which are combined with probabilities of Ramachandran, and side chain angles for each amino acid residue (40, 61).

## Code/data availability statement

The mass spectrometry proteomics data have been deposited to the ProteomeXchange Consortium via the PRIDE (62) partner repository with the dataset identifier PXD083738. The custom Python architecture written for automated cleavage identification and peptide splicing extraction is hosted on GitHub at https://github.com/D4rk1705/catk-splicing-ra and will be made publicly available upon publication; access is available to editors and reviewers upon request during the review process.

## Acknowledgements

We thank Dr. Preety Panwar, Dipon Saha, and Parsa Shafiekhanii for assistance with data collection and analysis. We thank Jeanne Yuan, Renata Moravcova, Maor Arad, Dr. Armando Alcazar, and other members of the Proteomics Core Facility at UBC for their advice and technical contributions in LC-MS/MS-based proteomics. We also thank Dr. Christopher M. Overall and Dr. François Jean for their insightful discussions and helpful suggestions. Graphs and figures were made using the Matplotlib Python library. The directed graph in **Figure 3** was created with [Cytoscape](https://cytoscape.org/) (63). The sequence conservation plots in **Figure 3** were created with the Logomaker Python API (64). Small molecules/peptides were illustrated using [ChemDraw](https://revvitysignals.com/products/research/chemdraw). Mass spectrometry infrastructure used here was supported by the Canada Foundation for Innovation, the BC Knowledge Development Fund, and the UBC Life Sciences Institute. This work was supported by Canadian Institutes of Health Research (CIHR) grants (PJT-155979) and a CIHR Canada Graduate Research Scholarship.

