## Supplemental Figures for "Cathepsin K Mediates the Formation of Potential Rheumatoid Arthritis-Relevant Cis- and Trans-Spliced Peptides Compatible With HLA-DR4 Presentation"

**A**

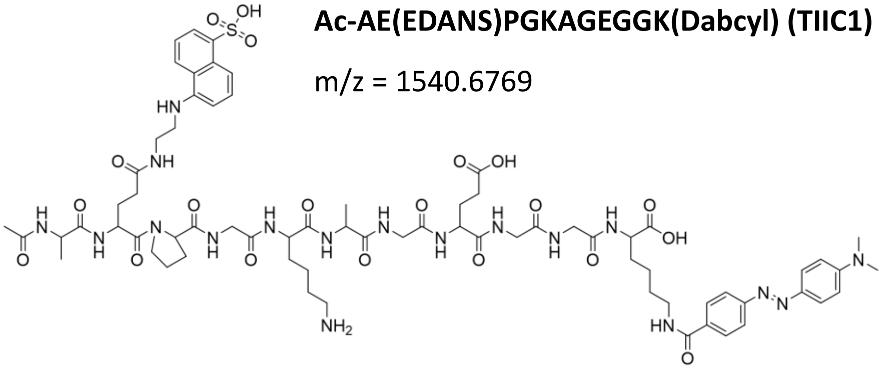

**Ac-AE(EDANS)PGKAGEGGK(Dabcyl) (TIIC1)**

m/z = 1540.6769

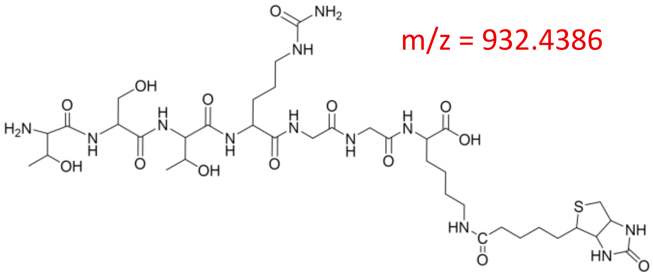

**TSTCitGGK-Biotin (V1)**

m/z = 932.4386

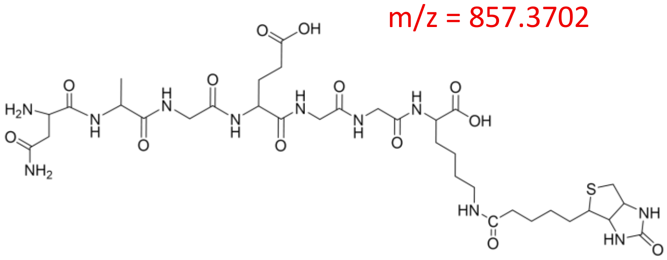

**NAGEGGK-Biotin (V2)**

m/z = 857.3702

**B**

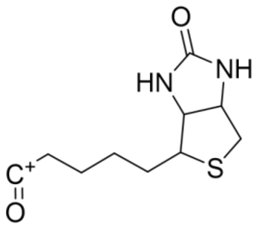

**Dehydrobiotin**

m/z = 227.0849

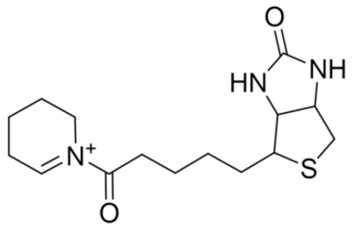

**ImKbiotin – $NH_3$**

m/z = 310.1584

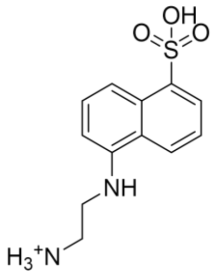

**EDANS**

m/z = 267.0798

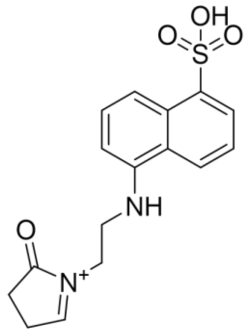

**ImEEDANS – $NH_3$**

m/z = 333.0904

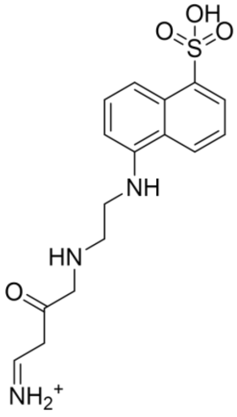

**ImEEDANS**

m/z = 350.1169

**Figure S1. Chemical structures and m/z values of A)** TIIC-derived (TIIC1) and vimentin-derived (V1, V2) peptides. **B)** signature ions observed following collision-induced dissociation (CID) of peptides/proteins labelled with Edans and Biotin.

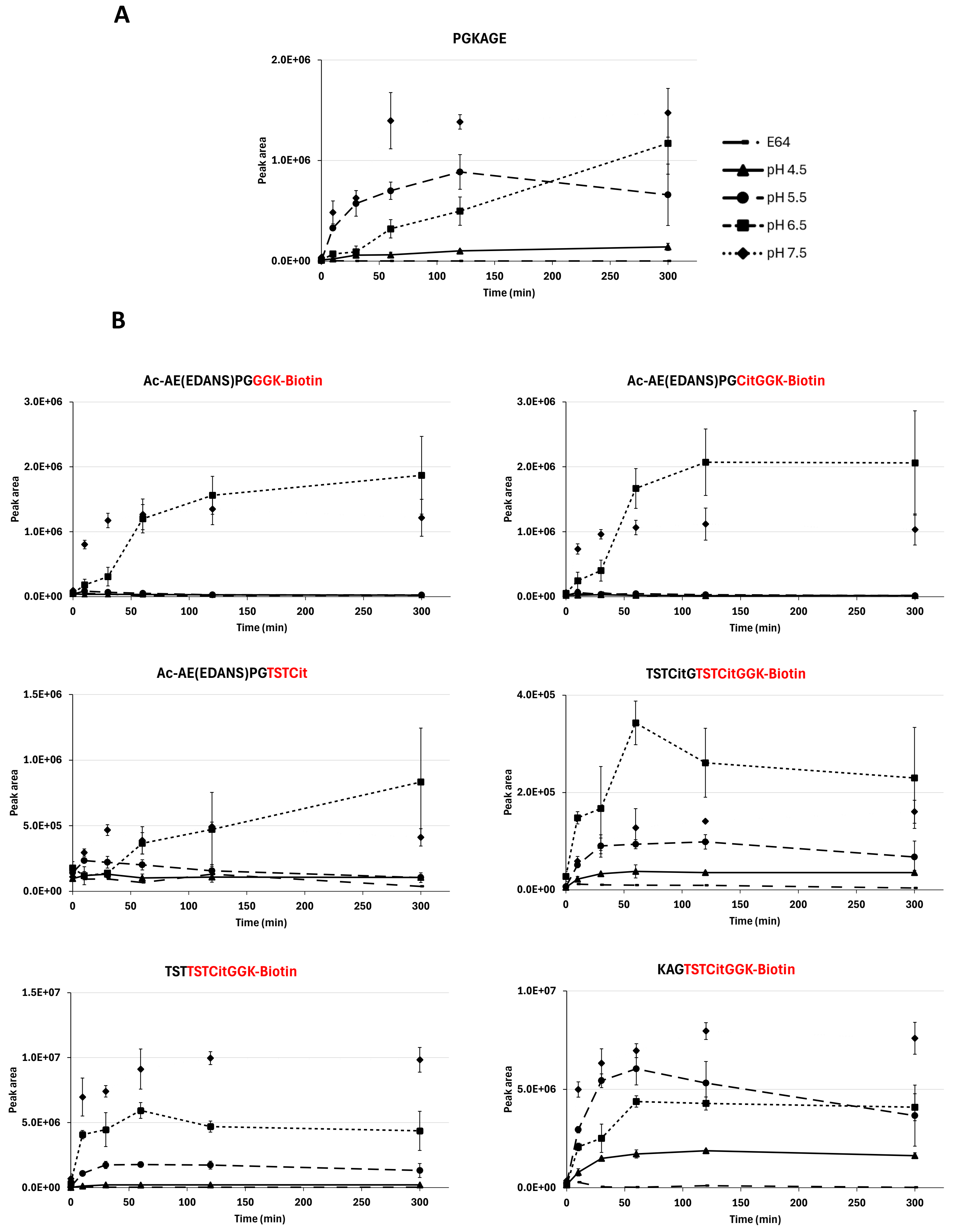


**Figure S2. Peak area over time for A)** other cleavage product(s) and **B)** other splicing products identified following digestion of TIIC1 and V1 by 500 nM hCatK, under four different pH conditions. Values represent three independent experiments; error bars indicate SD.

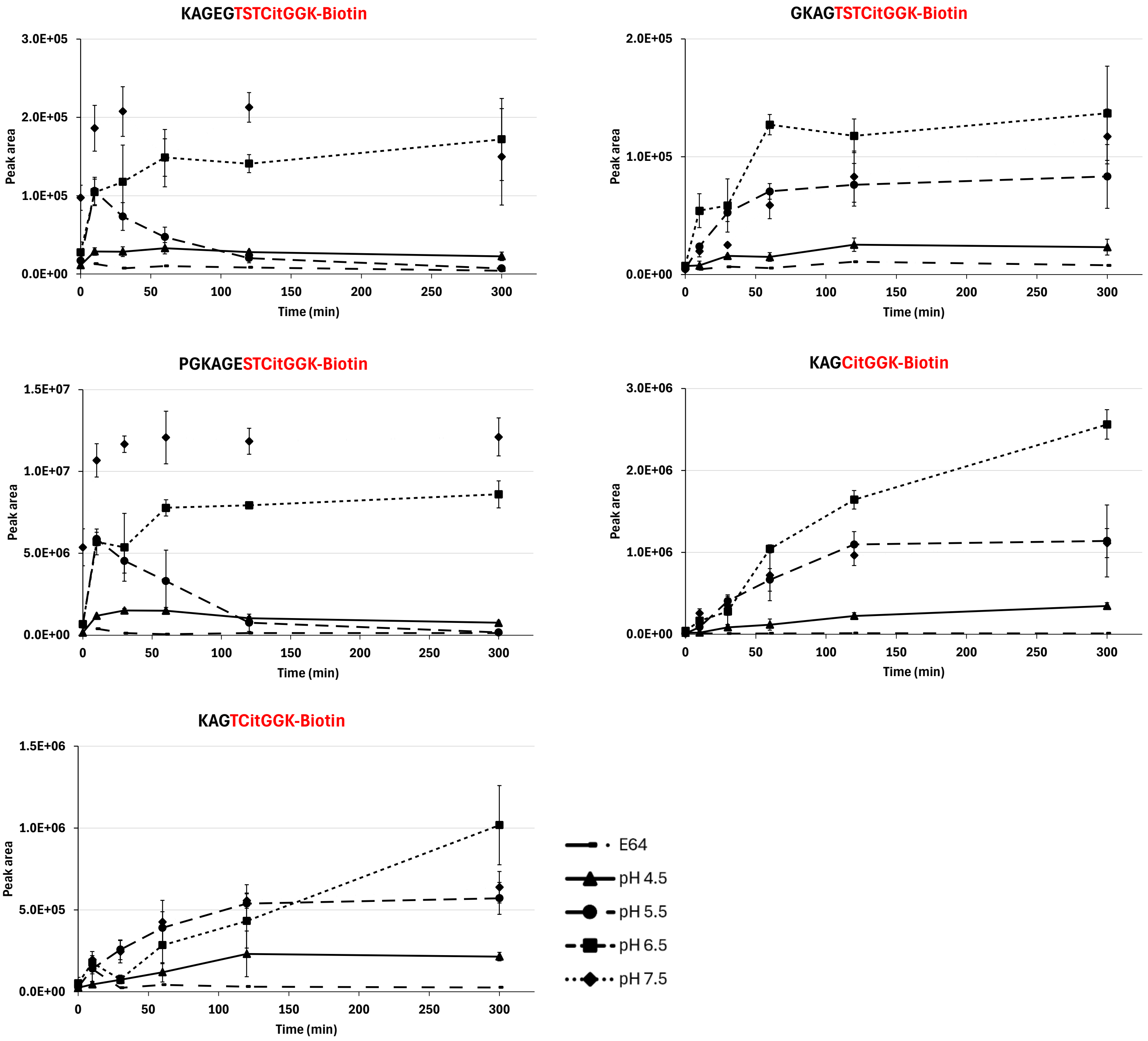


**Figure S2. (continued)**

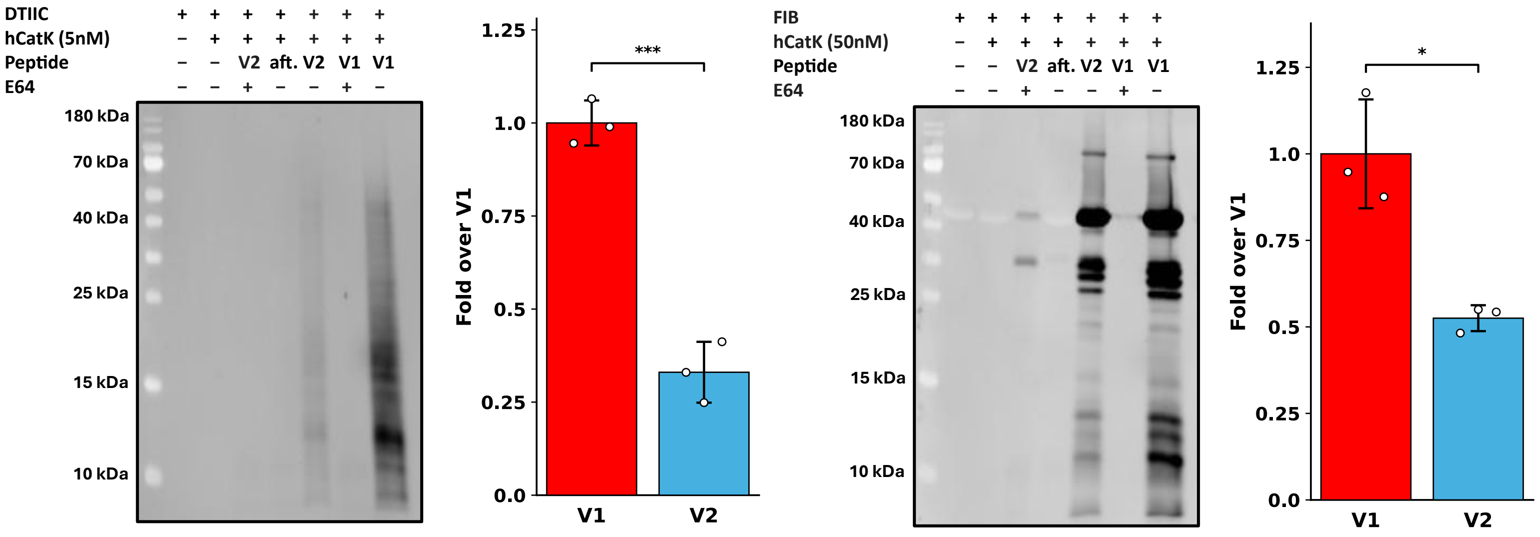


**Figure S3. Transpeptidation patterns observed following hCatK activity on RA-related substrates co-incubated with vimentin-derived peptides (V1 or V2).** Comparison and quantification of transpeptidation patterns observed for DTIIC or for fibrinogen co-incubated with either V1 or V2. Band intensities from avidin-FITC-stained membranes were quantified using ImageJ; aft. (after digestion). Values represent three independent experiments; error bars indicate SD.

Legend
Acceptor sequences
Acceptor site
**Disordered regions**

### Alpha 1 chain of bovine type II collagen

```
M I R L G A P Q T L V L L T L L V A A V L R C H G Q D V Q K A G S C V Q D G Q R Y N D K D V W K P E
P C R I C V C D T G T V L C D D I I C E D M K D C L S P E T P F G E C C P I C S A D L P T A S G Q P
G P K G Q K G E P G D I K D I V G P K G P P G P Q G P A G E Q G P R G D R G D K G E K G A P G P R G
R D G E P G T P G N P G P P G P P G P P G P P G L G G N F A A Q M A G G F D E K A G G A Q M G V M Q
G P M G P M G P R G P P G P A G A P G P Q G F Q G N P G E P G E P G V S G P M G P R G P P G P P G K
P G D D G E A G K P G K S G E R G P P G P Q G A R G F P G T P G L P G V K G H R G Y P G L D G A K G
E A G A P G V K G E S G S P G E N G S P G P M G P R G L P G E R G R T G P A G A A G A R G N D G Q P
G P A G P P G P V G P A G G P G F P G A P G A K G E A G P T G A R G P E G A Q G P R G E P G T P G S
P G P A G A A G N P G T D G I P G A K G S A G A P G I A G A P G F P G P R G P P G P Q G A T G P L G
P K G Q T G E P G I A G F K G E Q G P K G E P G P A G P Q G A P G P A G E E G K R G A R G E P G G A
G P A G P P G E R G A P G N R G F P G Q D G L A G P K G A P G E R G P S G L A G P K G A N G D P G R
P G E P G L P G A R G L T G R P G D A G P Q G K V G P S G A P G E D G R P G P P G P Q G A R G Q P G
V M G F P G P K G A N G E P G K A G E K G L P G A P G L R G L P G K D G E T G A A G P P G P A G P A
G E R G E Q G A P G P S G F Q G L P G P P G P P G E G G K P G D Q G V P G E A G A P G L V G P R G E
R G F P G E R G S P G S Q G L Q G A R G L P G T P G T D G P K G A A G P A G P P G A Q G P P G L Q G
M P G E R G A A G I A G P K G D R G D V G E K G P E G A P G K D G G R G L T G P I G P P G P A G A N
G E K G E V G P P G P A G T A G A R G A P G E R G E T G P P G P A G F A G P P G A D G Q P G A K G E
Q G E A G Q K G D A G A P G P Q G P S G A P G P Q G P T G V T G P K G A R G A Q G P P G A T G F P G
A A G R V G P P G S N G N P G P P G P P G P S G K D G P K G A R G D S G P P G R A G D P G L Q G P A
G P P G E K G E P G D D G P S G P D G P P G P Q G L A G Q R G I V G L P G Q R G E R G F P G L P G P
S G E P G K Q G A P G A S G D R G P P G P V G P P G L T G P A G E P G R E G S P G A D G P P G R D G
A A G V K G D R G E T G A V G A P G A P G P P G S P G P A G P I G K Q G D R G E A G A Q G P M G P A
G P A G A R G M P G P Q G P R G D K G E T G E A G E R G L K G H R G F T G L Q G L P G P P G P S G D
Q G A S G P A G P S G P R G P P G P V G P S G K D G A N G I P G P I G P P G P R G R S G E T G P A G
P P G N P G P P G P P G P P G P G I D M S A F A G L G Q R E K G P D P L Q Y M R A D E A A G N L R Q
H D A E V D A T L K S L N N Q I E S L R S P E G S R K N P A R T C R D L K L C H P E W K S G D Y W I
D P N Q G C T L D A M K V F C N M E T G E T C V Y P N P A S V P K K N W W S S K S K D K K H I W F G
E T I N G G F H F S Y G D D N L A P N T A N V Q M T F L R L L S T E G S Q N I T Y H C K N S I A Y L
D E A A G N L K K A L L I Q G S N D V E I R A E G N S R F T Y T V L K D G C T K H T G K W G K T M I
E Y R S Q K T S R L P I I D I A P M D I G G P E Q E F G V D I G P V C F L
```

**Figure S4. Transpeptidation sites identified in denatured type II collagen (DTIIC) after hCatK activity, with V1.** Sequences highlighted in purple are acceptor sequences identified in confirmed trans-spliced peptides with V1, and purple arrows indicate the identified transpeptidation sites.

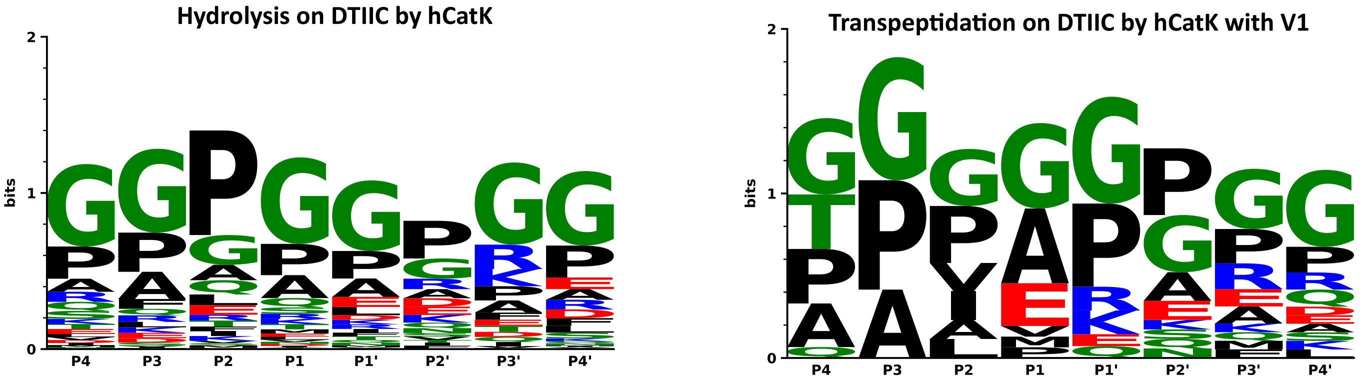


**Figure S5. Comparison between hydrolysis & transpeptidation patterns with V1 in DTIIC following hCatK activity.** Hydrolysis and transpeptidation happen at the scissile bond between the P1 and P1’ positions. Weblogos depict sequence conservation at each position for the two types of enzymatic events on DTIIC. For complete trans-spliced peptide identities, see Tables 2 and S3.

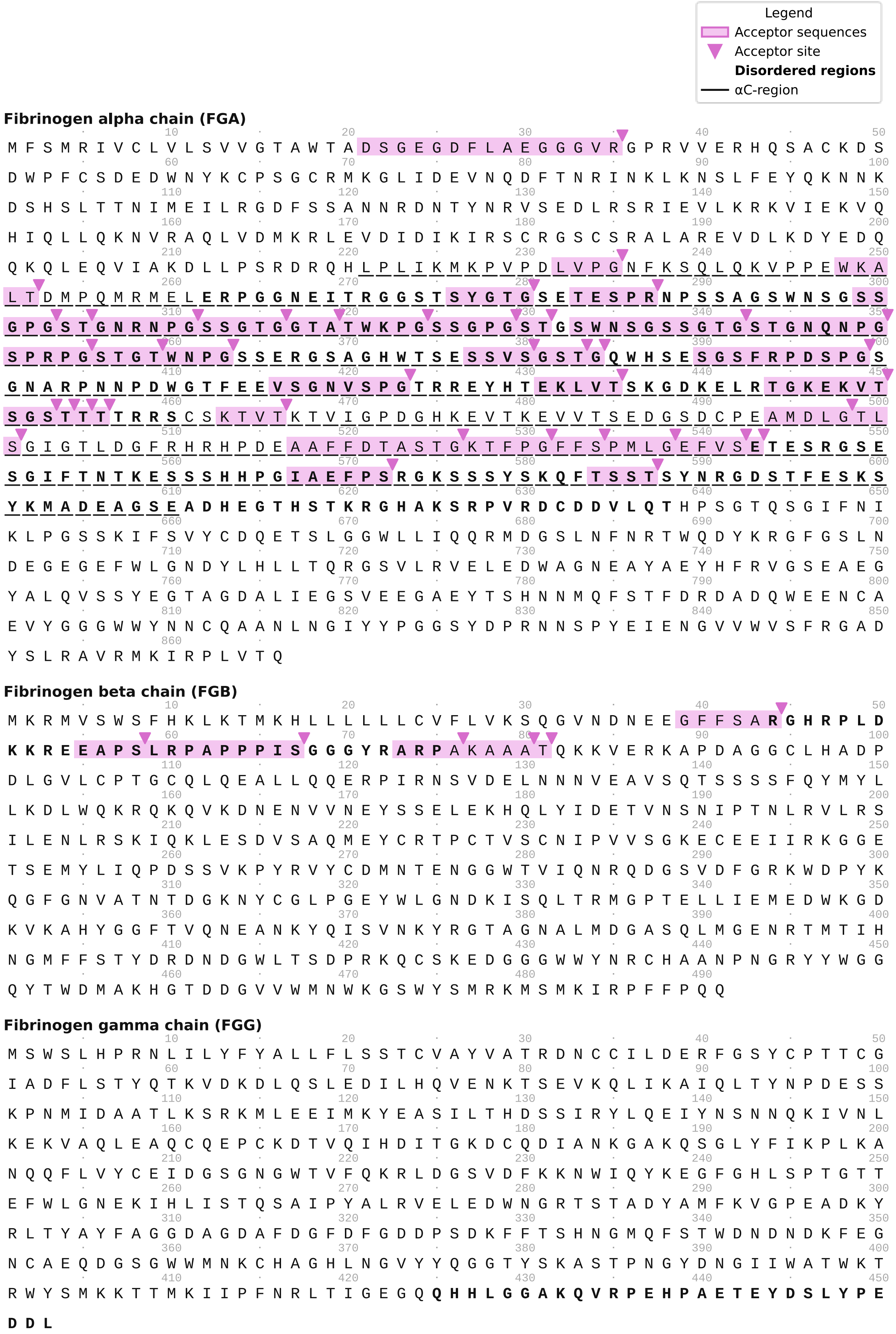


**Figure S6. Transpeptidation sites identified in fibrinogen after hCatK activity, with V1.** Bolded sequences indicate disordered regions in the primary protein sequence, according to region annotation in Uniprot of human fibrinogen α, β, and γ chains (Uniprot P02671, P02675, P02679). The underlined sequence represents the αC-region. Sequences highlighted in purple are acceptor sequences identified in confirmed trans-spliced peptides with V1, and purple arrows indicate the identified transpeptidation sites.

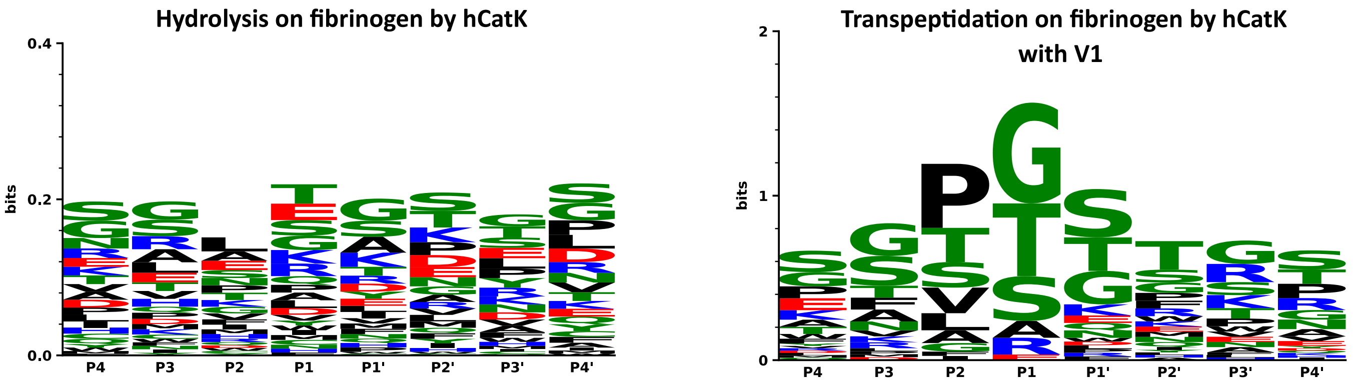


**Figure S7. Comparison between hydrolysis & transpeptidation patterns with V1 in fibrinogen following hCatK activity.** Hydrolysis and transpeptidation happen at the scissile bond between the P1 and P1' positions. Weblogos depict sequence conservation at each position for the two types of enzymatic events on fibrinogen. For complete trans-spliced peptide identities, see Tables 3 and S3.

Legend
Acceptor sequences
Acceptor site
**Disordered regions**

**S1 Protein (SARS-CoV-2)**

Q C V N L T T R T Q L P P A Y T N S F T R G V Y Y P D K V F R S S V L H S T Q D L F L P F F S N V T

W F H A I H V S G T N G T K R F D N P V L P F N D G V Y F A S T E K S N I I R G W I F G T T L D S K

T Q S L L I V N N A T N V V I K V C E F Q F C N D P F L G V Y Y H K N N K S W M E S E F R V Y S S A

N N C T F E Y V S Q P F L M D L E G K Q G N F K N L R E F V F K N I D G Y F K I Y S K H T P I N L V

R D L P Q G F S A L E P L V D L P I G I N I T R F Q T L L A L H R S Y L T P G D S S S G W T A G A A

A Y Y V G Y L Q P R T F L L K Y N E N G T I T D A V D C A L D P L S E T K C T L K S F T V E K G I Y

Q T S N F R V Q P T E S I V R F P N I T N L C P F G E V F N A T R F A S V Y A W N R K R I S N C V A

D Y S V L Y N S A S F S T F K C Y G V S P T K L N D L C F T N V Y A D S F V I R G D E V R Q I A P G

Q T G K I A D Y N Y K L P D D F T G C V I A W N S N N L D S K V G G N Y N Y L Y R L F R K S N L K P

F E R D I S T E I Y Q A G S T P C N G V E G F N C Y F P L Q S Y G F Q P T N G V G Y Q P Y R V V V L

S F E L L H A P A T V C G P K K S T N L V K N K C V N F N F N G L T G T G V L T E S N K K F L P F Q

Q F G R D I A D T T D A V R D P Q T L E I L D I T P C S F G G V S V I T P G T N T S N Q V A V L Y Q

D V N C T E V P V A I H A D Q L T P T W R V Y S T G S N V F Q T R A G C L I G A E H V N N S Y E C D

I P I G A G I C A S Y Q T Q T N S P R R A R

**S2 Protein (SARS-CoV-2)**

S V A S Q S I I A Y T M S L G A E N S V A Y S N N S I A I P T N F T I S V T T E I L P V S M T K T S

V D C T M Y I C G D S T E C S N L L L Q Y G S F C T Q L N R A L T G I A V E Q D K N T Q E V F A Q V

K Q I Y K T P P I K D F G G F N F S Q I L P D P S K P S K R S F I E D L L F N K V T L A D A G F I K

Q Y G D C L G D I A A R D L I C A Q K F N G L T V L P P L L T D E M I A Q Y T S A L L A G T I T S G

W T F G A G A A L Q I P F A M Q M A Y R F N G I G V T Q N V L Y E N Q K L I A N Q F N S A I G K I Q

D S L S S T A S A L G K L Q D V V N Q N A Q A L N T L V K Q L S S N F G A I S S V L N D I L S R L D

K V E A E V Q I D R L I T G R L Q S L Q T Y V T Q Q L I R A A E I R A S A N L A A T K M S E C V L G

Q S K R V D F C G K G Y H L M S F P Q S A P H G V V F L H V T Y V P A Q E K N F T T A P A I C H D G

K A H F P R E G V F V S N G T H W F V T Q R N F Y E P Q I I T T D N T F V S G N C D V V I G I V N N

T V Y D P L Q P E L D S F K E E L D K Y F K N H T S P D V D L G D I S G I N A S V V N I Q K E I D R

L N E V A K N L N E S L I D L Q E L G K Y E Q Y I K W P W Y I W L G F I A G L I A I V M V T I M L C

C M T S C C S C L K G C C S C G S C C K F D E D D S E P V L K G V K L H Y T

**Figure S8. Transpeptidation sites identified in Spike after hCatK activity, with V1.** Bolded sequences indicate disordered regions in the primary protein sequence, according to region annotation in Uniprot of SARS-CoV-2 Spike glycoprotein (Uniprot P0DTC2). Sequences highlighted in purple are acceptor sequences identified in confirmed trans-spliced peptides with V1, and purple arrows indicate the identified transpeptidation sites.

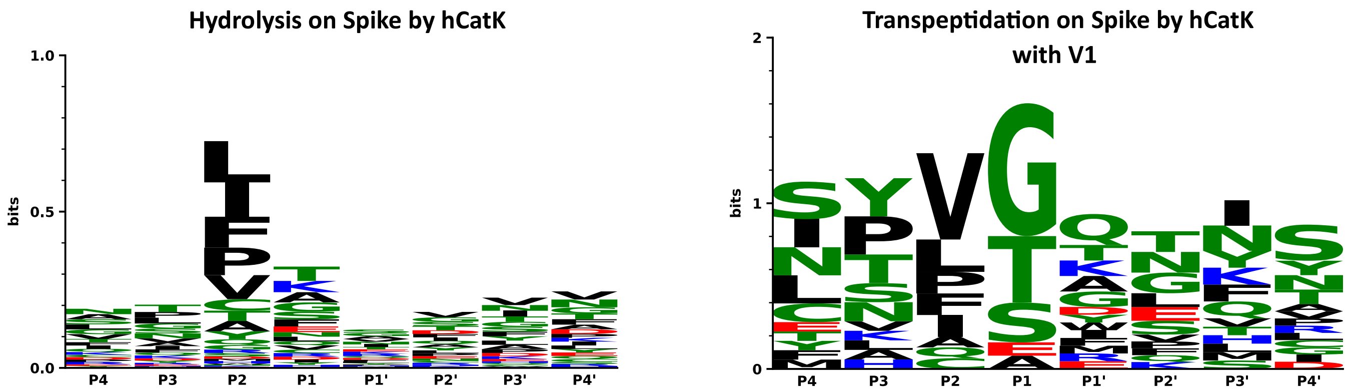


**Figure S9. Comparison between hydrolysis & transpeptidation patterns with V1 in Spike following hCatK activity.** Hydrolysis and transpeptidation happen at the scissile bond between the P1 and P1' positions. Weblogos depict sequence conservation at each position for the two types of enzymatic events on Spike. For complete trans-spliced peptide identities, see Tables 4 and S3.

Legend

- Acceptor sequences
- Acceptor site
- Donor sequences
- Donor site
- **Disordered regions**

**Alpha 1 chain of bovine type II collagen**

M I R L G A P Q T L V L L T L L V A A V L R C H G Q D V Q K A G S C V Q D G Q R Y N D K D V W K P E
P C R I C V C D T G T V L C D D I I C E D M K D C L S P E T P F G E C C P I C S A D L P T A S G Q P
G P K G Q K G E P G D I K D I V G P K G P P G P Q G P A G E Q G P R G D R G D K G E K G A P G P R G
R D G E P G T P G N P G P P G P P G P P G P P G L G G N F A A Q M A G G F D E K A G G A Q M G V M Q
G P M G P M G P R G P P G P A G A P G P Q G F Q G N P G E P G E P G V S G P M G P R G P P G P P G K
P G D D G E A G K P G K S G E R G P P G P Q G A R G F P G T P G L P G V K G H R G Y P G L D G A K G
E A G A P G V K G E S G S P G E N G S P G P M G P R G L P G E R G R T G P A G A A G A R G N D G Q P
G P A G P P G P V G P A G G P G F P G A P G A K G E A G P T G A R G P E G A Q G P R G E P G T P G S
P G P A G A A G N P G T D G I P G A K G S A G A P G I A G A P G F P G P R G P P G P Q G A T G P L G
P K G Q T G E P G I A G F K G E Q G P K G E P G P A G P Q G A P G P A G E E G K R G A R G E P G G A
G P A G P P G E R G A P G N R G F P G Q D G L A G P K G A P G E R G P S G L A G P K G A N G D P G R
P G E P G L P G A R G L T G R P G D A G P Q G K V G P S G A P G E D G R P G P P G P Q G A R G Q P G
V M G F P G P K G A N G E P G K A G E K G L P G A P G L R G L P G K D G E T G A A G P P G P A G P A
G E R G E Q G A P G P S G F Q G L P G P P G P P G E G G K P G D Q G V P G E A G A P G L V G P R G E
R G F P G E R G S P G S Q G L Q G A R G L P G T P G T D G P K G A A G P A G P P G A Q G P P G L Q G
M P G E R G A A G I A G P K G D R G D V G E K G P E G A P G K D G G R G L T G P I G P P G P A G A N
G E K G E V G P P G P A G T A G A R G A P G E R G E T G P P G P A G F A G P P G A D G Q P G A K G E
Q G E A G Q K G D A G A P G P Q G P S G A P G P Q G P T G V T G P K G A R G A Q G P P G A T G F P G
A A G R V G P P G S N G N P G P P G P P G P S G K D G P K G A R G D S G P P G R A G D P G L Q G P A
G P P G E K G E P G D D G P S G P D G P P G P Q G L A G Q R G I V G L P G Q R G E R G F P G L P G P
S G E P G K Q G A P G A S G D R G P P G P V G P P G L T G P A G E P G R E G S P G A D G P P G R D G
A A G V K G D R G E T G A V G A P G A P G P P G S P G P A G P I G K Q G D R G E A G A Q G P M G P A
G P A G A R G M P G P Q G P R G D K G E T G E A G E R G L K G H R G F T G L Q G L P G P P G P S G D
Q G A S G P A G P S G P R G P P G P V G P S G K D G A N G I P G P I G P P G P R G R S G E T G P A G
P P G N P G P P G P P G P P G P G I D M S A F A G L G Q R E K G P D P L Q Y M R A D E A A G N L R Q
H D A E V D A T L K S L N N Q I E S L R S P E G S R K N P A R T C R D L K L C H P E W K S G D Y W I
D P N Q G C T L D A M K V F C N M E T G E T C V Y P N P A S V P K K N W W S S K S K D K K H I W F G
E T I N G G F H F S Y G D D N L A P N T A N V Q M T F L R L L S T E G S Q N I T Y H C K N S I A Y L
D E A A G N L K K A L L I Q G S N D V E I R A E G N S R F T Y T V L K D G C T K H T G K W G K T M I
E Y R S Q K T S R L P I I D I A P M D I G G P E Q E F G V D I G P V C F L

**Figure S10. Cispeptidation sites identified in denatured type II collagen (DTIIC) after hCatK activity.** Sequences highlighted in purple and yellow are acceptor and donor sequences, respectively, identified in confirmed cis-spliced peptides; and purple and yellow arrows indicate the identified cispeptidation acceptor and donor sites, respectively.

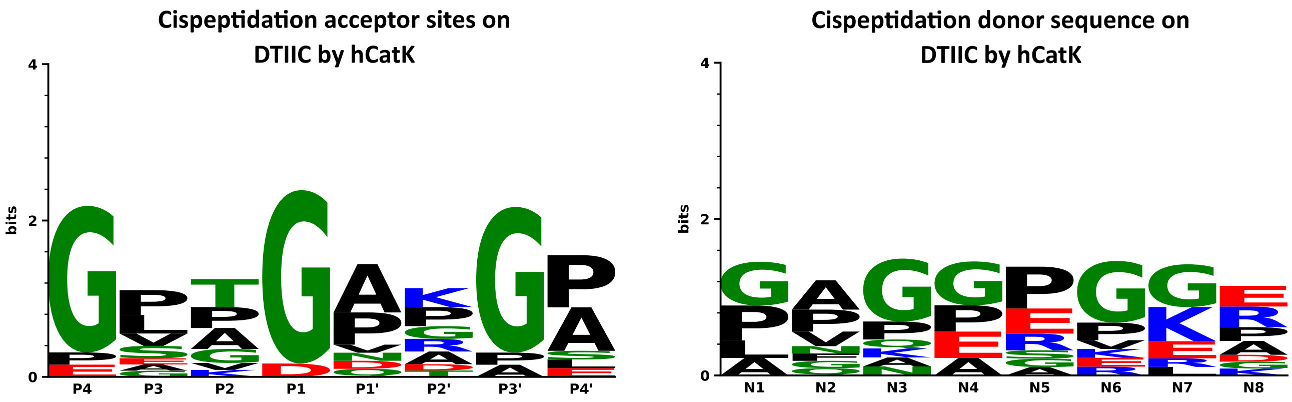


**Figure S11. Acceptor and donor sequences in DTIIC involved in cispeptidation by hCatK.** Cispeptidation happens at the scissile bond between the P1 and P1' positions and uses the primary amine at the N-terminal end of N1 as the nucleophilic donor. Weblogos depict sequence conservation at each position for both types of sequences on DTIIC. For complete cis-spliced peptide identities, see Tables 5 and S3.

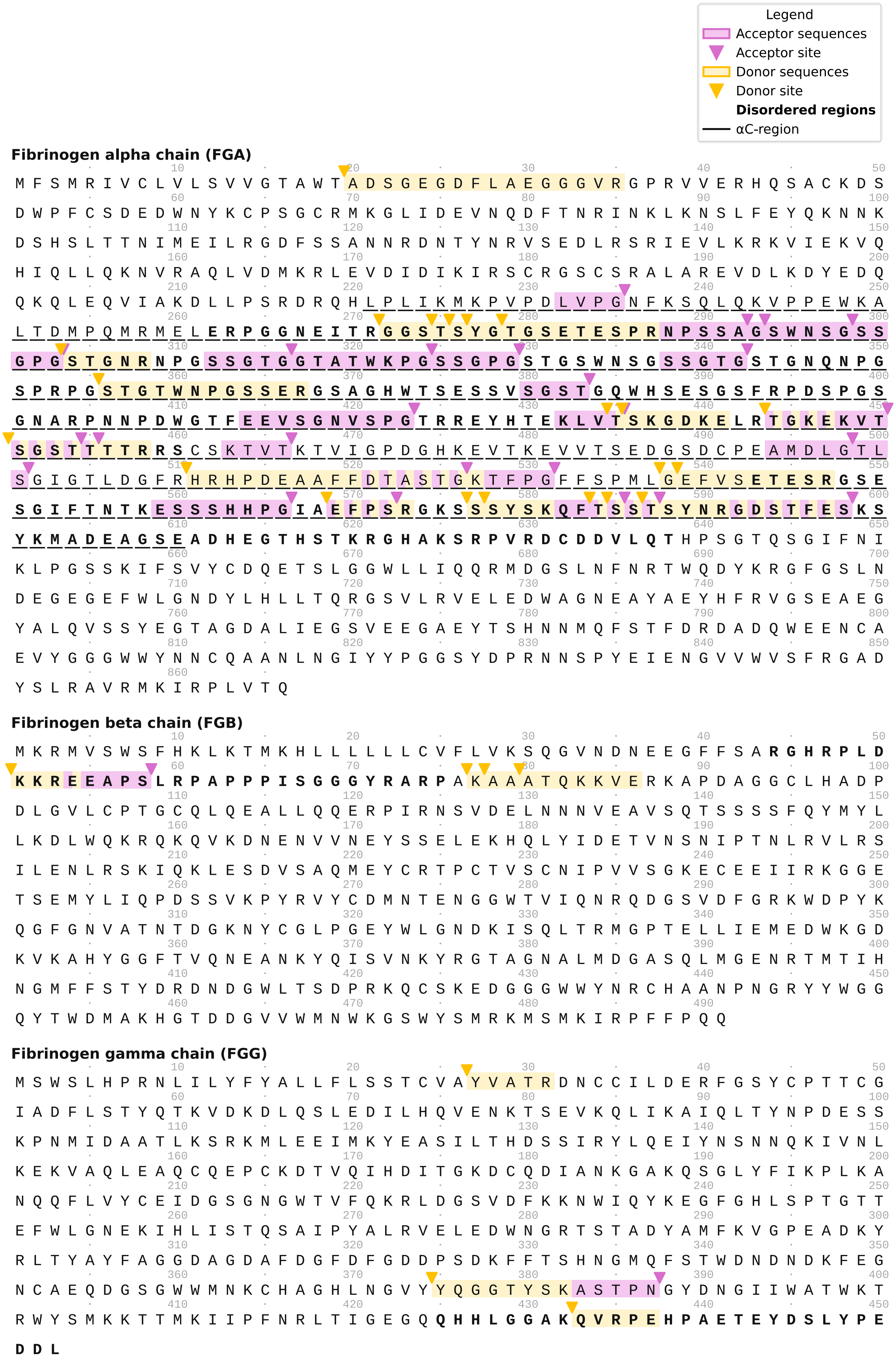


**Figure S12. Cispeptidation sites identified in fibrinogen after hCatK activity.** Bolded sequences indicate disordered regions in the primary protein sequence, according to region annotation in Uniprot of human fibrinogen α, β, and γ chains (Uniprot P02671, P02675, P02679). The underlined sequence represents the αC-region. Sequences highlighted in purple and yellow are acceptor and donor sequences, respectively, identified in confirmed cis-spliced peptides; and purple and yellow arrows indicate the identified cispeptidation acceptor and donor sites, respectively. Brown arrows indicate sites acting both as acceptor and donor.

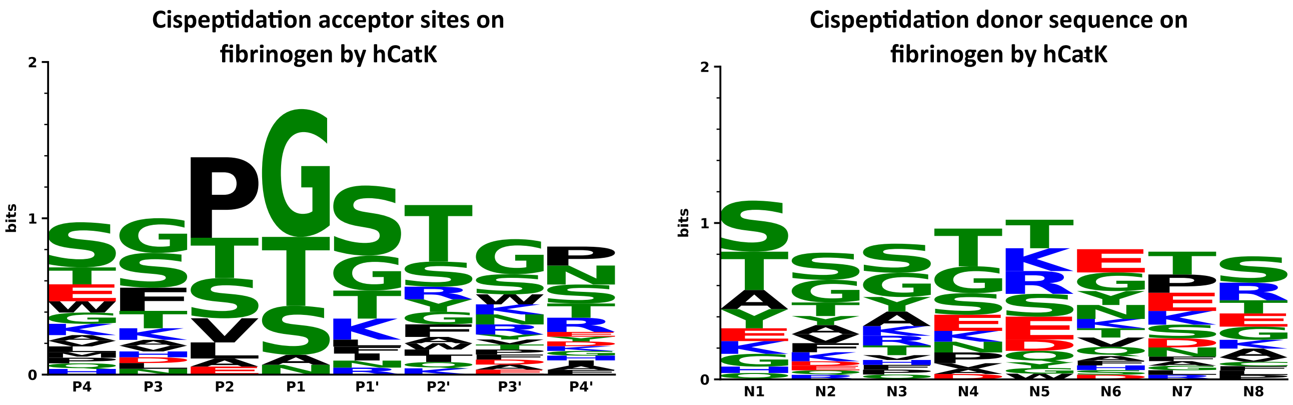


**Figure S13. Acceptor and donor sequences in fibrinogen involved in cispeptidation by hCatK.** Cispeptidation happens at the scissile bond between the P1 and P1' positions and uses the primary amine at the N-terminal end of N1 as the nucleophilic donor. Weblogos depict sequence conservation at each position for both types of sequences on fibrinogen. For complete cis-spliced peptide identities, see Tables 6 and S3.

Legend

Acceptor sequences

Acceptor site

Donor sequences

Donor site

**Disordered regions**

**S1 Protein (SARS-CoV-2)**

**Q C V N L T T R T Q L P P** A Y T N S F T R G V Y Y P D K V F R S S V L H S T Q D L F L P F F S N V T
W F H **A I H V S G T N G T K R F D** N P V L P F N D G V Y F A S T E K S N I I R G W I F G T T L D S K
T Q S L L I V N N A T N V V I K V C E F Q F C N D P F L **G V Y Y H K N N K S W M E S E F R V Y S S A**
**N N** C T F E Y V S **Q P F L M D L E G K Q G N** F K N L R E F V F K N I D G Y F K I Y S K H T P I N L V
R D L P Q G F S A L E P L V D L P I G I N I T R F Q T L L A L H **R S Y L T P G D S S S G W T A G A** A
A Y Y V G Y L Q P R T F L L K Y N E N G T I T D A V D C A L D P L S E T K C T L K S F T V E K G I Y
Q T S N F **R V Q P T E S I V R F P N I T N L** C P F G E V F N A T R F A S V Y A W N R K R I S N C V A
D Y S V L Y N S A S F S T F K C Y G V S P T K L N D L C F T N V Y A D S F V I R G D E V R Q I A P G
Q T G K I A D Y N Y K L P D D F T G C V I A W N S N N L D S K V G G N Y N Y L Y R **L F R K S N L K P**
**F E R D I S T E I Y Q A G S T P C N G V E G F N C Y F** P L Q S Y G F Q P T N G V G Y Q P Y R V V V L
S F E L L **H A P A T V C G P K K S T N L V K N K C V N F** N F N G L T G T G V L T E S N K K F L P F Q
Q F G R D I A D T T D A V R D P Q T L E I L D I T P C S F G G V S V I T P G T N T S N Q V A V L Y Q
D V N C T E V **P V A I H A D Q L T P T W R V Y S T G S** N V F Q T R A G C L I G A E H V N N S Y E C D
I P I G A G I C A **S Y Q T Q T N S P R R A R**

**S2 Protein (SARS-CoV-2)**

**S V** A S Q S I I A Y T M S L G A E N S V A Y S N N S I A I P T N F T I S V T T E I L P V S M T K T S
V D C T M Y I C G D S T E C S N L L L Q Y G S F C T Q L N R A L T G I A V E Q D K N T Q E V F A Q V
K Q I Y K T P P I K D F G G F N F S Q I L P D P S K P S K R S F I E D L L F N K V T **L A D A G F I K**
**Q Y G D C L G D I A A R D L I C A Q** K F N G L T V L P P L L T D E M I A Q Y T S A L L A G T I T S G
W T F G A G A A L Q I P F A M Q M A Y R F N G I G V T Q N V L Y E N Q K L I A N Q F N S A I G K I Q
D S L S S T A S A L G K L Q D V V N Q N A Q A L N T L V K Q L S S N F G A I S S V L N D I L S R L D
K V E A E V Q I D R L I T G R L Q S L Q T Y V T Q Q L I R A A E I R A S A N L A A T K M S E C V L G
Q S K R V D F C G K G Y H L M S F P Q S A P H G V V F L H V T Y V P A Q E K N F T T A P A I C H D G
K A H F P R E G V F V S N G T H W F V T Q R N F Y E P Q I I T T D N T F V S G N C D V V I G I V N N
T V Y D P L Q P E L D S F K E E L D K Y F K N H T S P D V D L G D I S G I N A S V V N I Q K E I D R
L N E V A K N L N E S L I D L Q E L G K Y E Q Y I K W P W Y I W L G F I A G L I A I V M V T I M L C
C M T S C C S C L K G C C S C G S C C K F D E D D S E P V L K G V K L H Y T

**Figure S14. Cispeptidation sites identified in Spike after hCatK activity.** Bolded sequences indicate disordered regions in the primary protein sequence, according to region annotation in Uniprot of SARS-CoV-2 Spike glycoprotein (Uniprot P0DTC2). Sequences highlighted in purple and yellow are acceptor and donor sequences, respectively, identified in confirmed cis-spliced peptides; and purple and yellow arrows indicate the identified cispeptidation acceptor and donor sites, respectively.

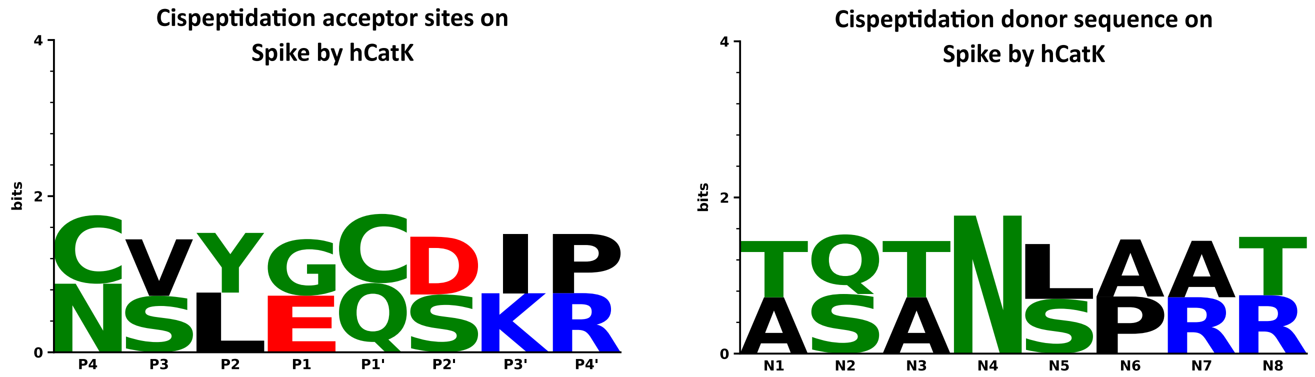


**Figure S15. Acceptor and donor sequences in Spike involved in cispeptidation by hCatK.** Cispeptidation happens at the scissile bond between the P1 and P1’ positions and uses the primary amine at the N-terminal end of N1 as the nucleophilic donor. Weblogos depict sequence conservation at each position for both types of sequences on Spike. For complete cis-spliced peptide identities, see Tables 7 and S3.

Legend
Acceptor sequences
Acceptor site
Donor sequences
Donor site
**Disordered regions**

**Alpha 1 chain of bovine type II collagen**

M I R L G A P Q T L V L L T L L V A A V L R C H G Q D V Q K A G S C V Q D G Q R Y N D K D V W K P E
P C R I C V C D T G T V L C D D I I C E D M K D C L S P E T P F G E C C P I C S A D L P T A S G Q P
G P K G Q K G E P G D I K D I V G P K G P P G P Q G P A G E Q G P R G D R G D K G E K G A P G P R G
R D G E P G T P G N P G P P G P P G P P G P P G L G G N F A A Q M A G G F D E K A G G A Q M G V M Q
G P M G P M G P R G P P G P A G A P G P Q G F Q G N P G E P G E P G V S G P M G P R G P P G P P G K
P G D D G E A G K P G K S G E R G P P G P Q G A R G F P G T P G L P G V K G H R G Y P G L D G A K G
E A G A P G V K G E S G S P G E N G S P G P M G P R G L P G E R G R T G P A G A A G A R G N D G Q P
G P A G P P G P V G P A G G P G F P G A P G A K G E A G P T G A R G P E G A Q G P R G E P G T P G S
P G P A G A A G N P G T D G I P G A K G S A G A P G I A G A P G F P G P R G P P G P Q G A T G P L G
P K G Q T G E P G I A G F K G E Q G P K G E P G P A G P Q G A P G P A G E E G K R G A R G E P G G A
G P A G P P G E R G A P G N R G F P G Q D G L A G P K G A P G E R G P S G L A G P K G A N G D P G R
P G E P G L P G A R G L T G R P G D A G P Q G K V G P S G A P G E D G R P G P P G P Q G A R G Q P G
V M G F P G P K G A N G E P G K A G E K G L P G A P G L R G L P G K D G E T G A A G P P G P A G P A
G E R G E Q G A P G P S G F Q G L P G P P G P P G E G G K P G D Q G V P G E A G A P G L V G P R G E
R G F P G E R G S P G S Q G L Q G A R G L P G T P G T D G P K G A A G P A G P P G A Q G P P G L Q G
M P G E R G A A G I A G P K G D R G D V G E K G P E G A P G K D G G R G L T G P I G P P G P A G A N
G E K G E V G P P G P A G T A G A R G A P G E R G E T G P P G P A G F A G P P G A D G Q P G A K G E
Q G E A G Q K G D A G A P G P Q G P S G A P G P Q G P T G V T G P K G A R G A Q G P P G A T G F P G
A A G R V G P P G S N G N P G P P G P P G P S G K D G P K G A R G D S G P P G R A G D P G L Q G P A
G P P G E K G E P G D D G P S G P D G P P G P Q G L A G Q R G I V G L P G Q R G E R G F P G L P G P
S G E P G K Q G A P G A S G D R G P P G P V G P P G L T G P A G E P G R E G S P G A D G P P G R D G
A A G V K G D R G E T G A V G A P G A P G P P G S P G P A G P I G K Q G D R G E A G A Q G P M G P A
G P A G A R G M P G P Q G P R G D K G E T G E A G E R G L K G H R G F T G L Q G L P G P P G P S G D
Q G A S G P A G P S G P R G P P G P V G P S G K D G A N G I P G P I G P P G P R G R S G E T G P A G
P P G N P G P P G P P G P P G P G I D M S A F A G L G Q R E K G P D P L Q Y M R A D E A A G N L R Q
H D A E V D A T L K S L N N Q I E S L R S P E G S R K N P A R T C R D L K L C H P E W K S G D Y W I
D P N Q G C T L D A M K V F C N M E T G E T C V Y P N P A S V P K K N W W S S K S K D K K H I W F G
E T I N G G F H F S Y G D D N L A P N T A N V Q M T F L R L L S T E G S Q N I T Y H C K N S I A Y L
D E A A G N L K K A L L I Q G S N D V E I R A E G N S R F T Y T V L K D G C T K H T G K W G K T M I
E Y R S Q K T S R L P I I D I A P M D I G G P E Q E F G V D I G P V C F L

**Figure S16. Transpeptidation sites identified in DTIIC, fibrinogen and Spike sequences after hCatK activity.** Bolded sequences indicate disordered regions in the primary protein sequences, according to region annotation in Uniprot of human fibrinogen α, β, and γ chains (Uniprot P02671, P02675, P02679), and SARS-CoV-2 Spike glycoprotein (Uniprot P0DTC2). The underlined sequence represents the αC-region of fibrinogen. Sequences highlighted in purple and yellow are acceptor and donor sequences, respectively, identified in confirmed trans-spliced peptides; and purple and yellow arrows indicate the identified transpeptidation acceptor and donor sites, respectively.

Legend

- Acceptor sequences
- Acceptor site
- Donor sequences
- Donor site
- **Disordered regions**
- αC-region

**Fibrinogen alpha chain (FGA)**

M F S M R I V C L V L S V V G T A W T A D S G E G D F L A E G G G V R G P R V V E R H Q S A C K D S

D W P F C S D E D W N Y K C P S G C R M K G L I D E V N Q D F T N R I N K L K N S L F E Y Q K N N K

D S H S L T T N I M E I L R G D F S S A N N R D N T Y N R V S E D L R S R I E V L K R K V I E K V Q

H I Q L L Q K N V R A Q L V D M K R L E V D I D I K I R S C R G S C S R A L A R E V D L K D Y E D Q

Q K Q L E Q V I A K D L L P S R D R Q H L P L I K M K P V P D L V P G N F K S Q L Q K V P P E W K A

L T D M P Q M R M E L E R P G G N E I T R G G S T S Y G T G S E T E S P R N P S S A G S W N S G S S

G P G S T G N R N P G S S G T G G T A T W K P G S S G P G S T G S W N S G S S G T G S T G N Q N P G

S P R P G S T G T W N P G S S E R G S A G H W T S E S S V S G S T G Q W H S E S G S F R P D S P G S

G N A R P N N P D W G T F E E V S G N V S P G T R R E Y H T E K L V T S K G D K E L R T G K E K V T

S G S T T T T R R S C S K T V T K T V I G P D G H K E V T K E V V T S E D G S D C P E A M D L G T L

S G I G T L D G F R H R H P D E A A F F D T A S T G K T F P G F F S P M L G E F V S E T E S R G S E

S G I F T N T K E S S S H H P G I A E F P S R G K S S S Y S K Q F T S S T S Y N R G D S T F E S K S

Y K M A D E A G S E A D H E G T H S T K R G H A K S R P V R D C D D V L Q T H P S G T Q S G I F N I

K L P G S S K I F S V Y C D Q E T S L G G W L L I Q Q R M D G S L N F N R T W Q D Y K R G F G S L N

D E G E G E F W L G N D Y L H L L T Q R G S V L R V E L E D W A G N E A Y A E Y H F R V G S E A E G

Y A L Q V S S Y E G T A G D A L I E G S V E E G A E Y T S H N N M Q F S T F D R D A D Q W E E N C A

E V Y G G G W W Y N N C Q A A N L N G I Y Y P G G S Y D P R N N S P Y E I E N G V V W V S F R G A D

Y S L R A V R M K I R P L V T Q

**Fibrinogen beta chain (FGB)**

M K R M V S W S F H K L K T M K H L L L L L L C V F L V K S Q G V N D N E E G F F S A R G H R P L D

K K R E E A P S L R P A P P P I S G G G Y R A R P A K A A A T Q K K V E R K A P D A G G C L H A D P

D L G V L C P T G C Q L Q E A L L Q Q E R P I R N S V D E L N N N V E A V S Q T S S S S F Q Y M Y L

L K D L W Q K R Q K Q V K D N E N V V N E Y S S E L E K H Q L Y I D E T V N S N I P T N L R V L R S

I L E N L R S K I Q K L E S D V S A Q M E Y C R T P C T V S C N I P V V S G K E C E E I I R K G G E

T S E M Y L I Q P D S S V K P Y R V Y C D M N T E N G G W T V I Q N R Q D G S V D F G R K W D P Y K

Q G F G N V A T N T D G K N Y C G L P G E Y W L G N D K I S Q L T R M G P T E L L I E M E D W K G D

K V K A H Y G G F T V Q N E A N K Y Q I S V N K Y R G T A G N A L M D G A S Q L M G E N R T M T I H

N G M F F S T Y D R D N D G W L T S D P R K Q C S K E D G G G W W Y N R C H A A N P N G R Y Y W G G

Q Y T W D M A K H G T D D G V V W M N W K G S W Y S M R K M S M K I R P F F P Q Q

**Fibrinogen gamma chain (FGG)**

M S W S L H P R N L I L Y F Y A L L F L S S T C V A Y V A T R D N C C I L D E R F G S Y C P T T C G

I A D F L S T Y Q T K V D K D L Q S L E D I L H Q V E N K T S E V K Q L I K A I Q L T Y N P D E S S

K P N M I D A A T L K S R K M L E E I M K Y E A S I L T H D S S I R Y L Q E I Y N S N N Q K I V N L

K E K V A Q L E A Q C Q E P C K D T V Q I H D I T G K D C Q D I A N K G A K Q S G L Y F I K P L K A

N Q Q F L V Y C E I D G S G N G W T V F Q K R L D G S V D F K K N W I Q Y K E G F G H L S P T G T T

E F W L G N E K I H L I S T Q S A I P Y A L R V E L E D W N G R T S T A D Y A M F K V G P E A D K Y

R L T Y A Y F A G G D A G D A F D G F D F G D D P S D K F F T S H N G M Q F S T W D N D N D K F E G

N C A E Q D G S G W W M N K C H A G H L N G V Y Y Q G G T Y S K A S T P N G Y D N G I I W A T W K T

R W Y S M K K T T M K I I P F N R L T I G E G Q Q H H L G G A K Q V R P E H P A E T E Y D S L Y P E

D D L

**Figure S16. (continued)**

Legend
Acceptor sequences
Acceptor site
Donor sequences
Donor site
**Disordered regions**

**S1 Protein (SARS-CoV-2)**

**Q C V N L T T R T Q L P P** A Y T N S F T R G V Y Y P D K V F R S S V L H S T Q D L F L P F F S N V T
W F H **A I H V S G T N G T K R F D** N P V L P F N D G V Y F A S T E K S N I I R G W I F G T T L D S K
T Q S L L I V N N A T N V V I K V C E F Q F C N D P F L **G V Y Y H K N N K S W M E S E F R V Y S S A**
**N N** C T F E Y V S **Q P F L M D L E G K Q G N** F K N L R E F V F K N I D G Y F K I Y S K H T P I N L V
R D L P Q G F S A L E P L V D L P I G I N I T R F Q T L L A L H **R S Y L T P G D S S S G W T A G A** A
A Y Y V G Y L Q P R T F L L K Y N E N G T I T D A V D C A L D P L S E T K C T L K S F T V E K G I Y
Q T S N F **R V Q P T E S I V R F P N I T N L** C P F G E V F N A T R F A S V Y A W N R K R I S N C V A
D Y S V L Y N S A S F S T F K C Y G V S P T K L N D L C F T N V Y A D S F V I R G D E V R Q I A P G
Q T G K I A D Y N Y K L P D D F T G C V I A W N S N N L D S K V G G N Y N Y L Y R **L F R K S N L K P**
**F E R D I S T E I Y Q A G S T P C N G V E G F N C Y F** P L Q S Y G F Q P T N G V G Y Q P Y R V V V L
S F E L L **H A P A T V C G P K K S T N L V K N K C V N F** N F N G L T G T G V L T E S N K K F L P F Q
Q F G R D I A D T T D A V R D P Q T L E I L D I T P C S F G G V S V I T P G T N T S N Q V A V L Y Q
D V N C T E V **P V A I H A D Q L T P T W R V Y S T G S** N V F Q T R A G C L I G A E H V N N S Y E C D
I P I G A G I C A **S Y Q T Q T N S P R R A R**

**S2 Protein (SARS-CoV-2)**

**S V** A S Q S I I A Y T M S L G A E N S V A Y S N N S I A I P T N F T I S V T T E I L P V S M T K T S
V D C T M Y I C G D S T E C S N L L L Q Y G S F C T Q L N R A L T G I A V E Q D K N T Q E V F A Q V
K Q I Y K T P P I K D F G G F N F S Q I L P D P S K P S K R S F I E D L L F N K V T **L A D A G F I K**
**Q Y G D C L G D I A A R D L I C A Q** K F N G L T V L P P L L T D E M I A Q Y T S A L L A G T I T S G
W T F G A G A A L Q I P F A M Q M A Y R F N G I G V T Q N V L Y E N Q K L I A N Q F N S A I G K I Q
D S L S S T A S A L G K L Q D V V N Q N A Q A L N T L V K Q L S S N F G A I S S V L N D I L S R L D
K V E A E V Q I D R L I T G R L Q S L Q T Y V T Q Q L I R A A E I R A S A N L A A T K M S E C V L G
Q S K R V D F C G K G Y H L M S F P Q S A P H G V V F L H V T Y V P A Q E K N F T T A P A I C H D G
K A H F P R E G V F V S N G T H W F V T Q R N F Y E P Q I I T T D N T F V S G N C D V V I G I V N N
T V Y D P L Q P E L D S F K E E L D K Y F K N H T S P D V D L G D I S G I N A S V V N I Q K E I D R
L N E V A K N L N E S L I D L Q E L G K Y E Q Y I K W P W Y I W L G F I A G L I A I V M V T I M L C
C M T S C C S C L K G C C S C G S C C K F D E D D S E P V L K G V K L H Y T

**Figure S16. (continued)**

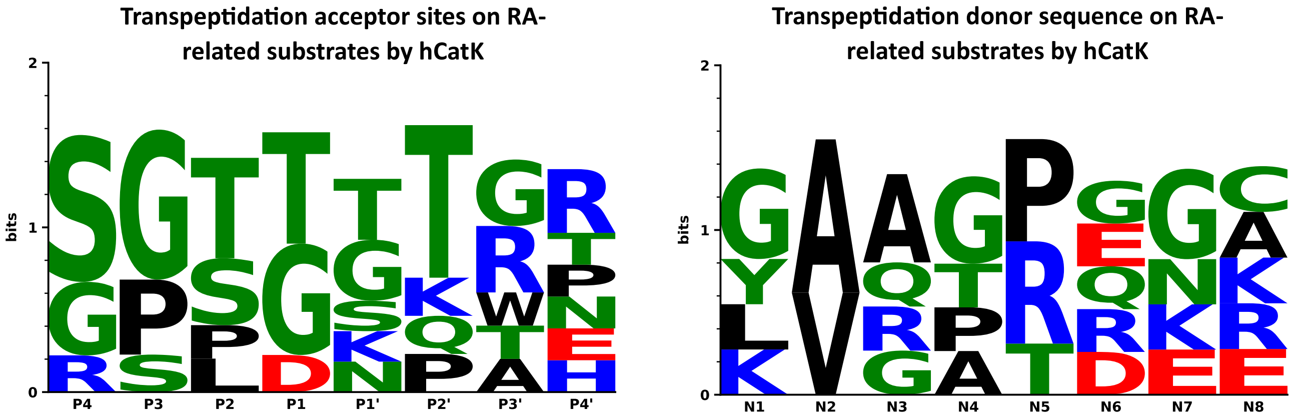


**Figure S17. Acceptor and donor sequences in DTIIC, fibrinogen, and Spike involved in transpeptidation by hCatK.** Transpeptidation happens at the scissile bond between the P1 and P1' positions and uses the primary amine at the N-terminal end of N1 as the nucleophilic donor. Weblogos depict sequence conservation at each position for both types of sequences on DTIIC, fibrinogen, and Spike. For complete trans-spliced peptide identities, see Tables 8 and S3.

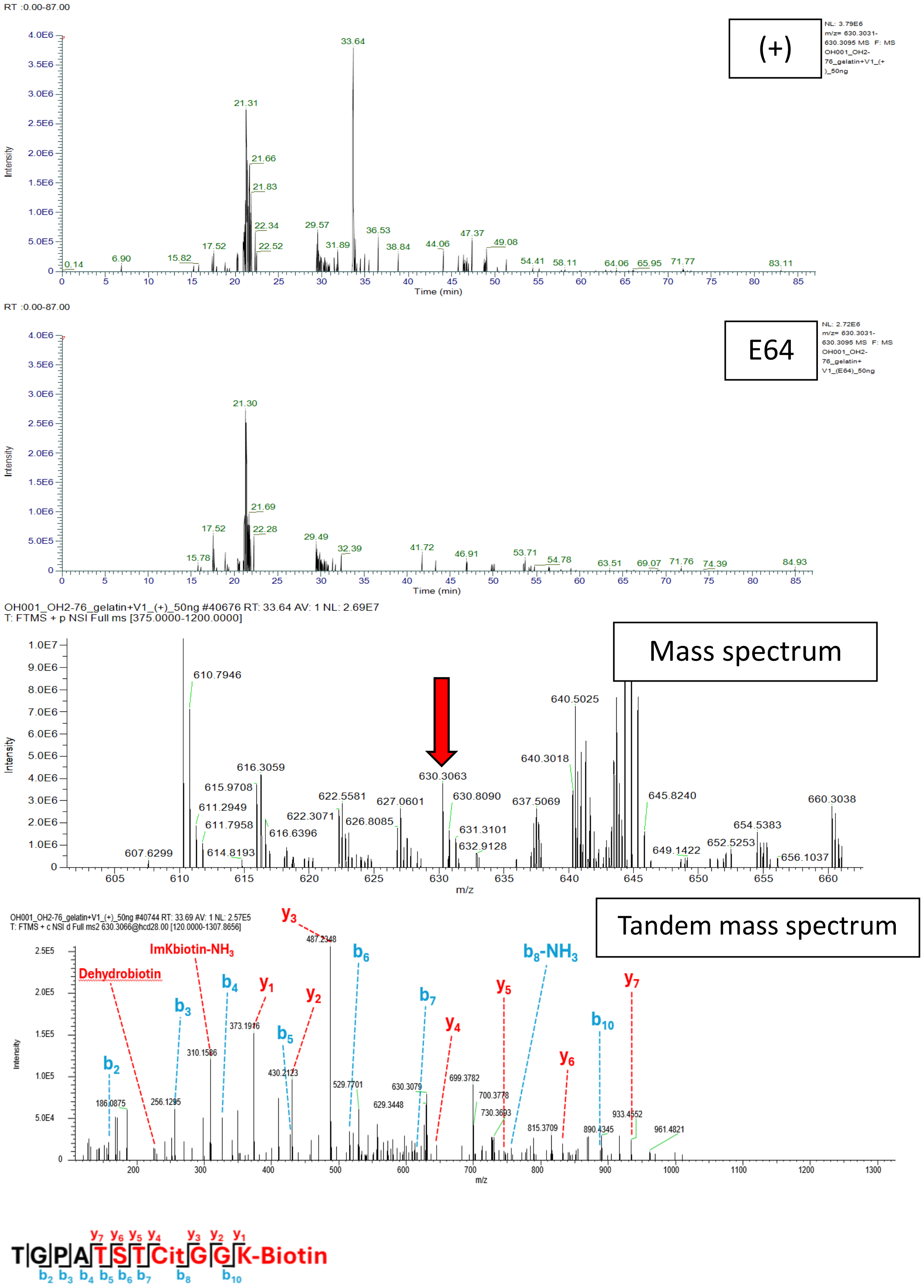


**Figure S18A. TGPATSTCitGGK-Biotin; Scan 40744; RT = 33.64; m/z = 630.3063. DTIIC and V1.**

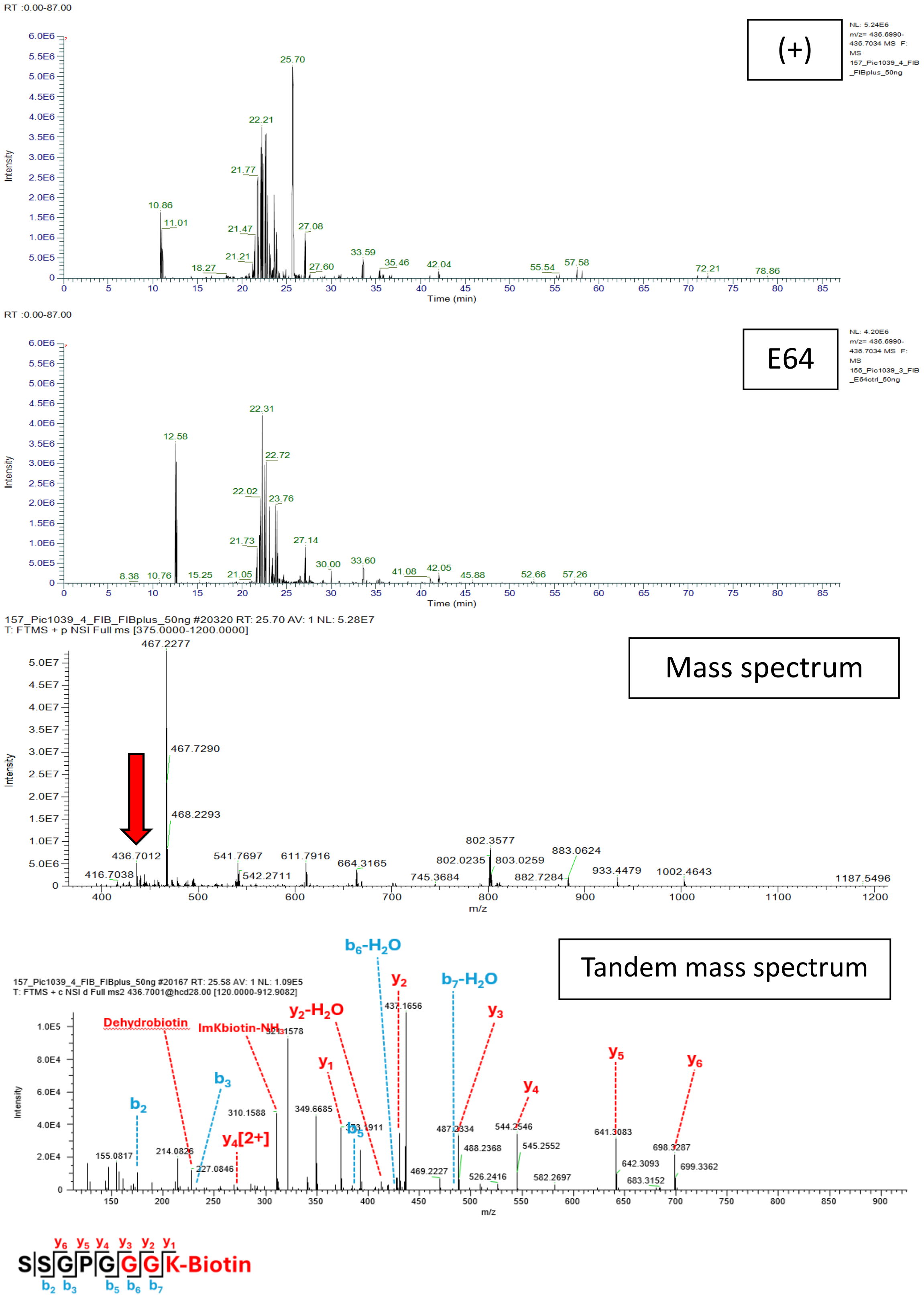


**Figure S18B. SSGPGGGK-Biotin; Scan 20167; RT = 25.70; m/z = 436.7012. Fibrinogen and V1.**

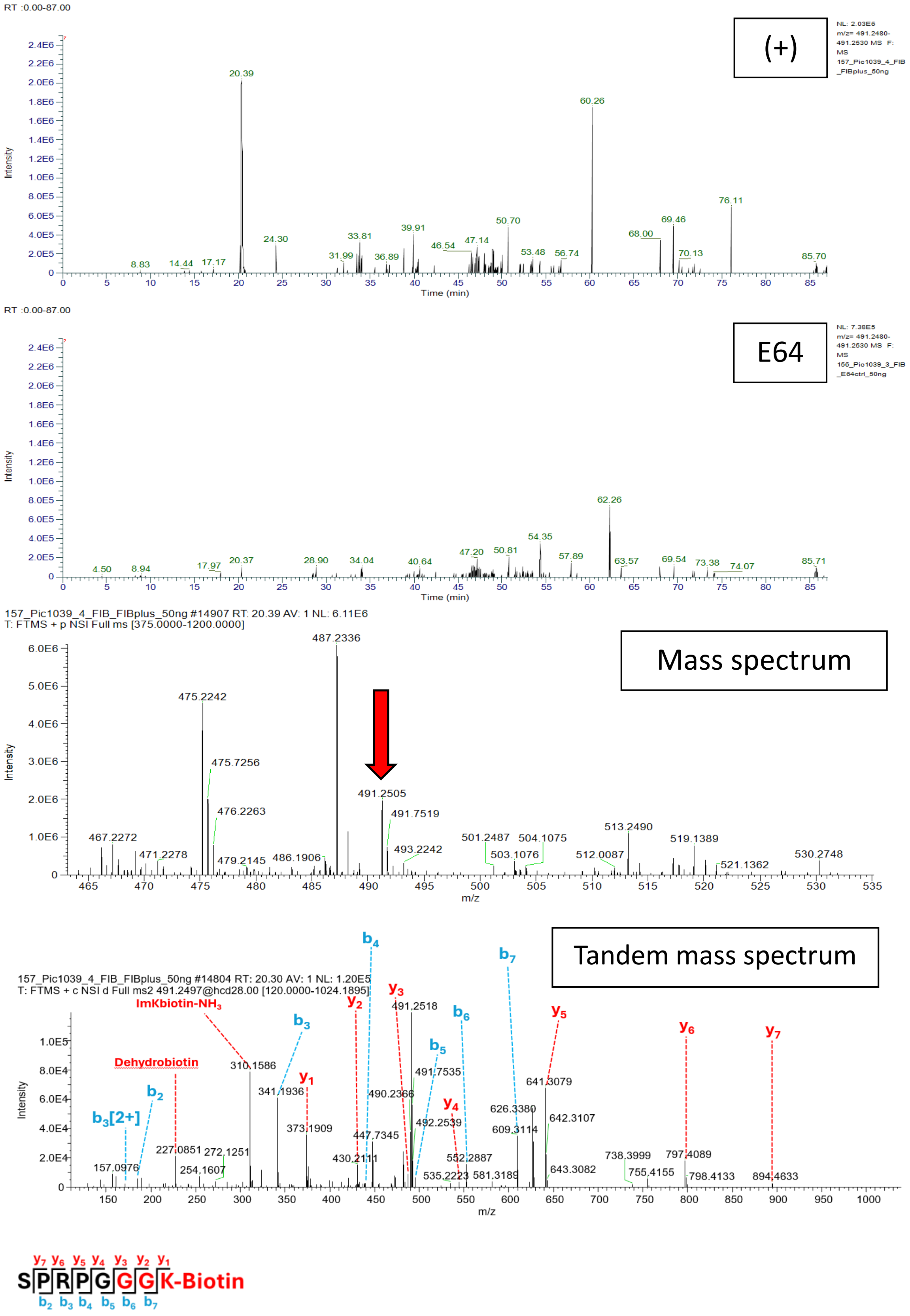


**Figure S18C. SPRPGGGK-Biotin; Scan 14804; RT = 20.39; m/z = 491.2505. Fibrinogen and V1.**

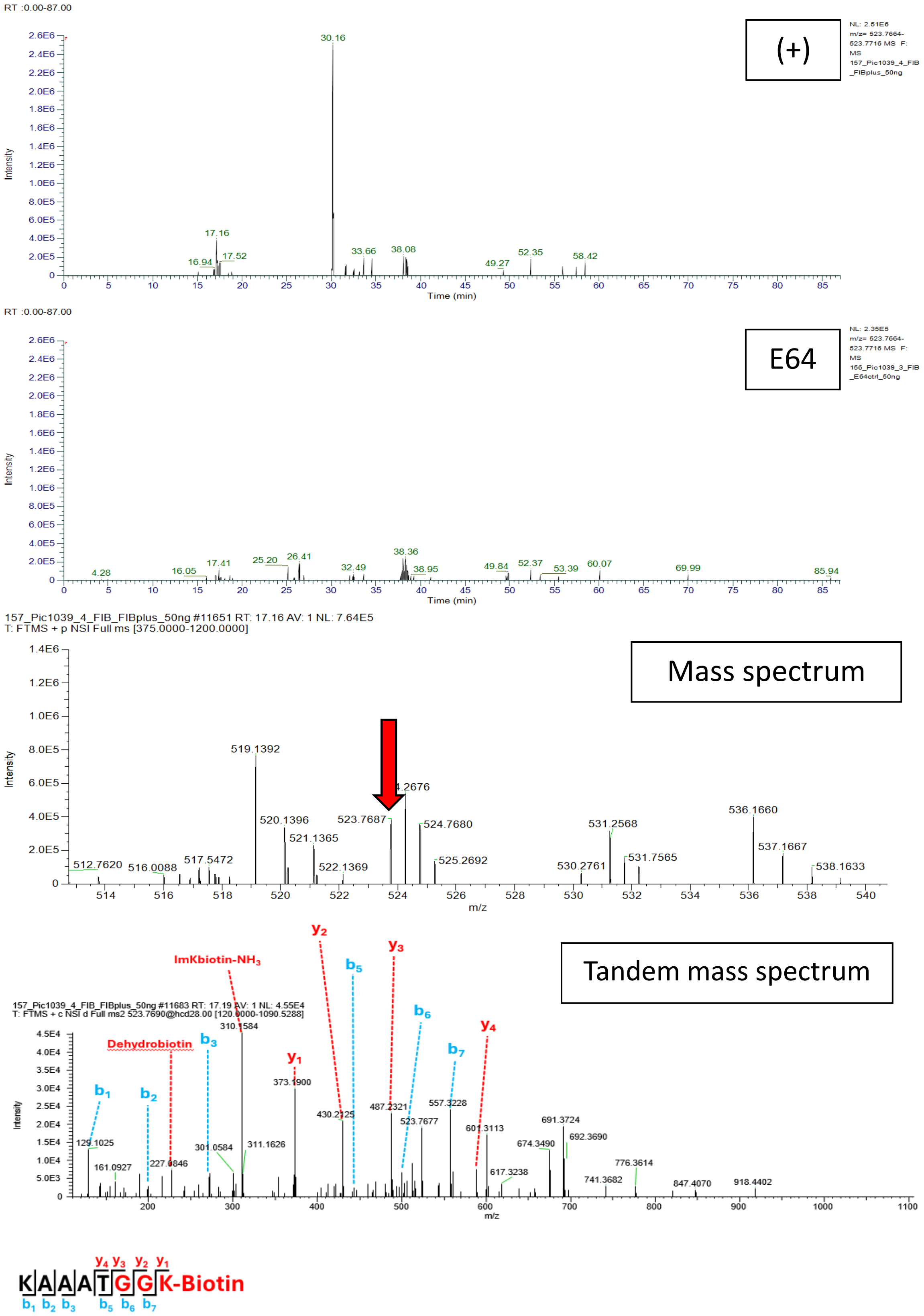


**Figure S18D. KAAATGGK-Biotin; Scan 11683; RT = 17.19; m/z = 523.7690. Fibrinogen and V1.**

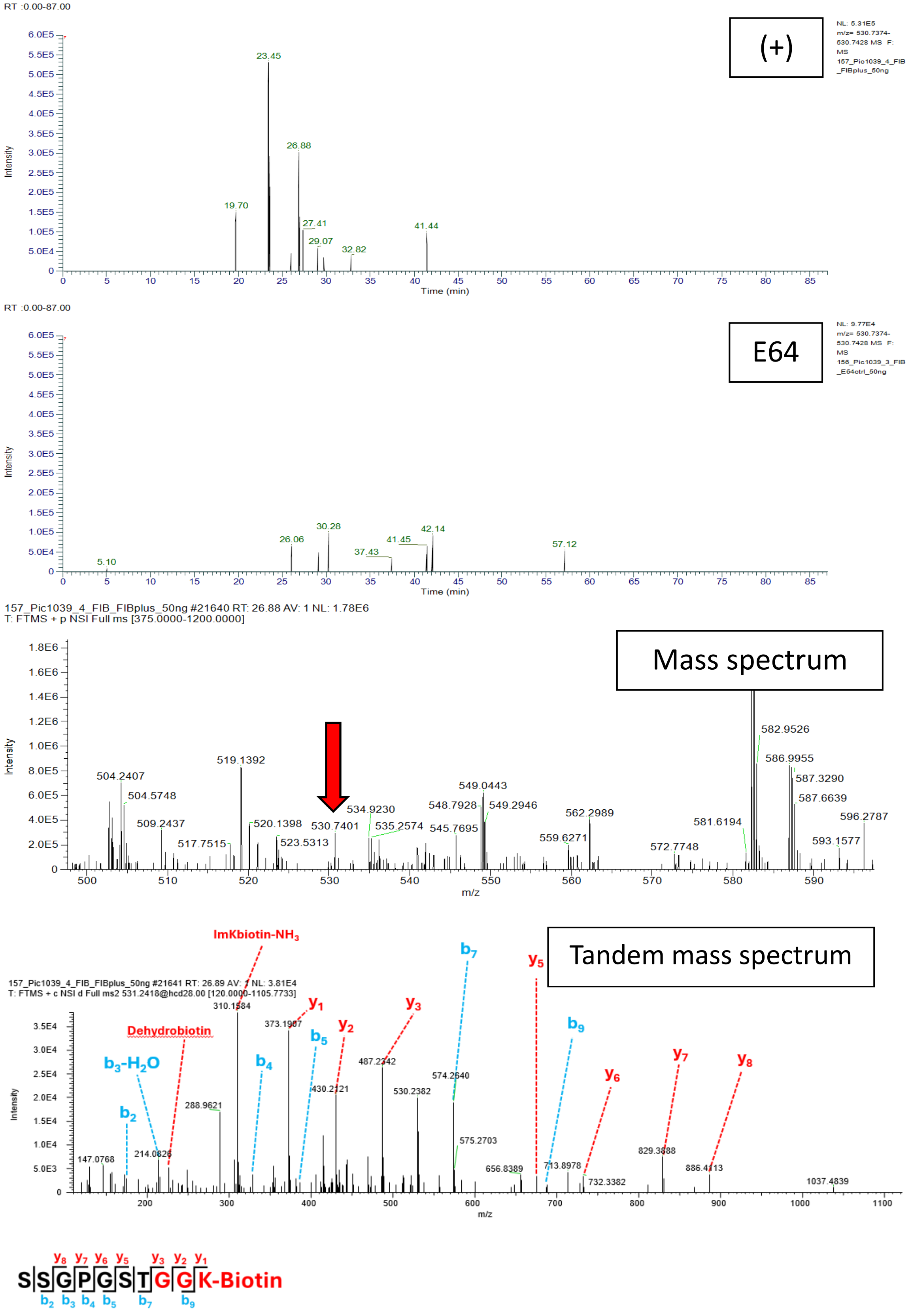


**Figure S18E. SSGPGSTGGK-Biotin; Scan 21641; RT = 26.89; m/z = 530.7401. Fibrinogen and V1.**

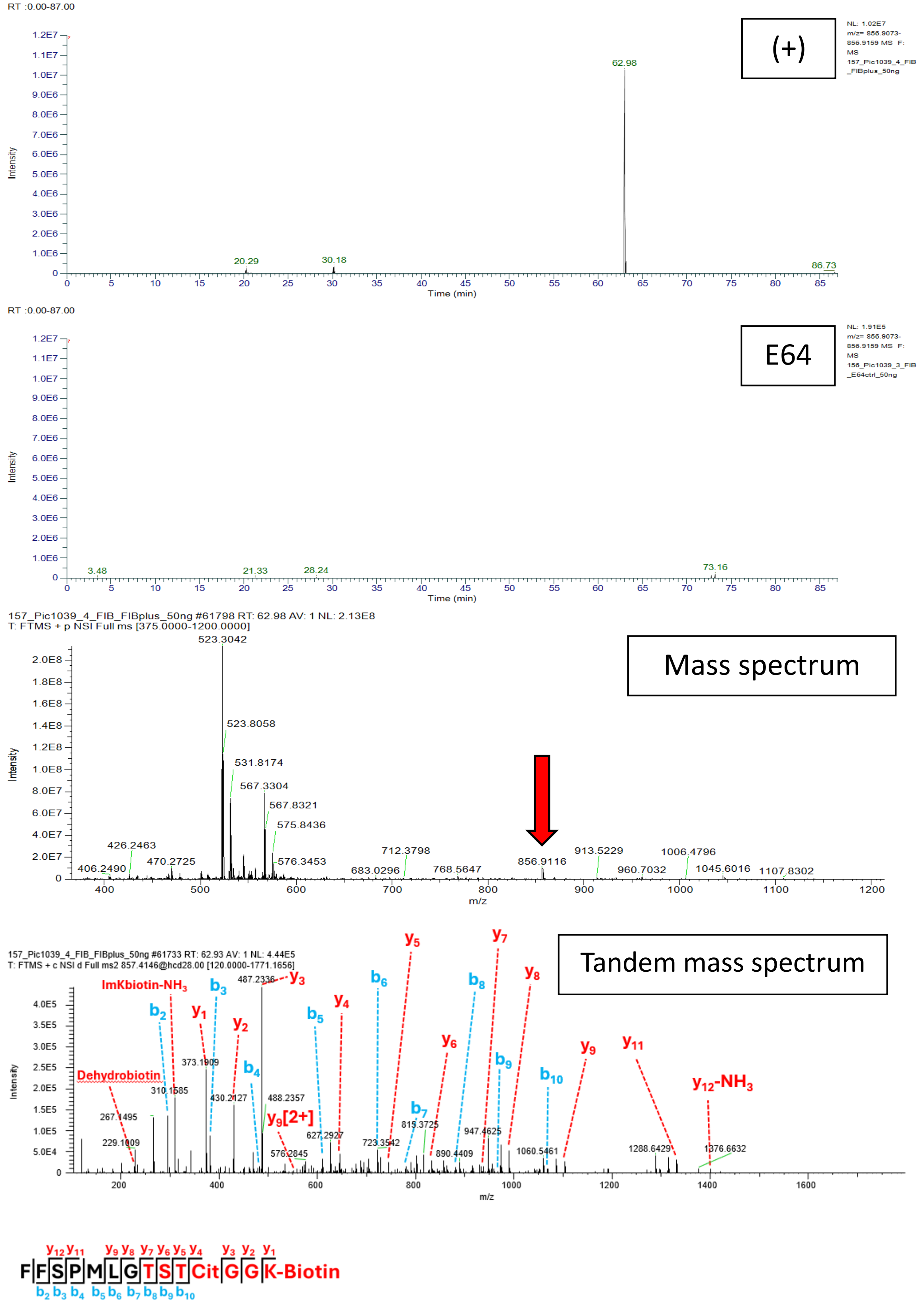


**Figure S18F. FFSPMLGTSTCitGGK-Biotin; Scan 61733; RT = 62.98; m/z = 856.9116. Fibrinogen and V1.**

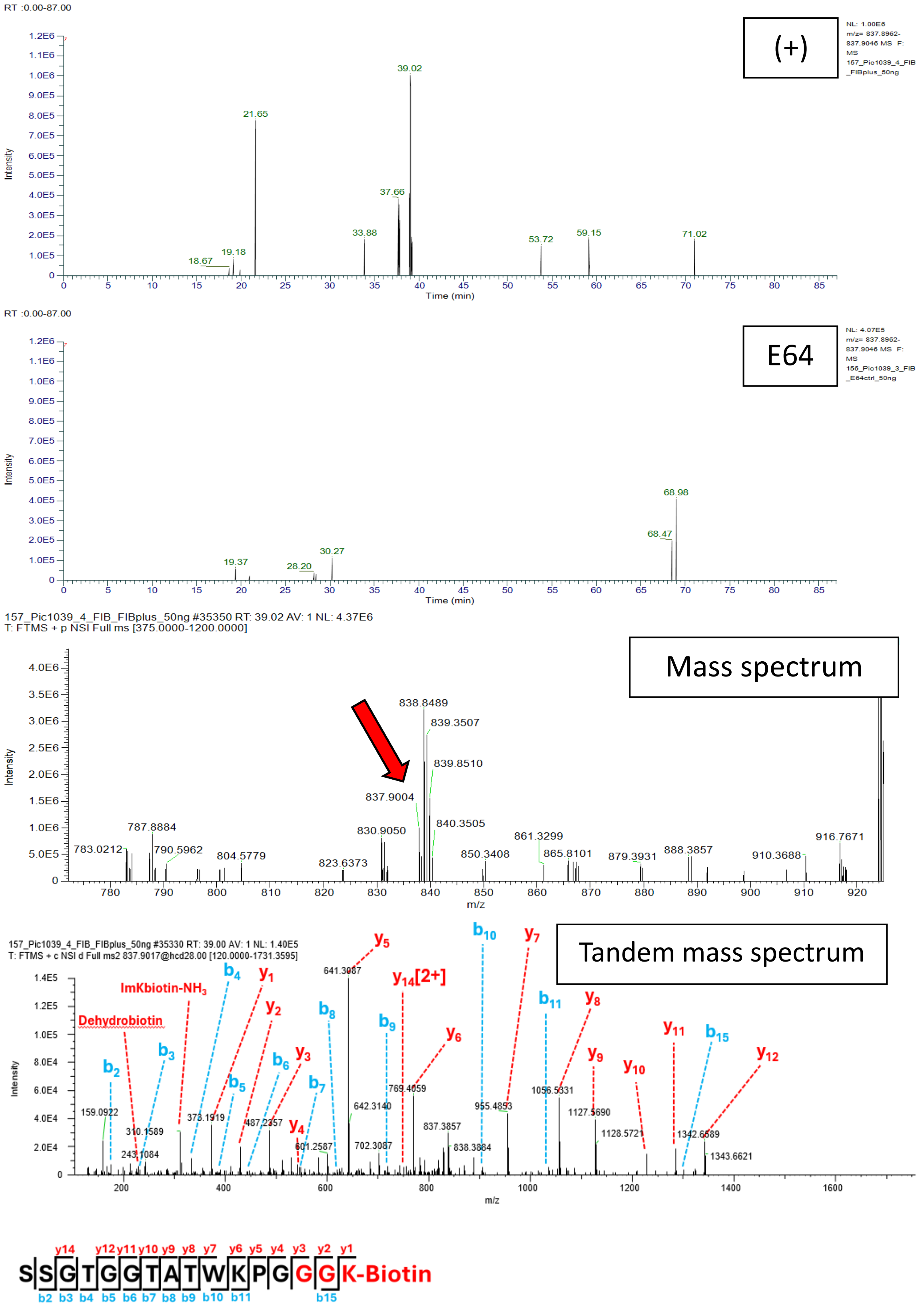


**Figure S18G. SSGTGGTATWKPGGGK-Biotin; Scan 35330; RT = 39.02; m/z = 837.9004. Fibrinogen and V1.**

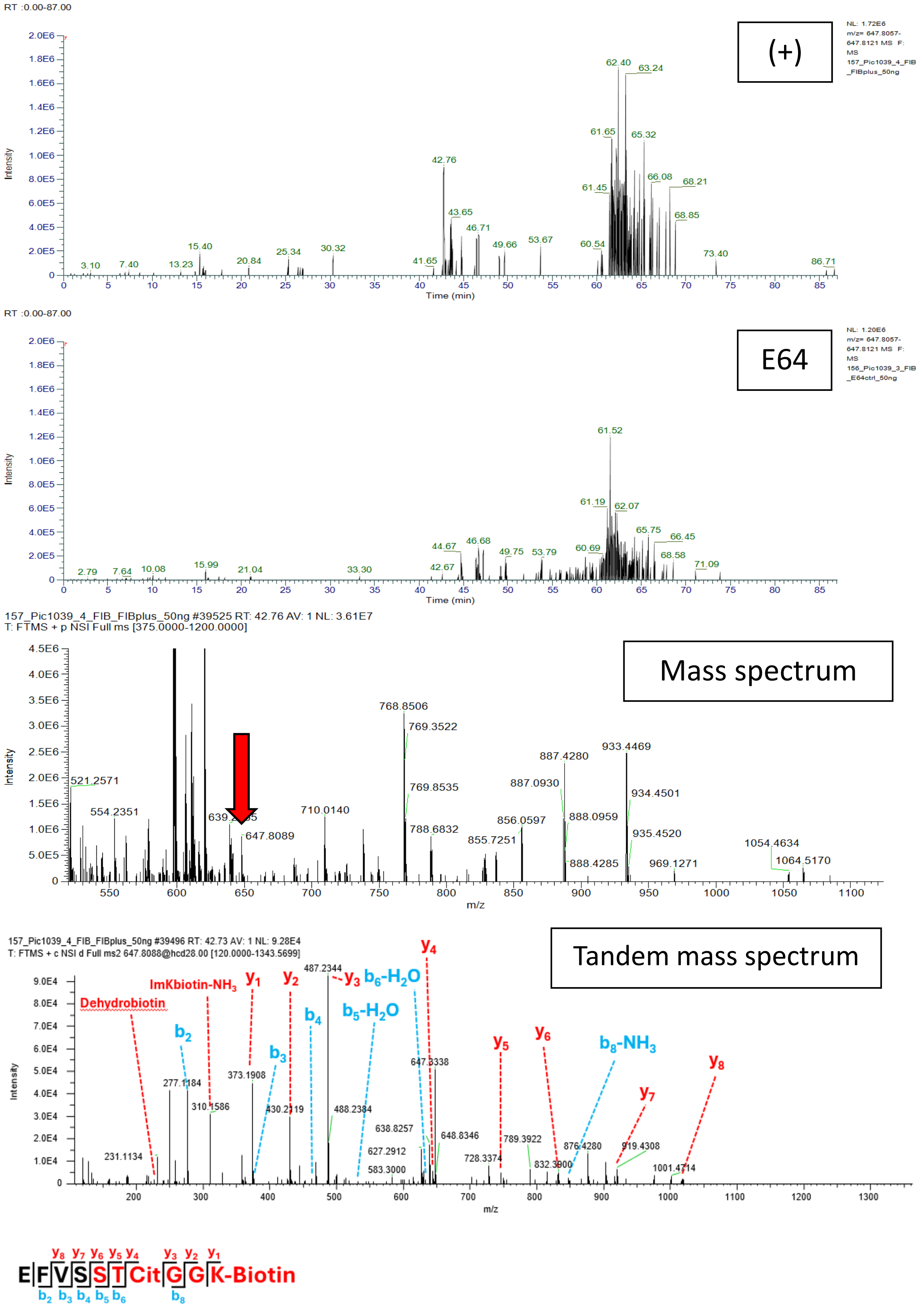


**Figure S18H. EFVSSTCitGGK-Biotin; Scan 39496; RT = 42.76; m/z = 647.8089. Fibrinogen and V1.**

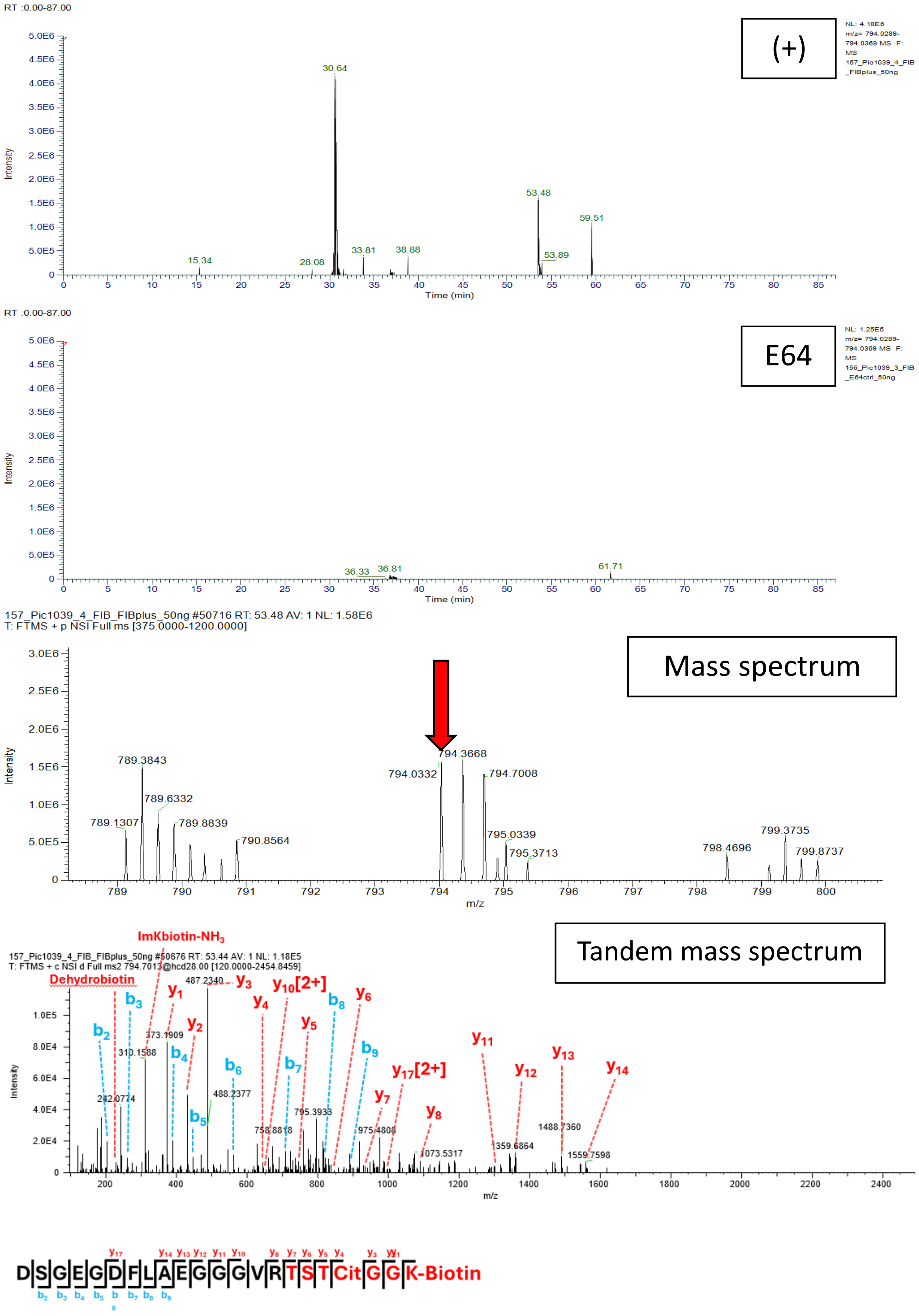


**Figure S18I. DSGEGDFLAEGGGVRTSTCitGGK-Biotin; Scan 50676; RT = 53.46; m/z = 794.0329. Fibrinogen and V1.**

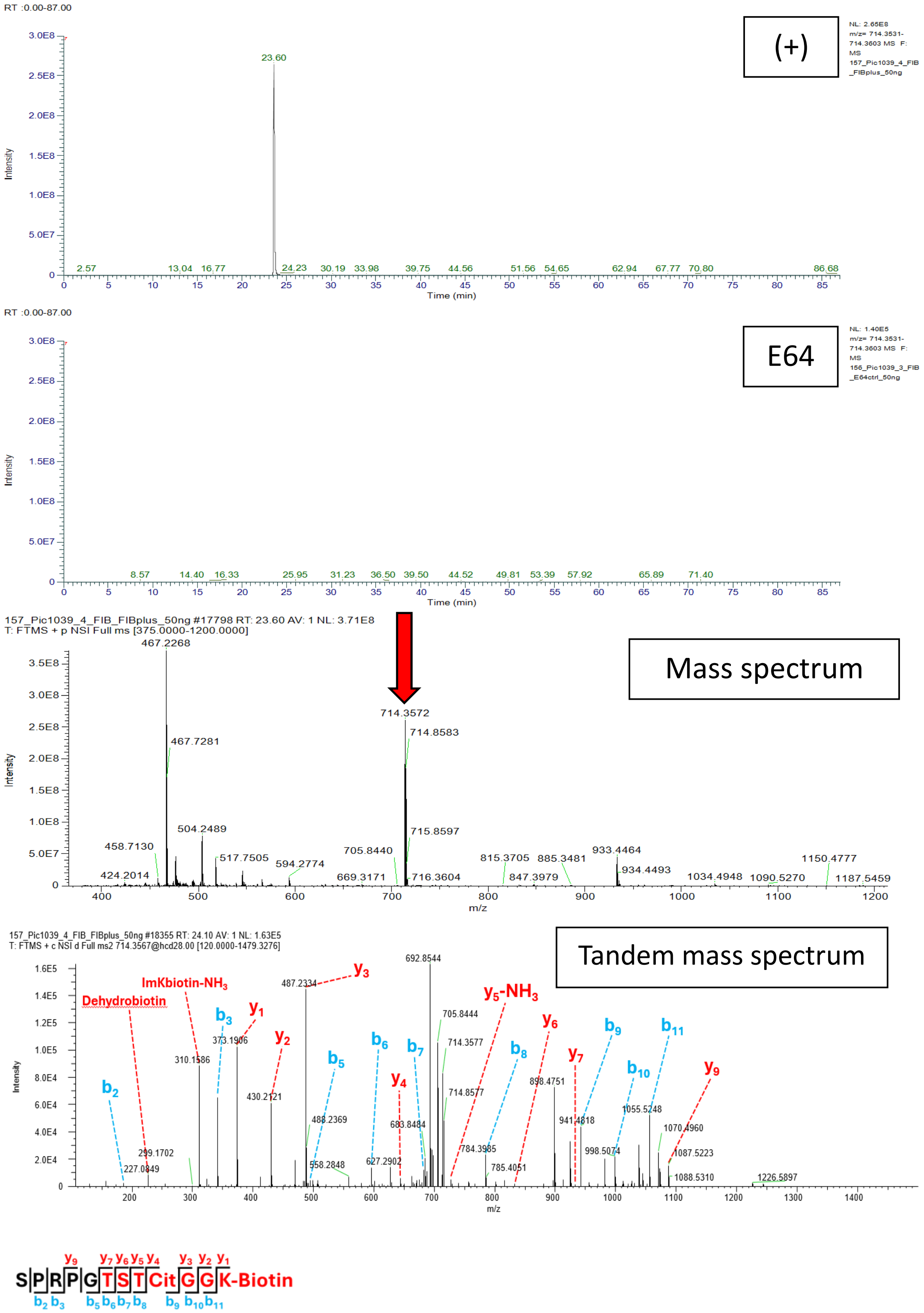


**Figure S18J. SPRPGTSTCitGGK-Biotin; Scan 18355; RT = 24.10; m/z = 714.3567. Fibrinogen and V1.**

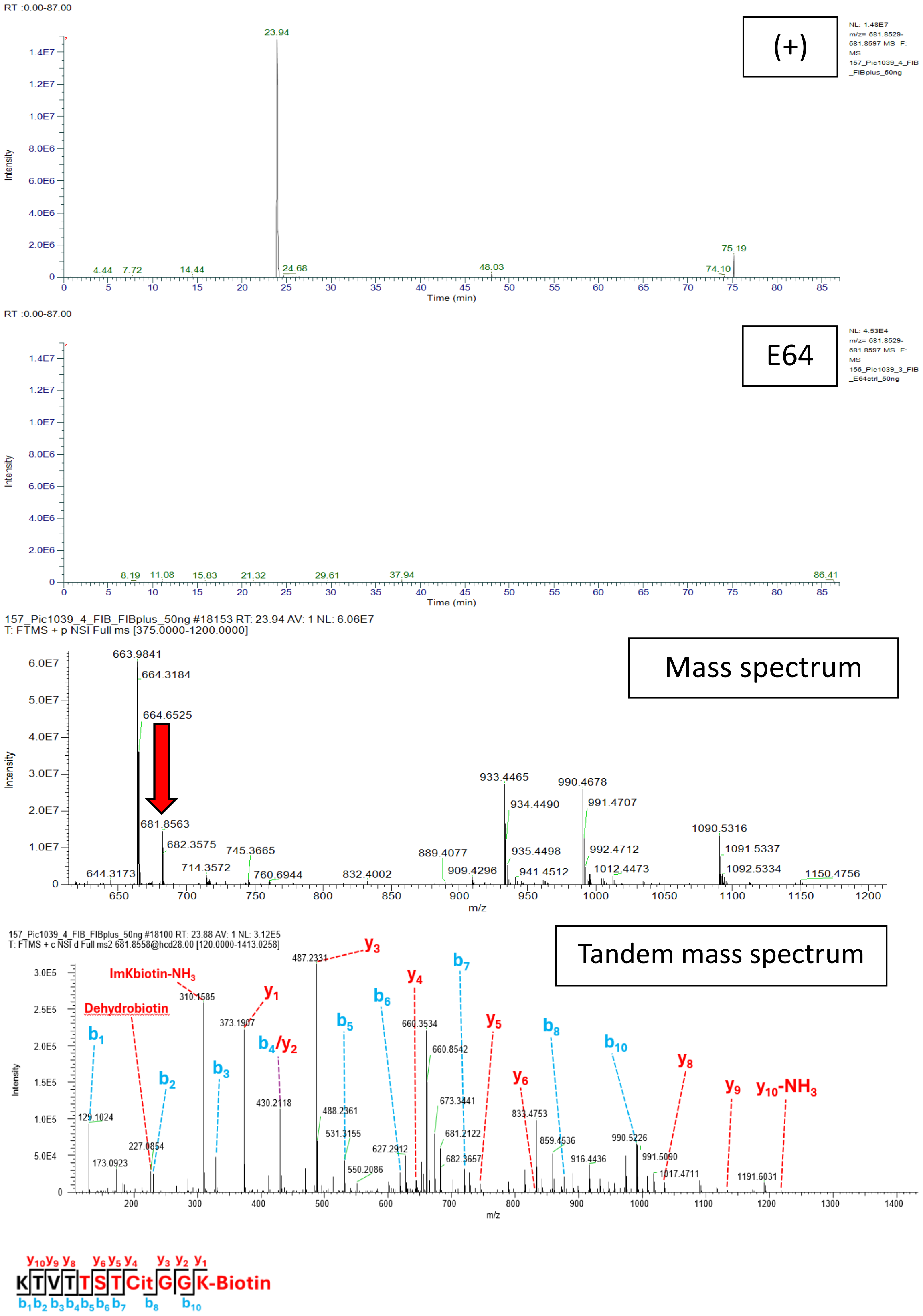


**Figure S18K. KTVTTSTCitGGK-Biotin; Scan 18100; RT = 23.94; m/z = 681.8563. Fibrinogen and V1.**

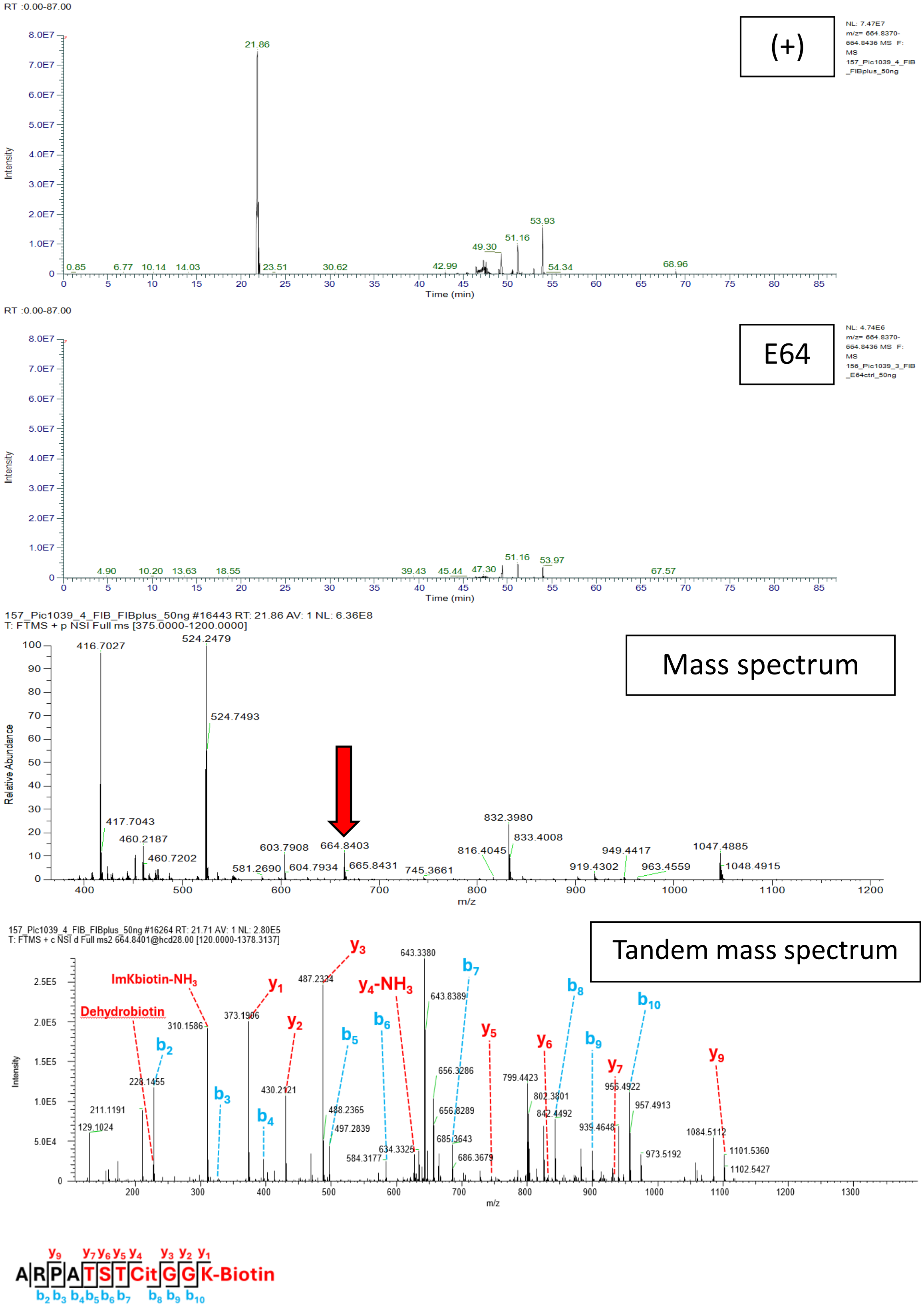


**Figure S18L. ARPATSTCitGGK-Biotin; Scan 16264; RT = 21.86; m/z = 664.8403. Fibrinogen and V1.**

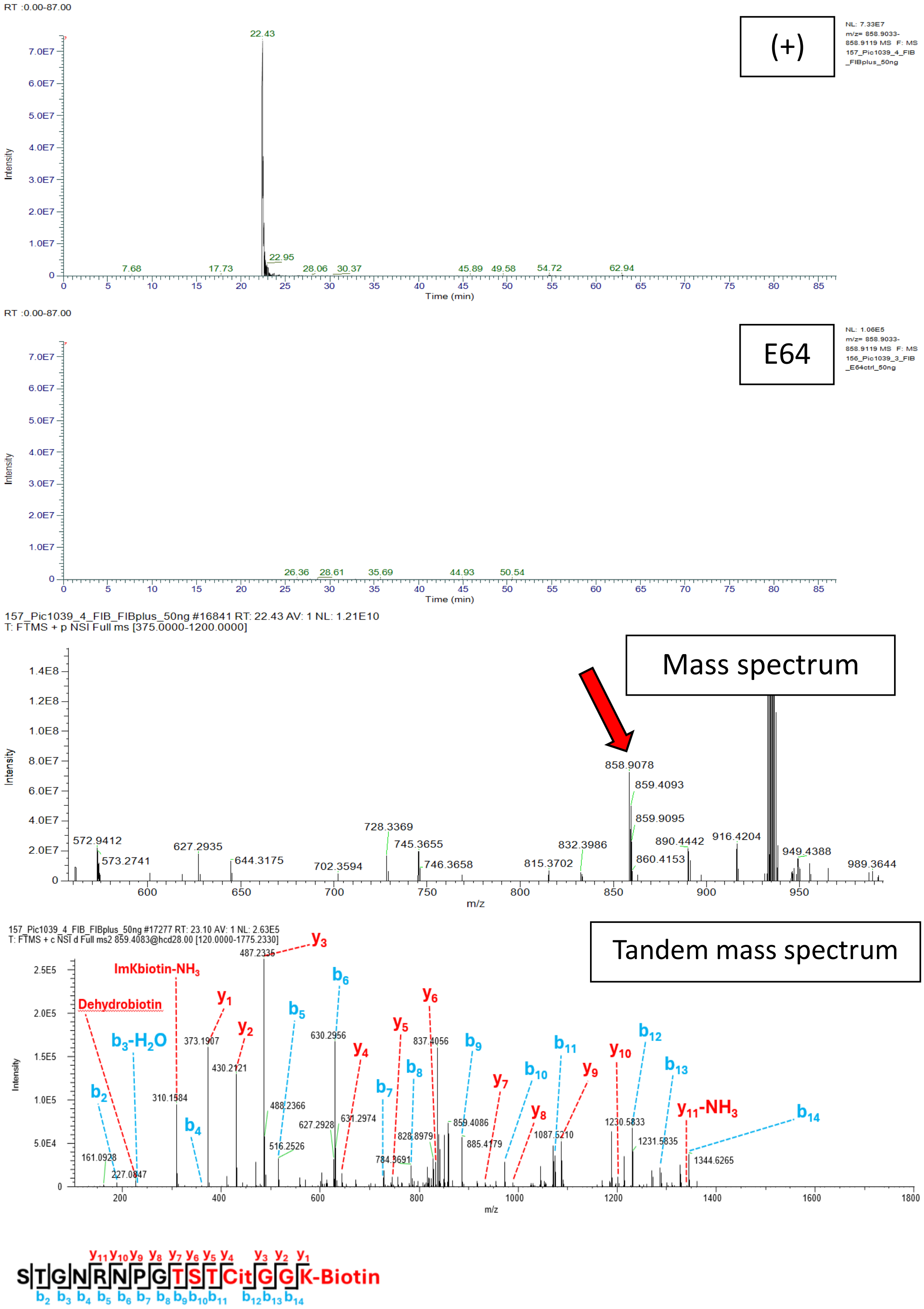


**Figure S18M. STGNRNPGTSTCitGGK-Biotin; Scan 17277; RT = 23.09; m/z = 858.9076. Fibrinogen and V1.**

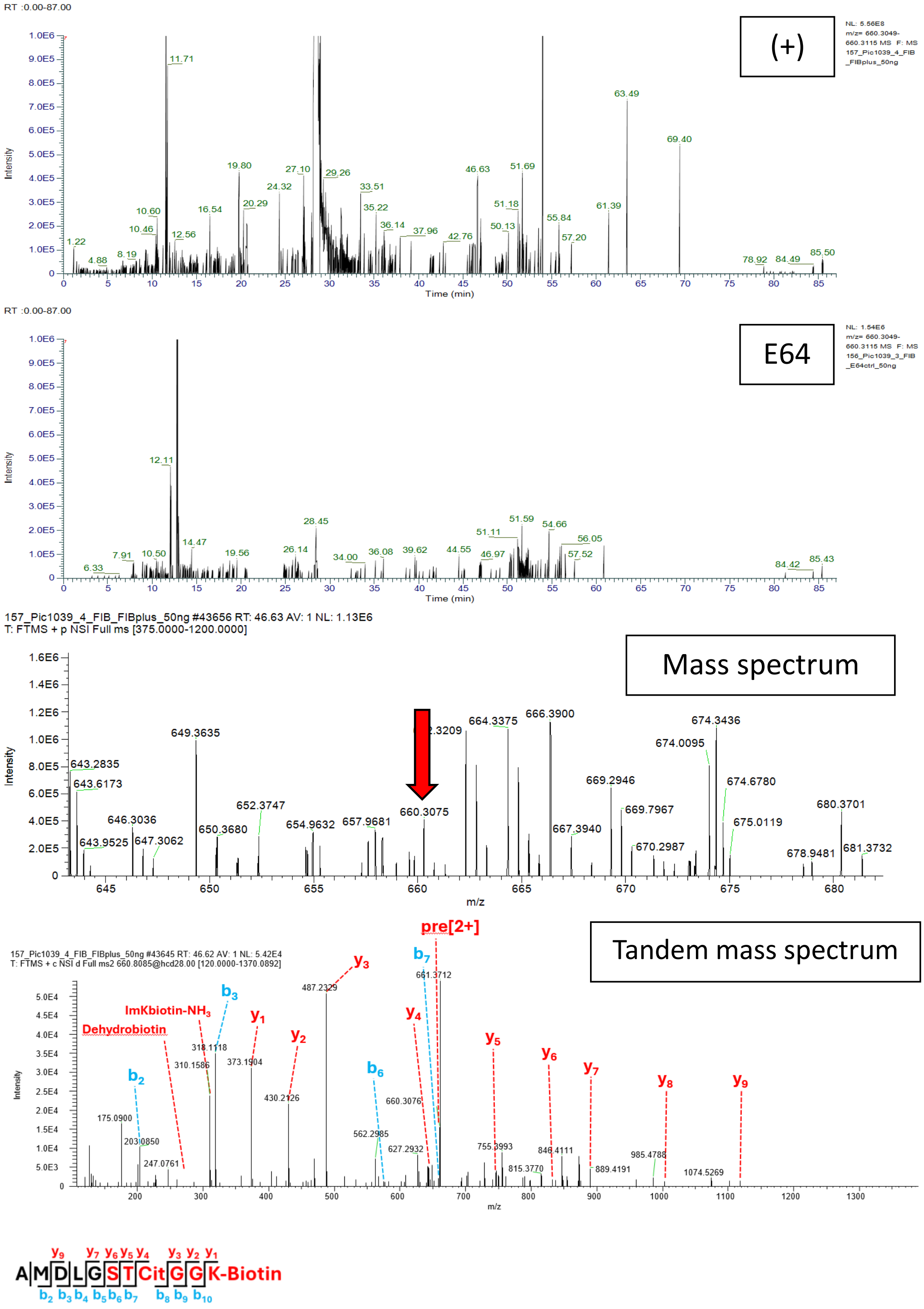


**Figure S18N. AMDLGSTCitGGK-Biotin; Scan 43645; RT = 46.64; m/z = 660.3082. Fibrinogen and V1.**

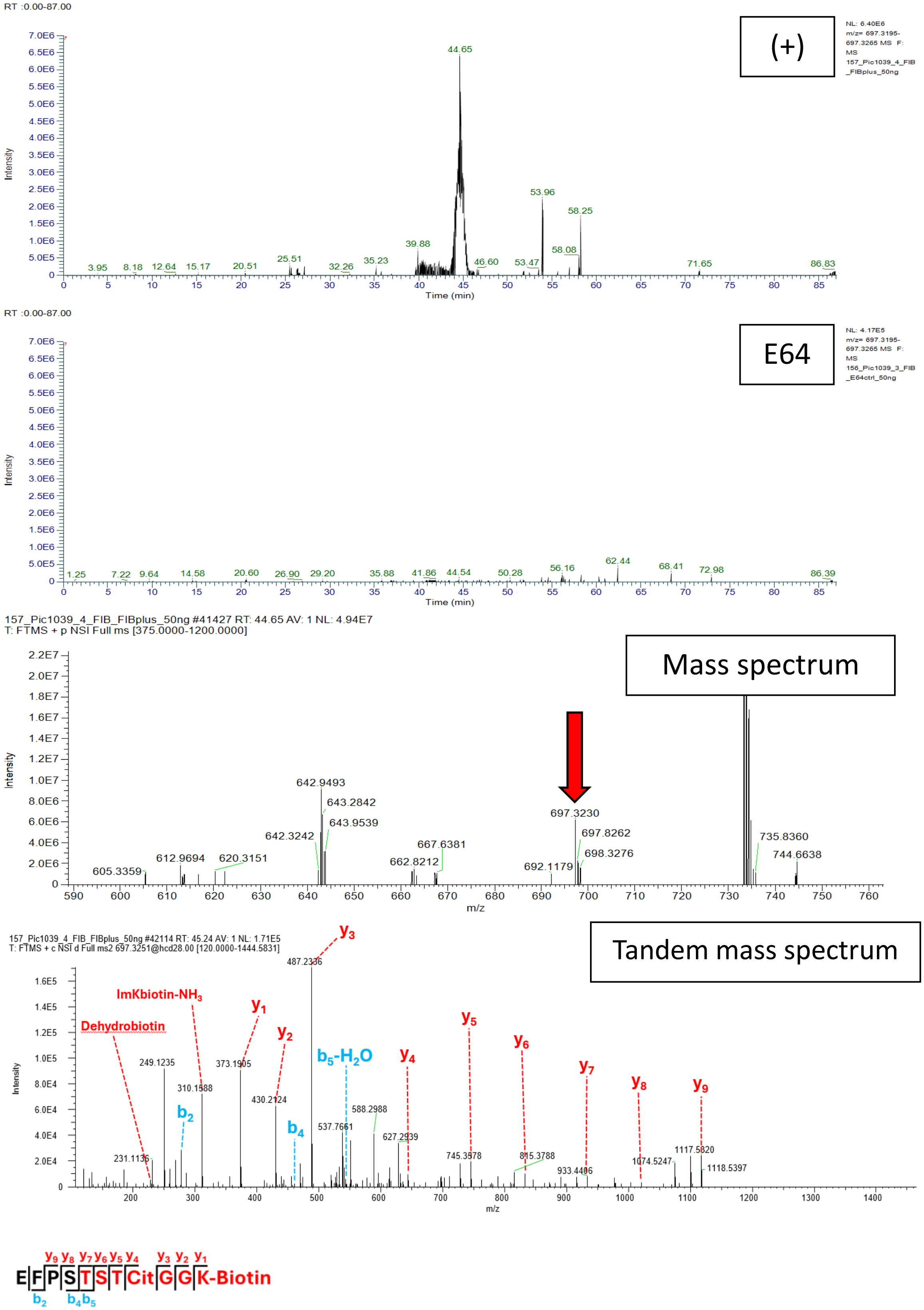


**Figure S18O.** **EFPSTSTCitGGK-Biotin; Scan 42114; RT = 44.66; m/z = 697.3230. Fibrinogen and V1.**

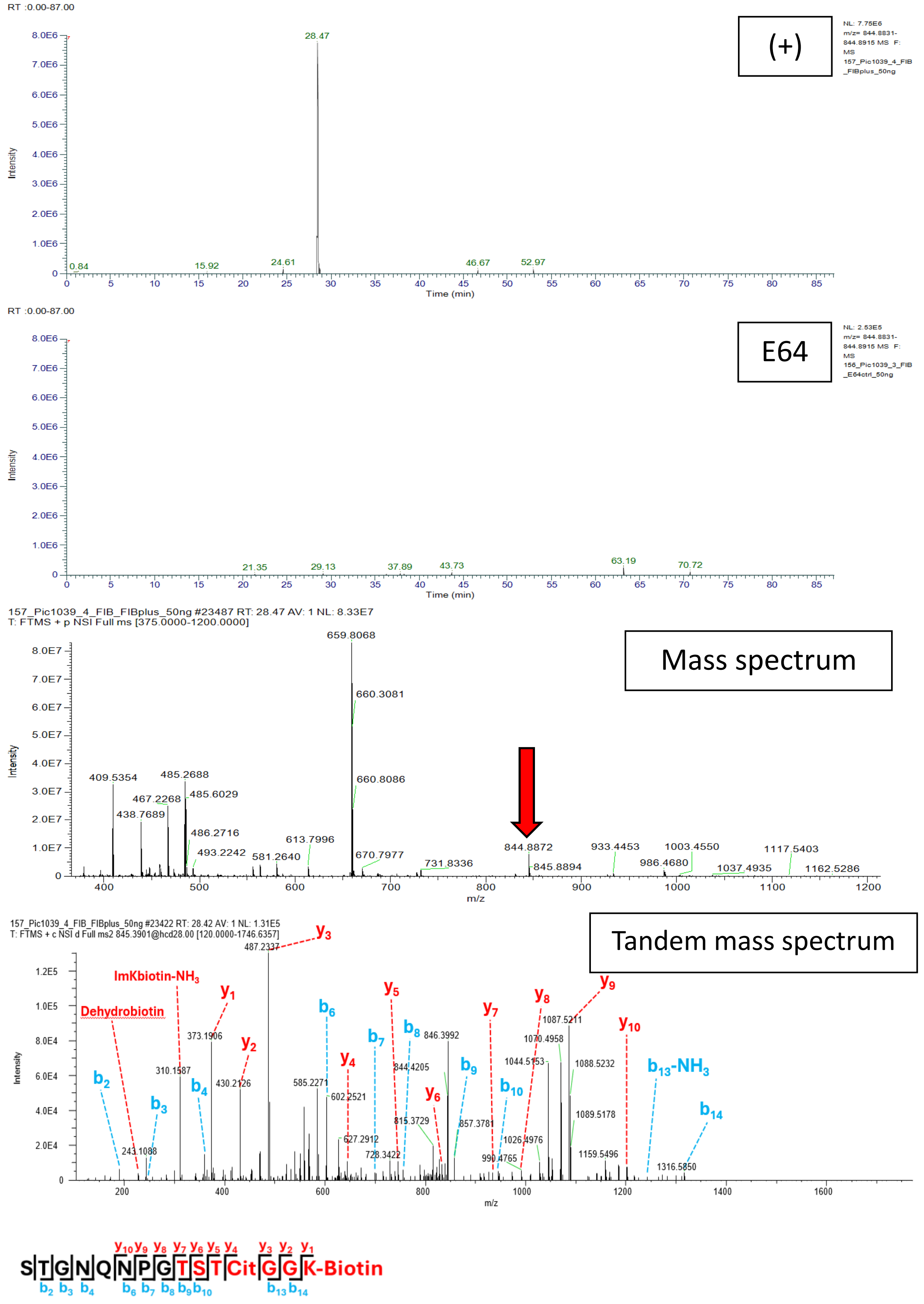


**Figure S18P.** STGNQNPGTSTCitGGK-Biotin; Scan 23422; RT = 28.48; m/z = 844.8873. Fibrinogen and V1.

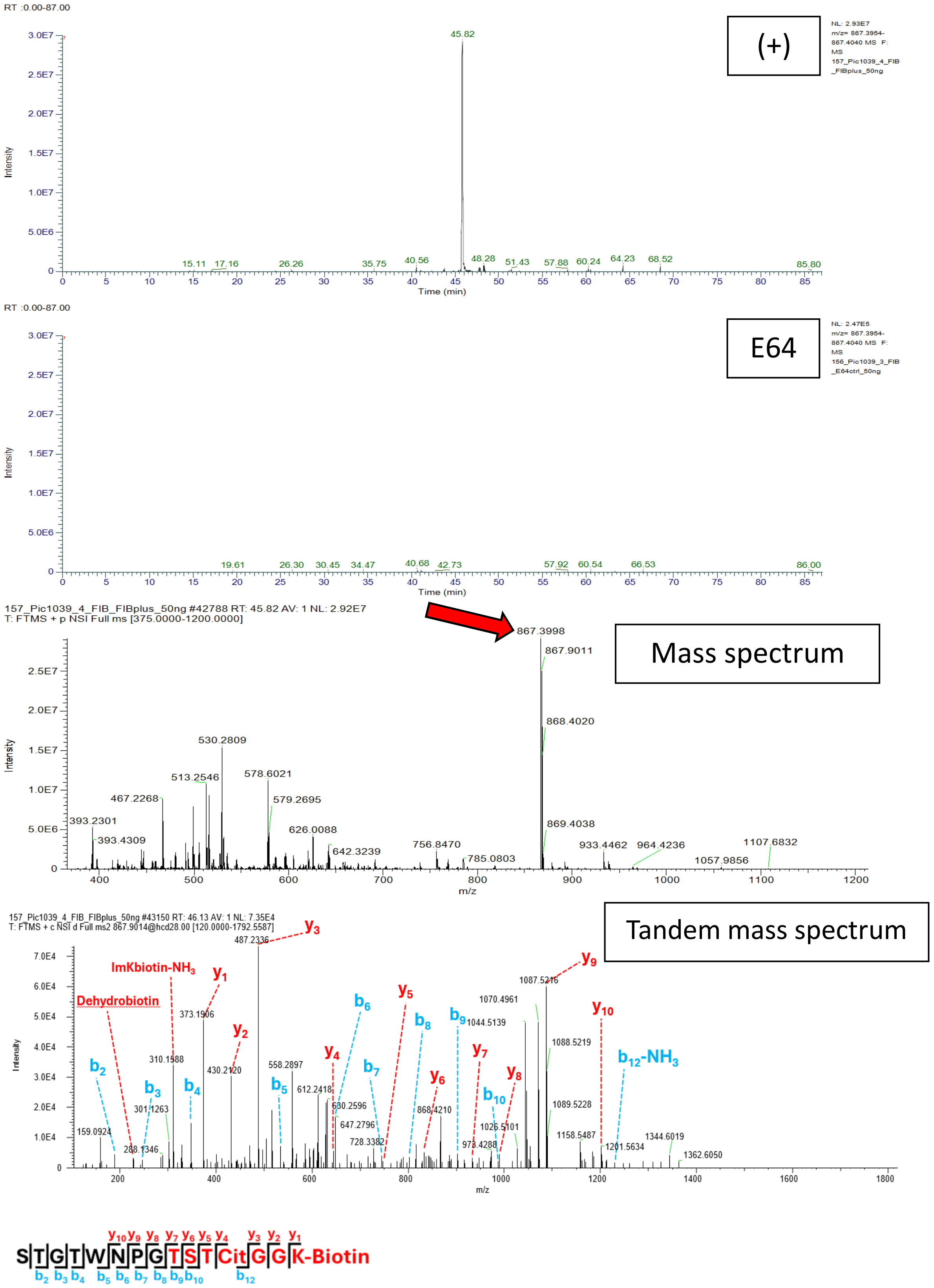


**Figure S18Q. STGTWNPGTSTCitGGK-Biotin; Scan 43150; RT = 45.83; m/z = 867.3997. Fibrinogen and V1.**

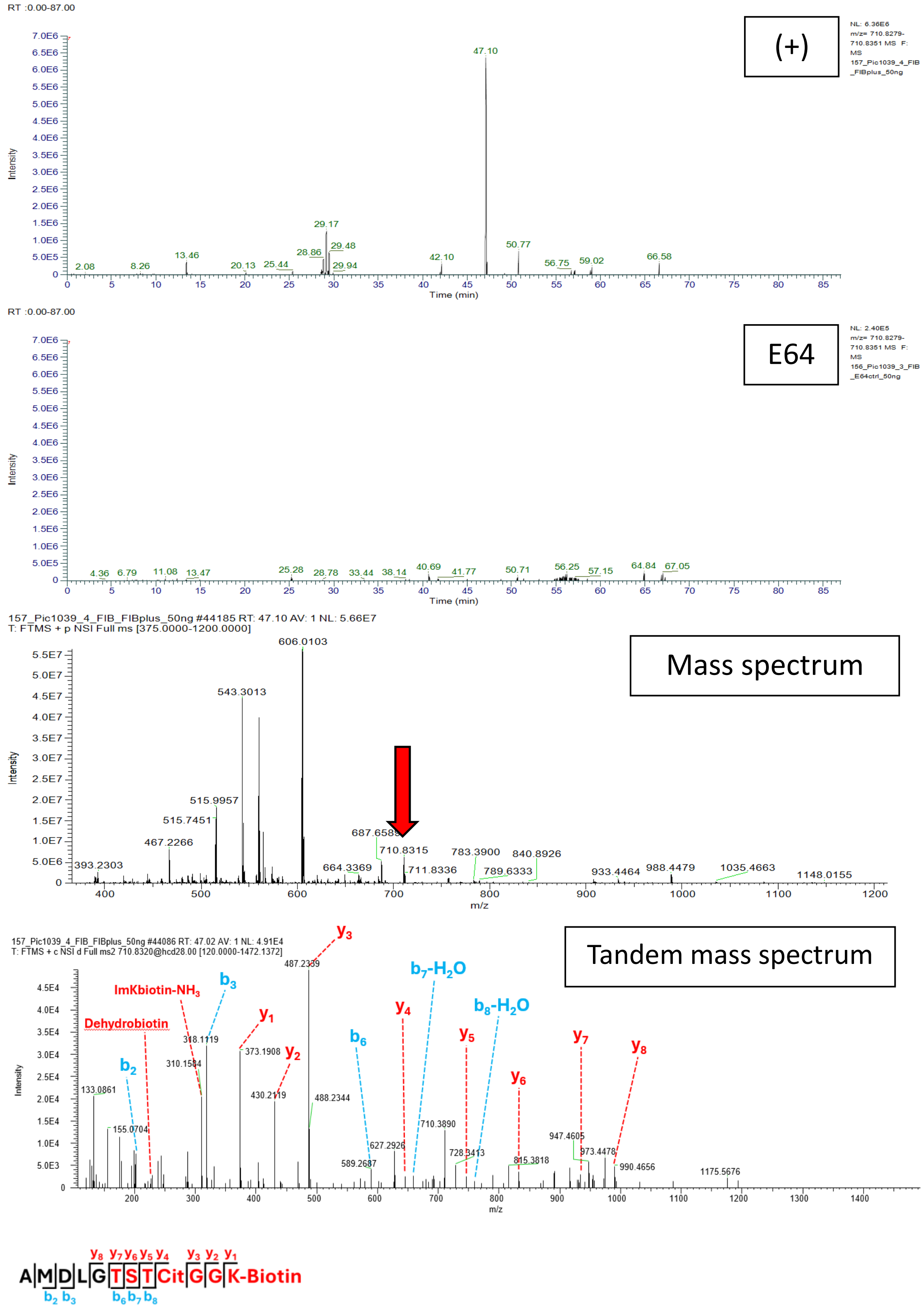


**Figure S18R. AMDLGTSTCitGGK-Biotin; Scan 44086; RT = 47.10; m/z = 710.8315. Fibrinogen and V1.**

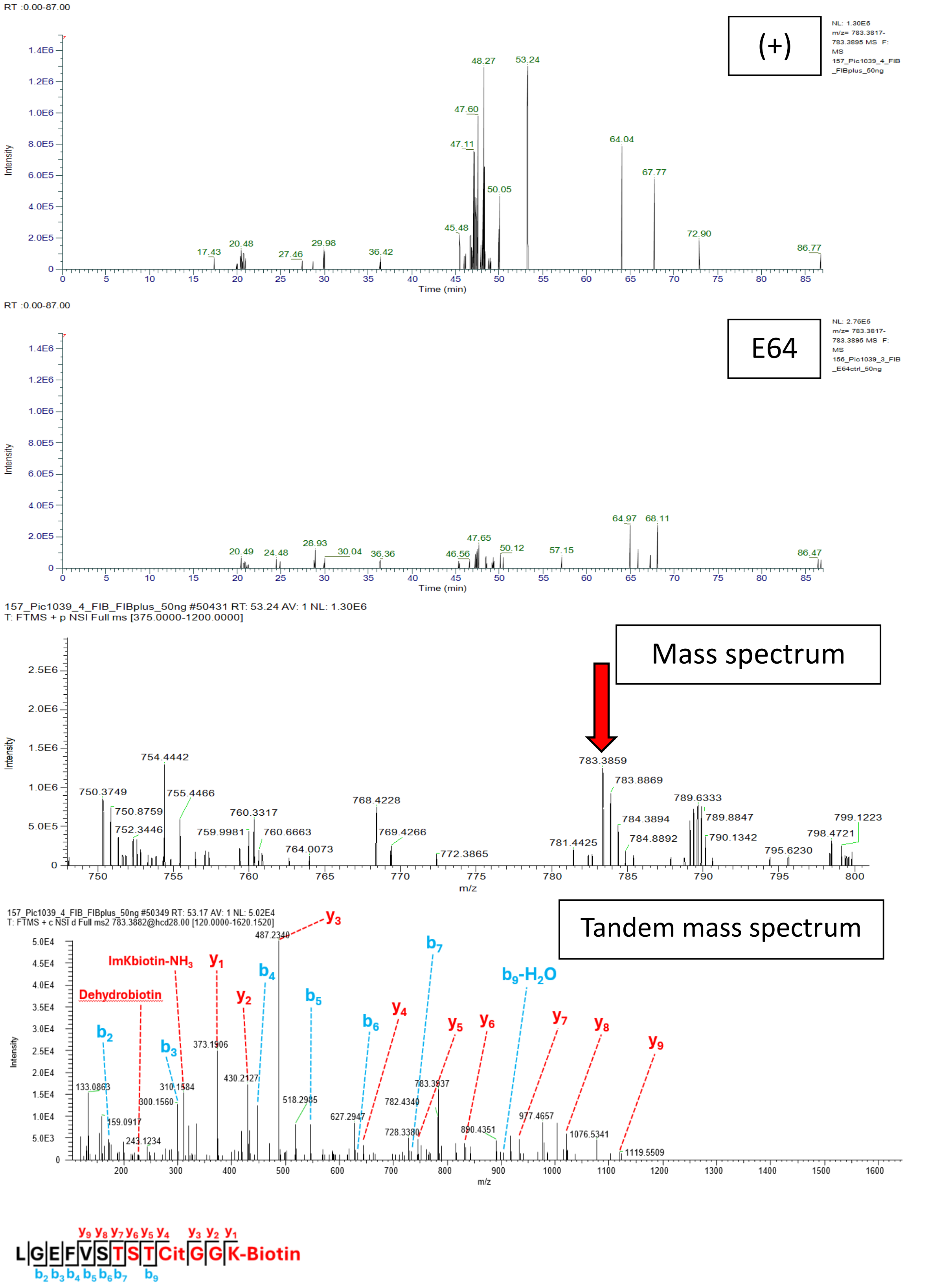


**Figure S18S. LGEFVSTSTCitGGK-Biotin; Scan 50349; RT = 53.23; m/z = 783.3856. Fibrinogen and V1.**

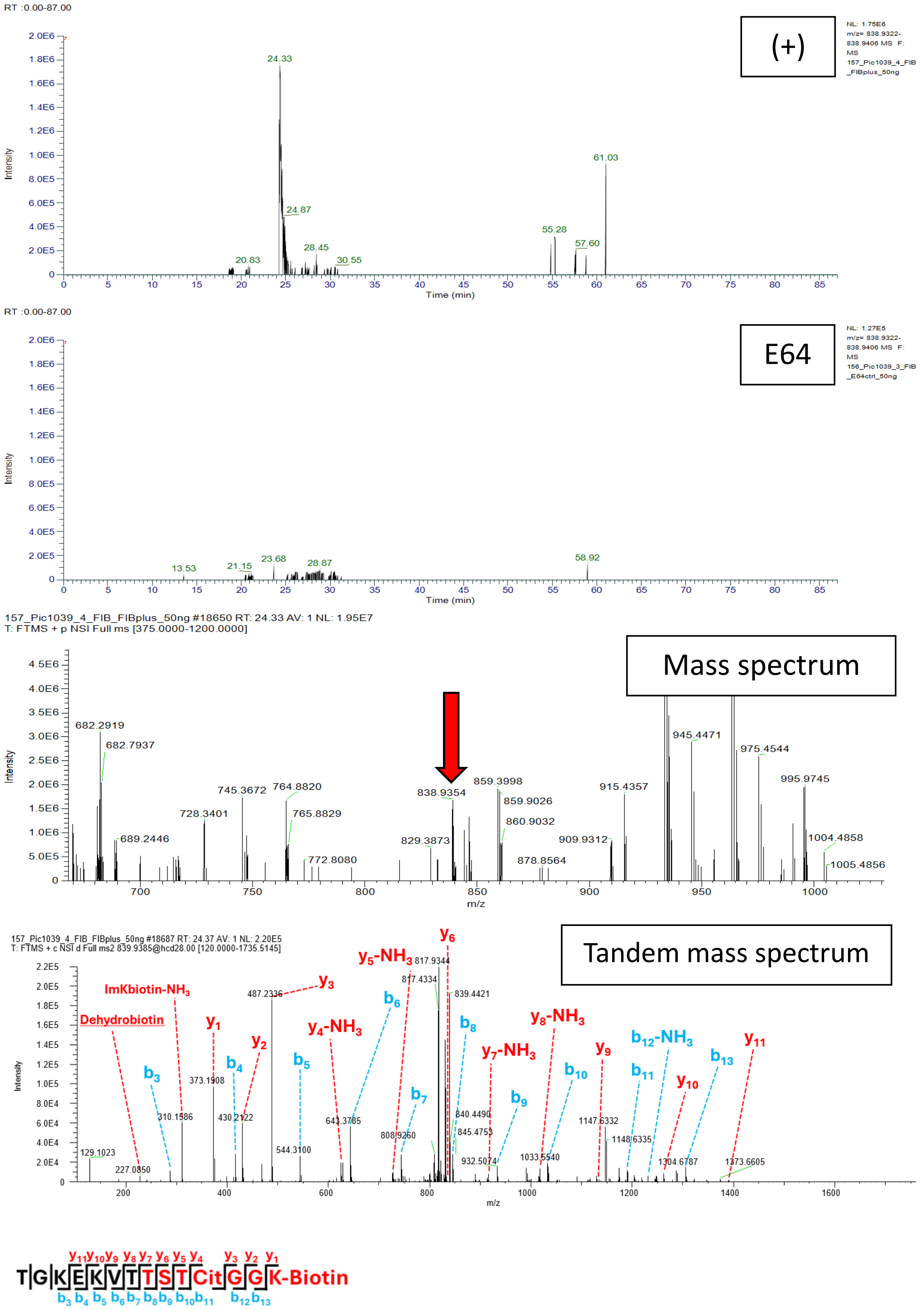


**Figure S18T. TGKEKVTTSTCitGGK-Biotin; Scan 18687; RT = 24.35; m/z = 838.9364. Fibrinogen and V1.**

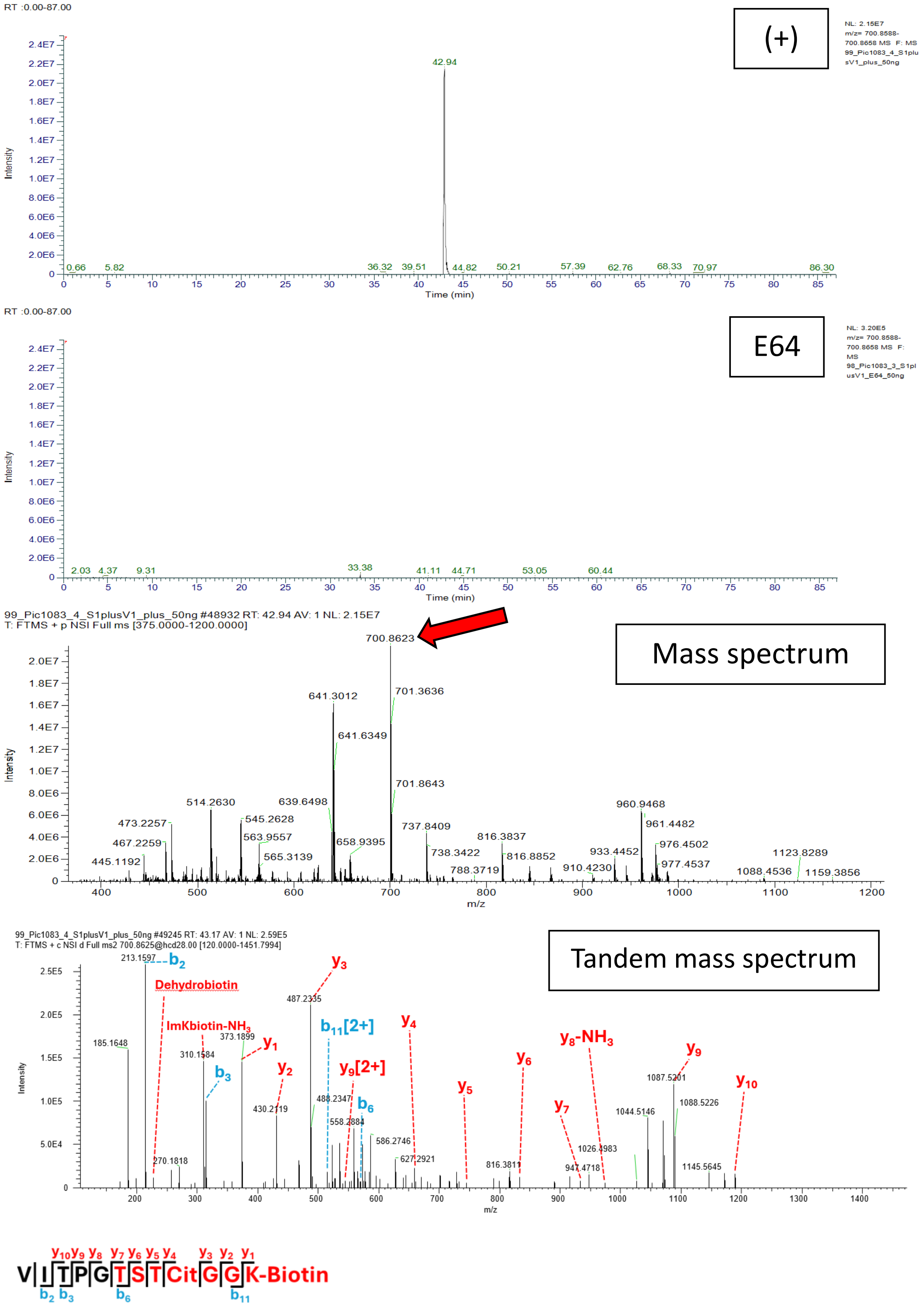


**Figure S18U.** **VITPGTSTCitGGK-Biotin; Scan 49245; RT = 42.92; m/z = 700.8623. Spike and V1.**

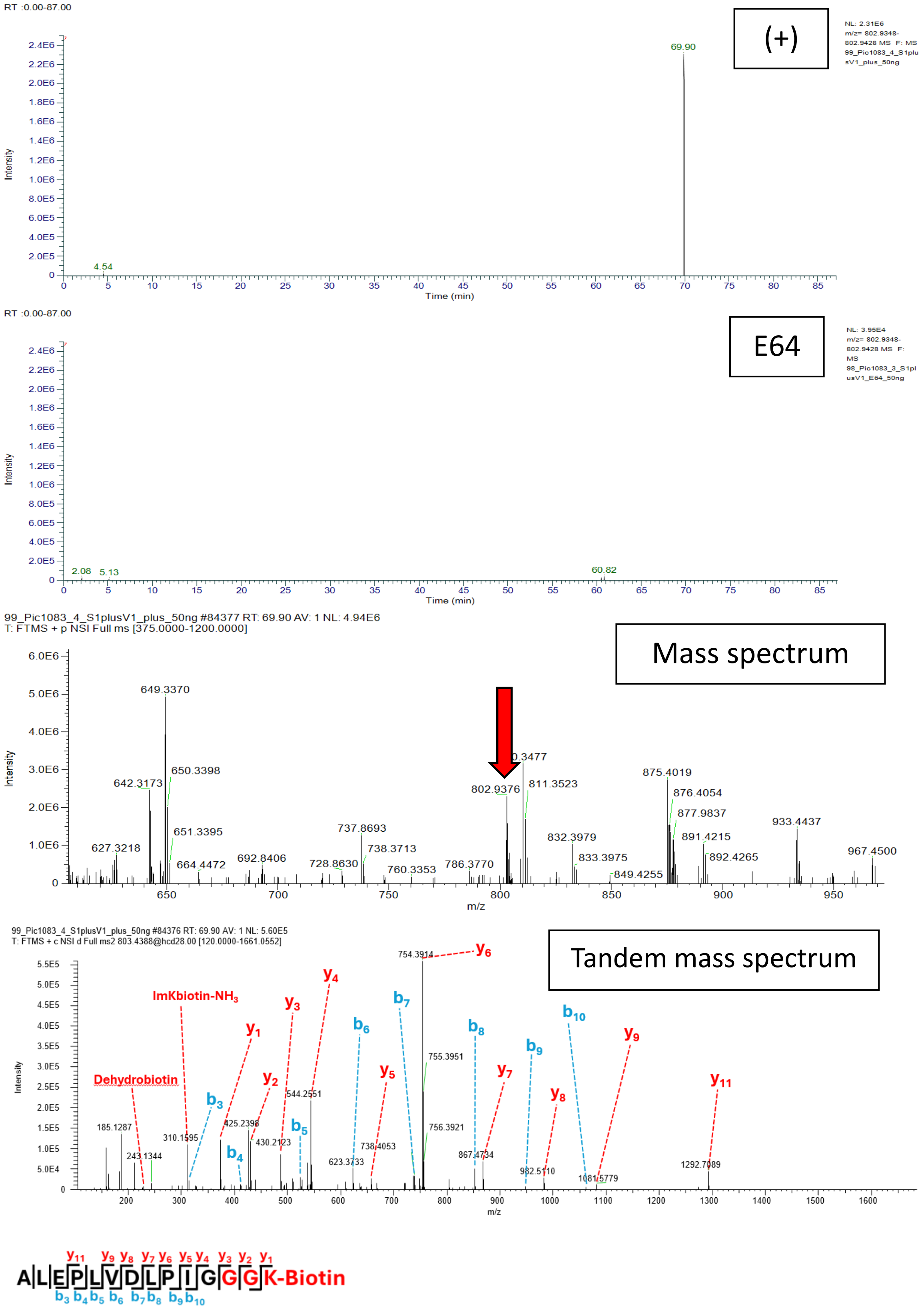


**Figure S18V. ALEPLVDLPIGGGK-Biotin; Scan 84376; RT = 69.91; m/z = 802.9388. Spike and V1.**

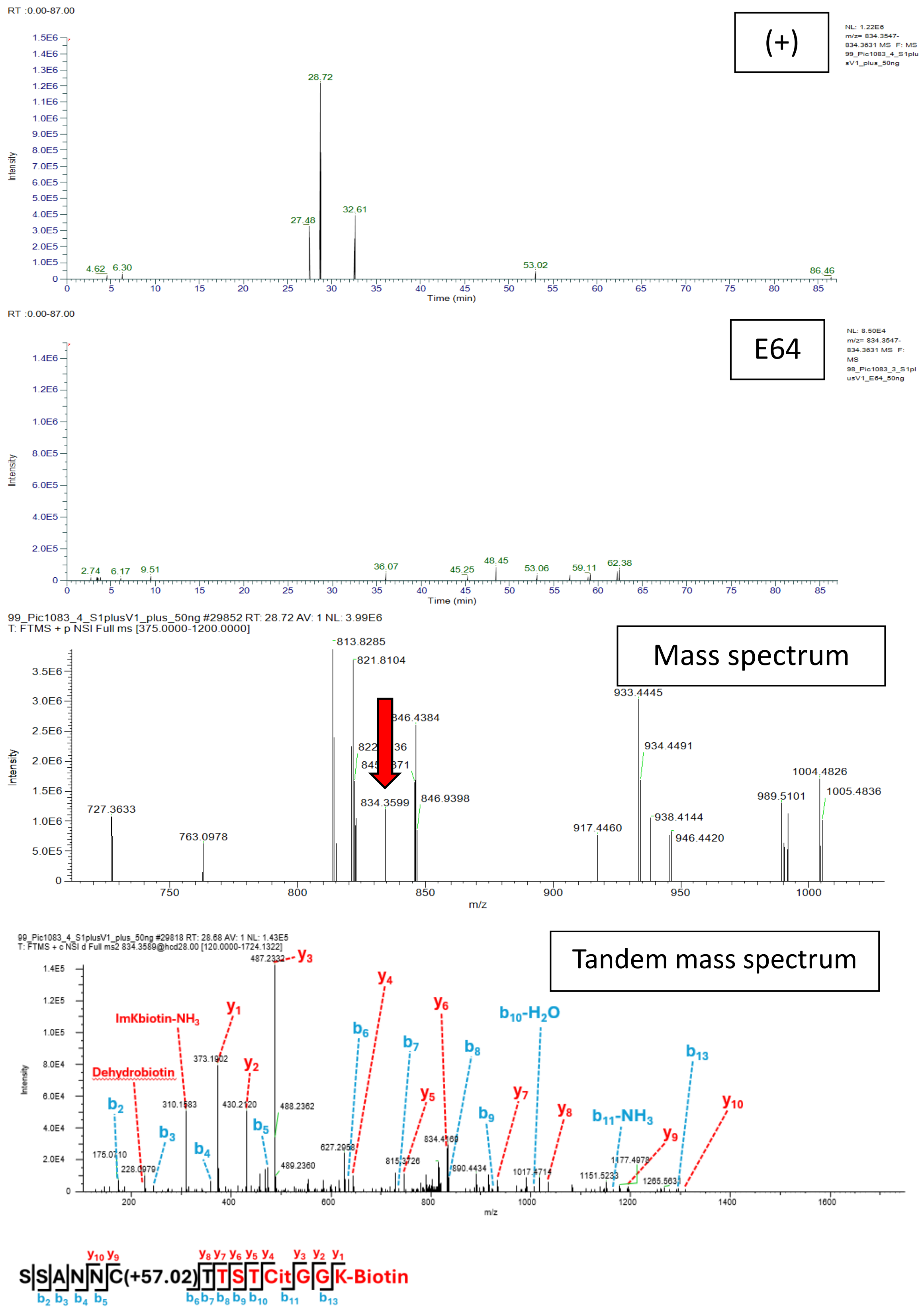


**Figure S18W. SSANNCTTSTCitGGK-Biotin; Scan 29818; RT = 28.68; m/z = 834.3589. Spike and V1.**

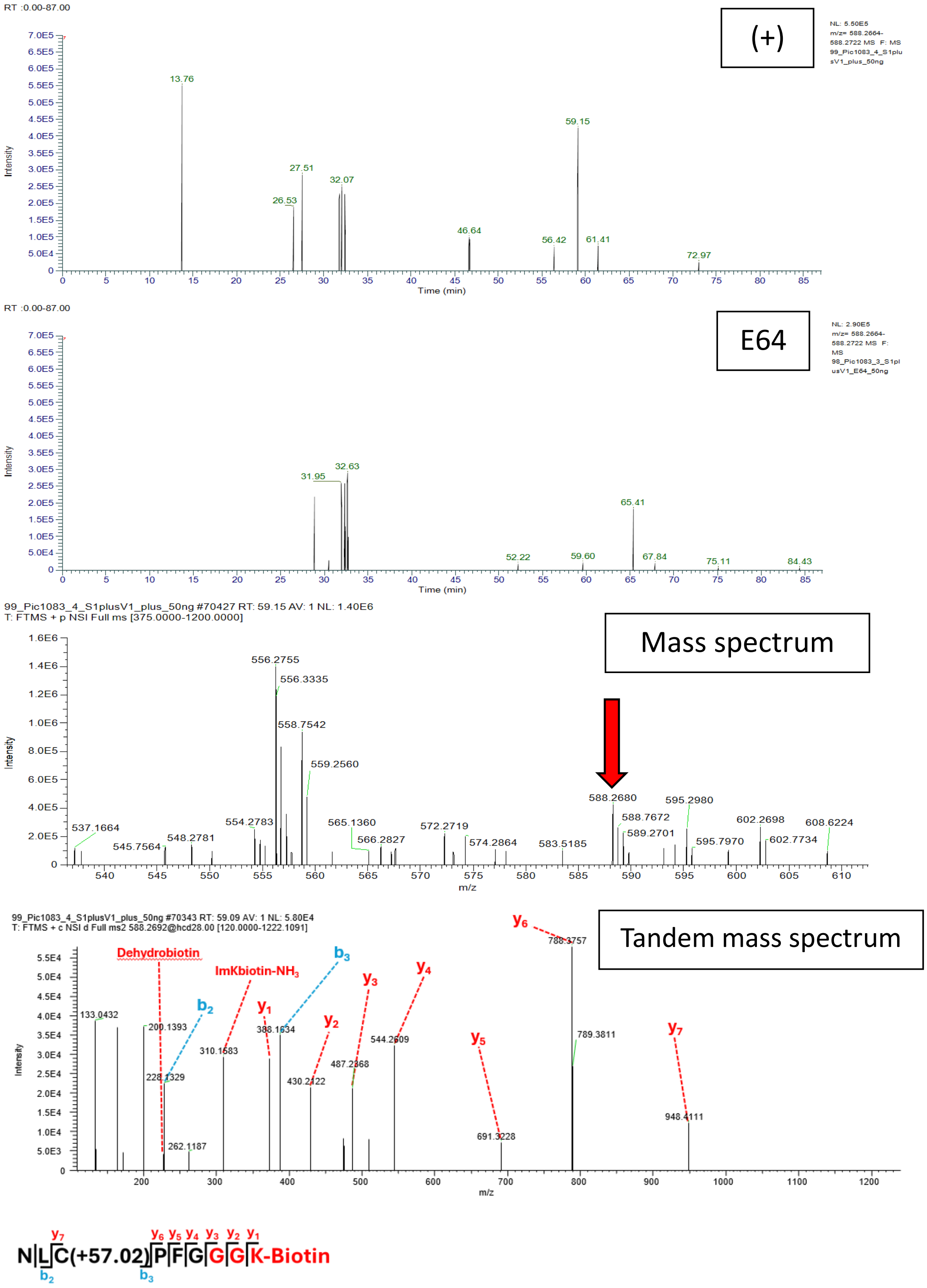


**Figure S18X. NLCPFGGGK-Biotin; Scan 70343; RT = 59.12; m/z = 588.2693. Spike and V1.**

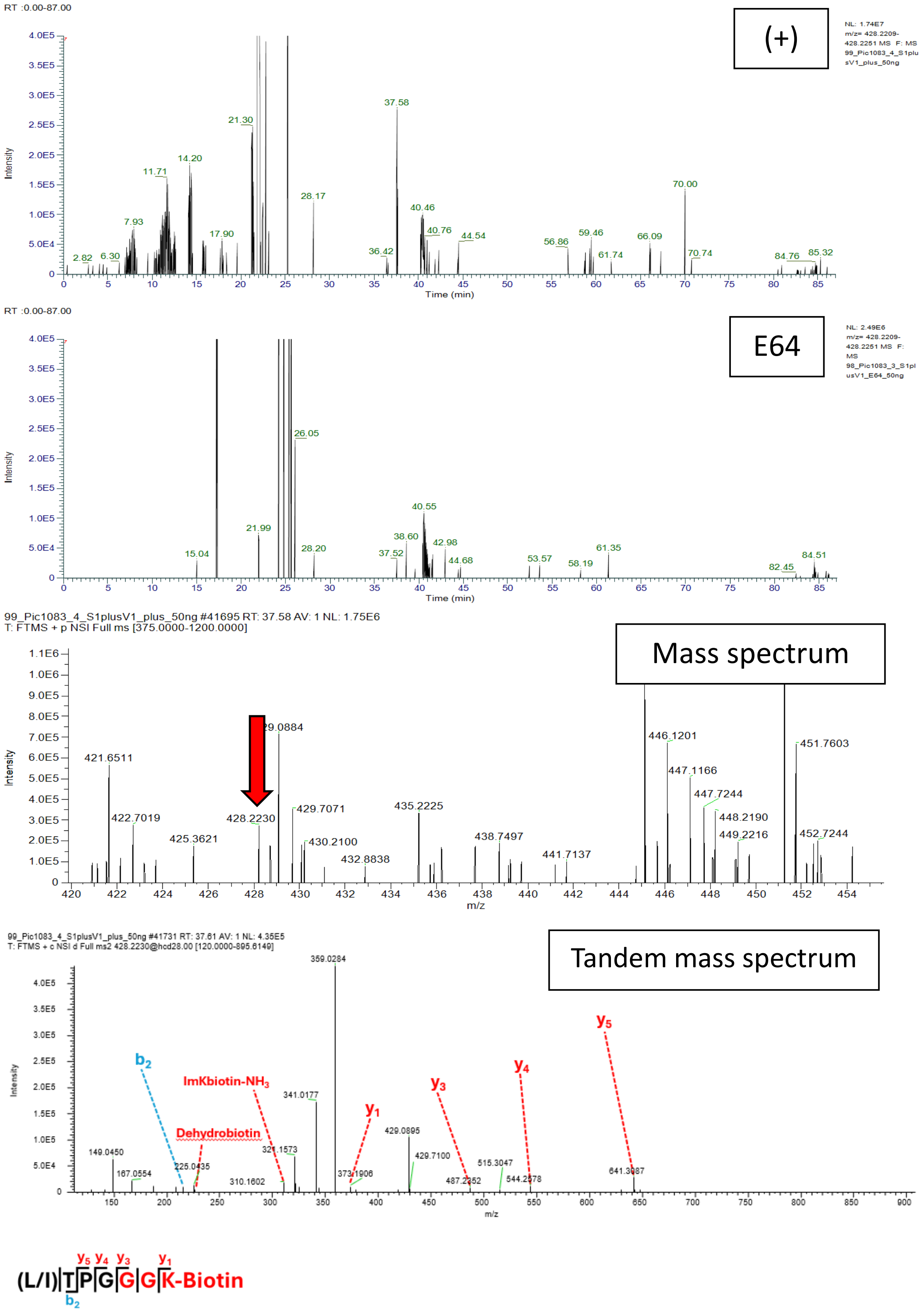


**Figure S18Y. (L/I)TPGGGK-Biotin; Scan 41731; RT = 37.61; m/z = 428.2230. Spike and V1.**

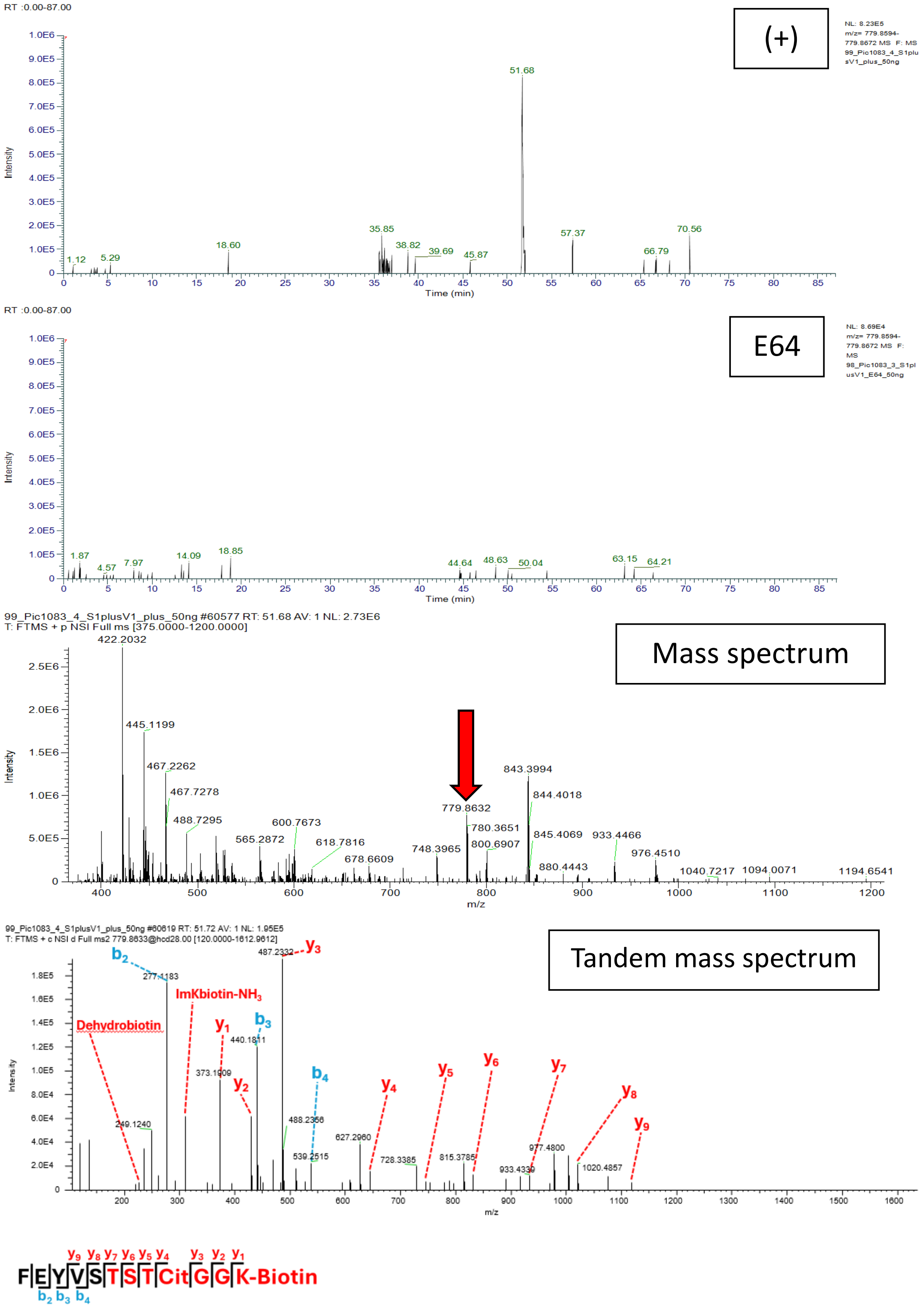


**Figure S18Z. FEYVSTSTCitGGK-Biotin; Scan 60619; RT = 51.70; m/z = 779.8633. Spike and V1.**

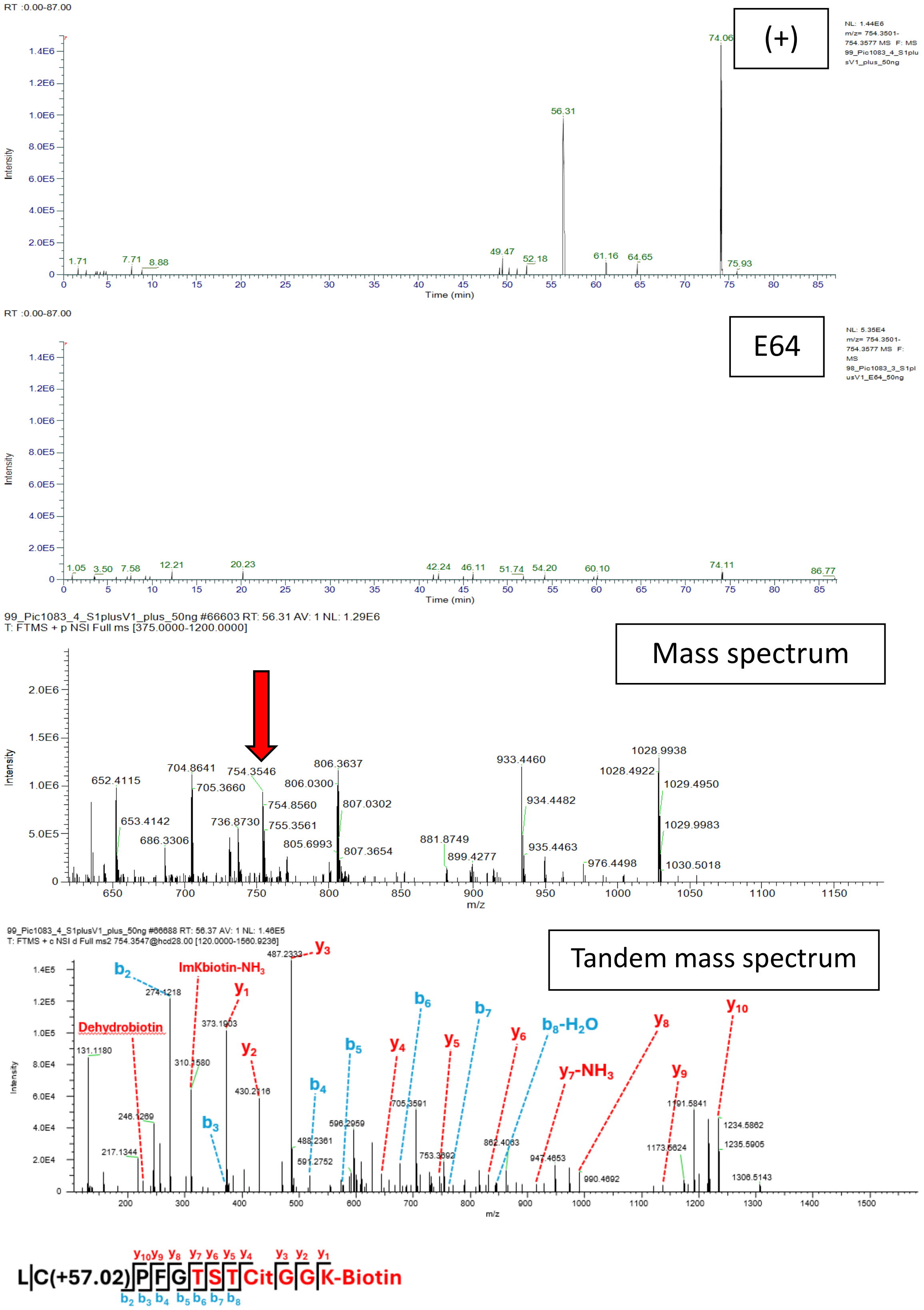


**Figure S18AA. LCPFGTSTCitGGK-Biotin; Scan 66688; RT = 56.34; m/z = 754.3539. Spike and V1.**

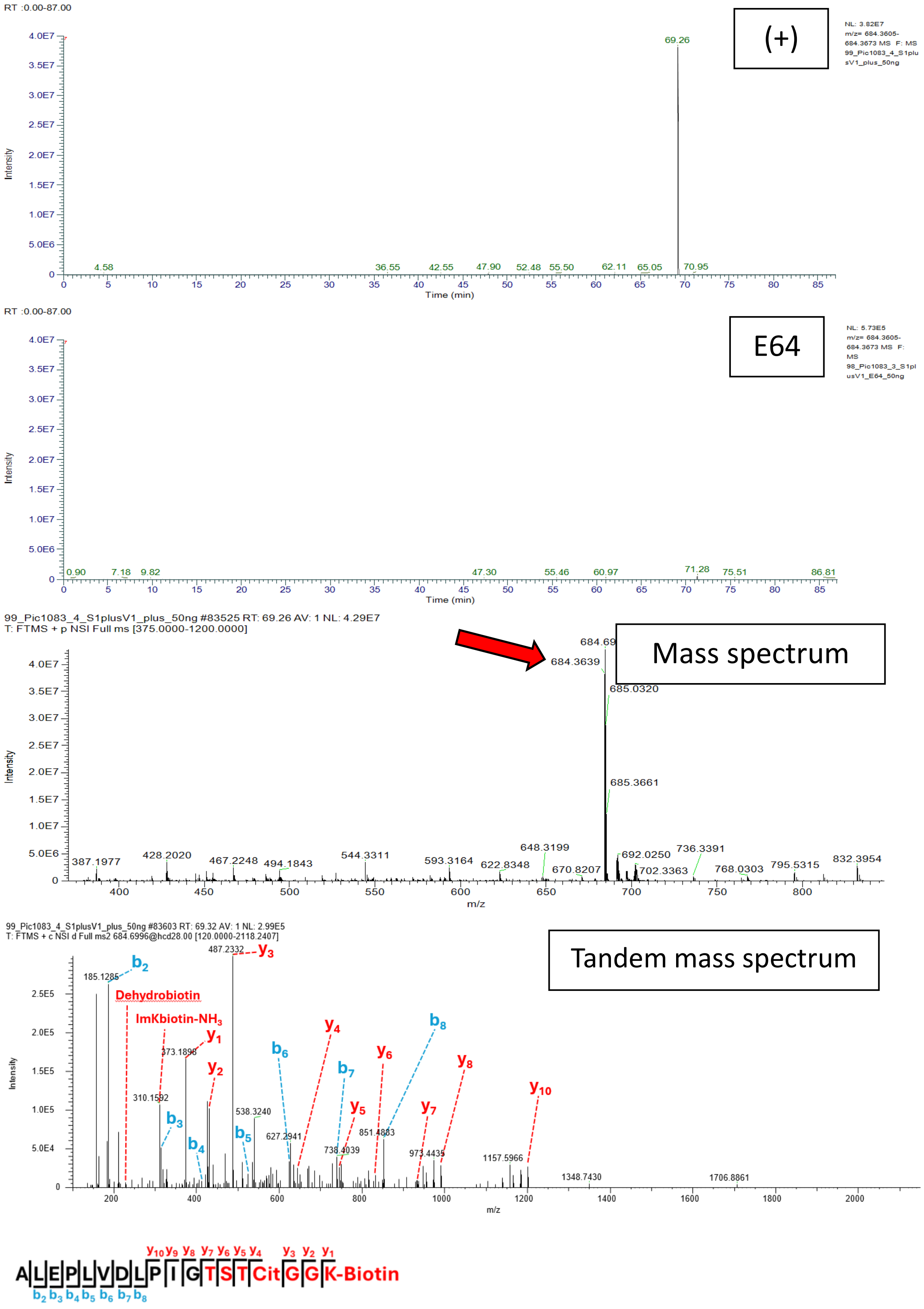


**Figure S18AB. ALEPLVDLPIGTSTCitGGK-Biotin; Scan 83603; RT = 69.26; m/z = 684.3639. Spike and V1.**

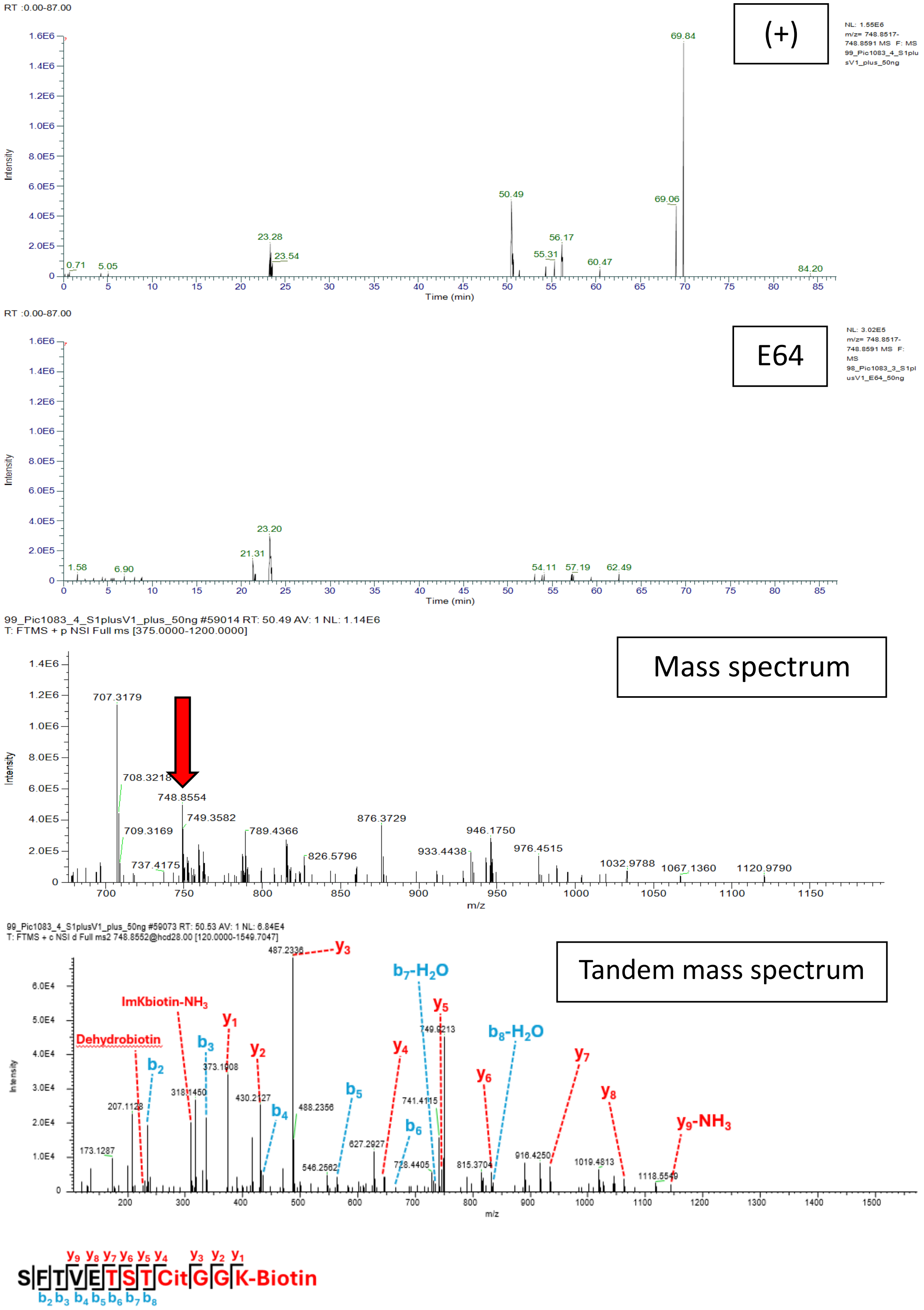


**Figure S18AC. SFTVETSTCitGGK-Biotin; Scan 59073; RT = 50.49; m/z = 748.8554. Spike and V1.**

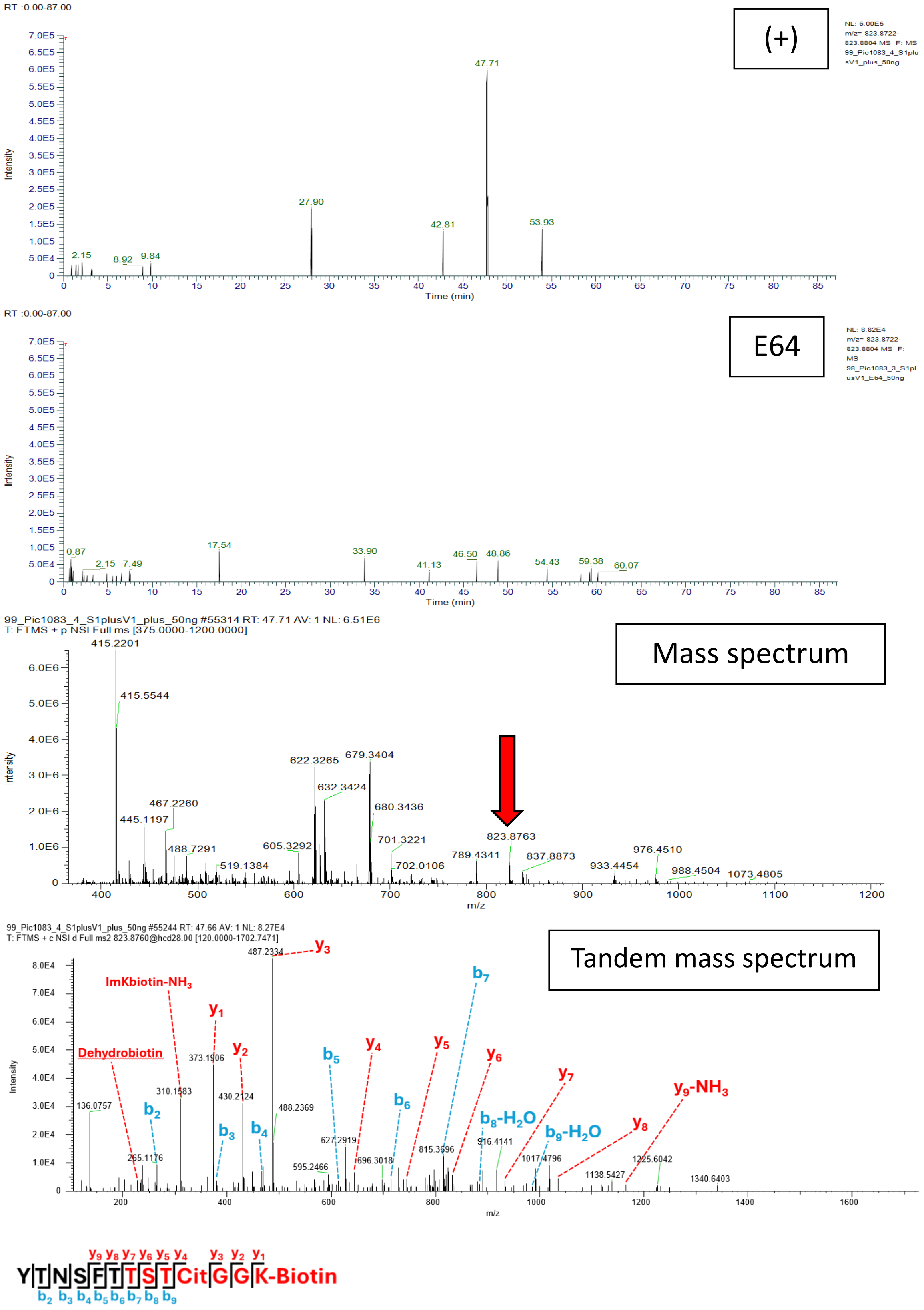


**Figure S18AD. YTNSFTTSTCitGGK-Biotin; Scan 55244; RT = 47.71; m/z = 823.8763. Spike and V1.**

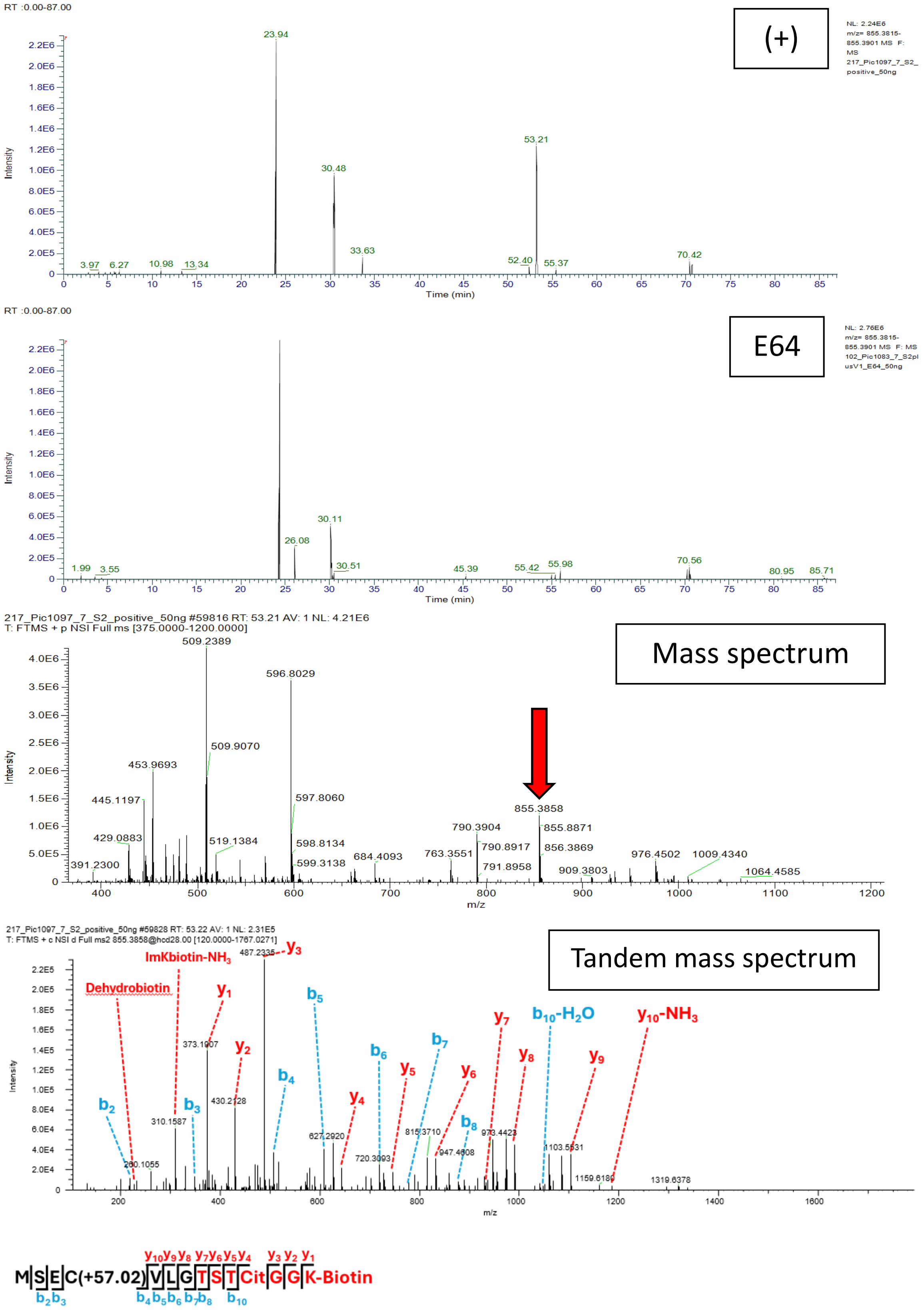


**Figure S18AE. MSECVLGTSTCitGGK-Biotin; Scan 59828; RT = 53.22; m/z = 855.3858. Spike and V1.**

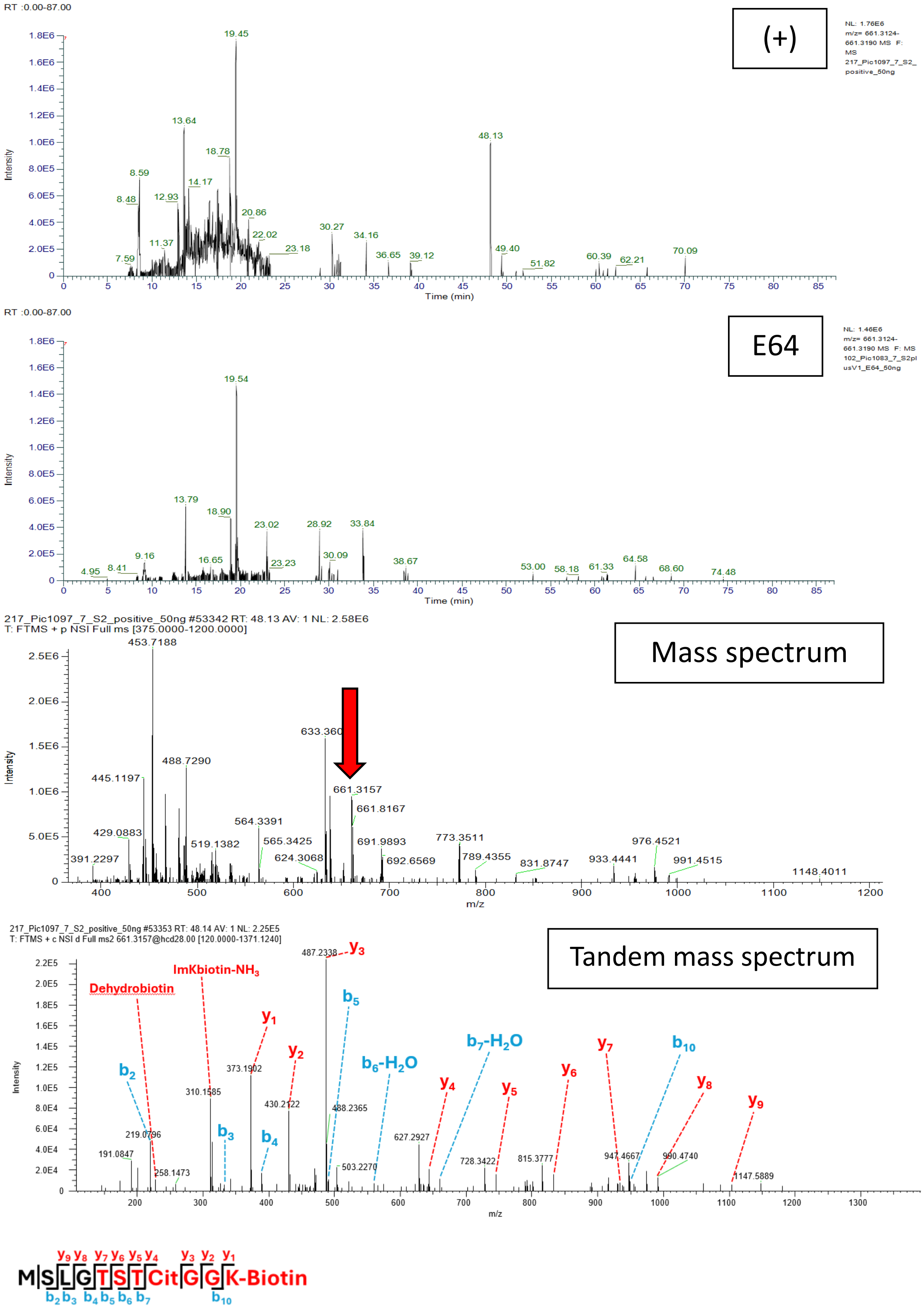


**Figure S18AF. MSLGTSTCitGGK-Biotin; Scan 53353; RT = 48.12; m/z = 661.3157. Spike and V1.**

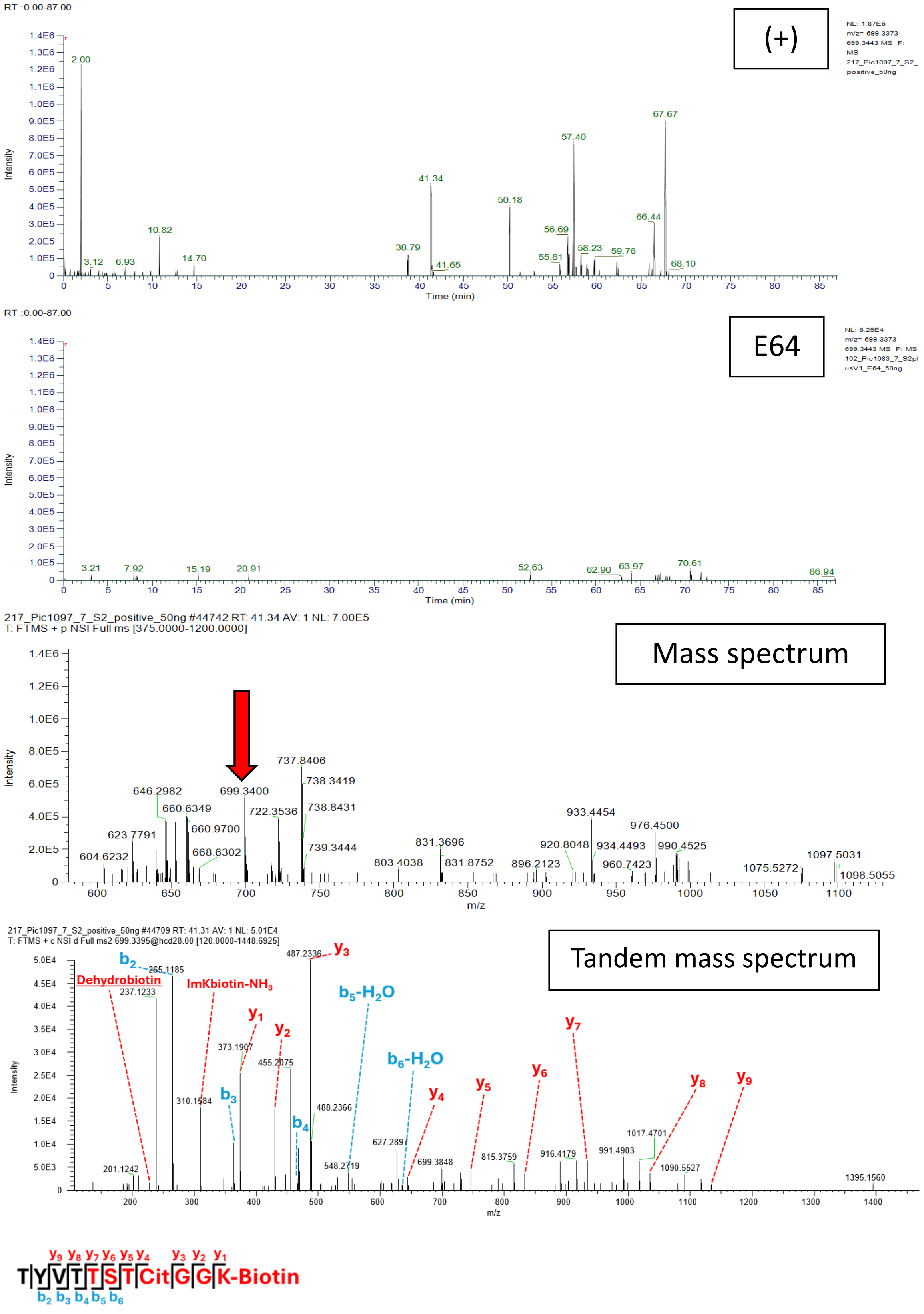


**Figure S18AG. TYVTTSTCitGGK-Biotin; Scan 44709; RT = 41.36; m/z = 699.3408. Spike and V1.**

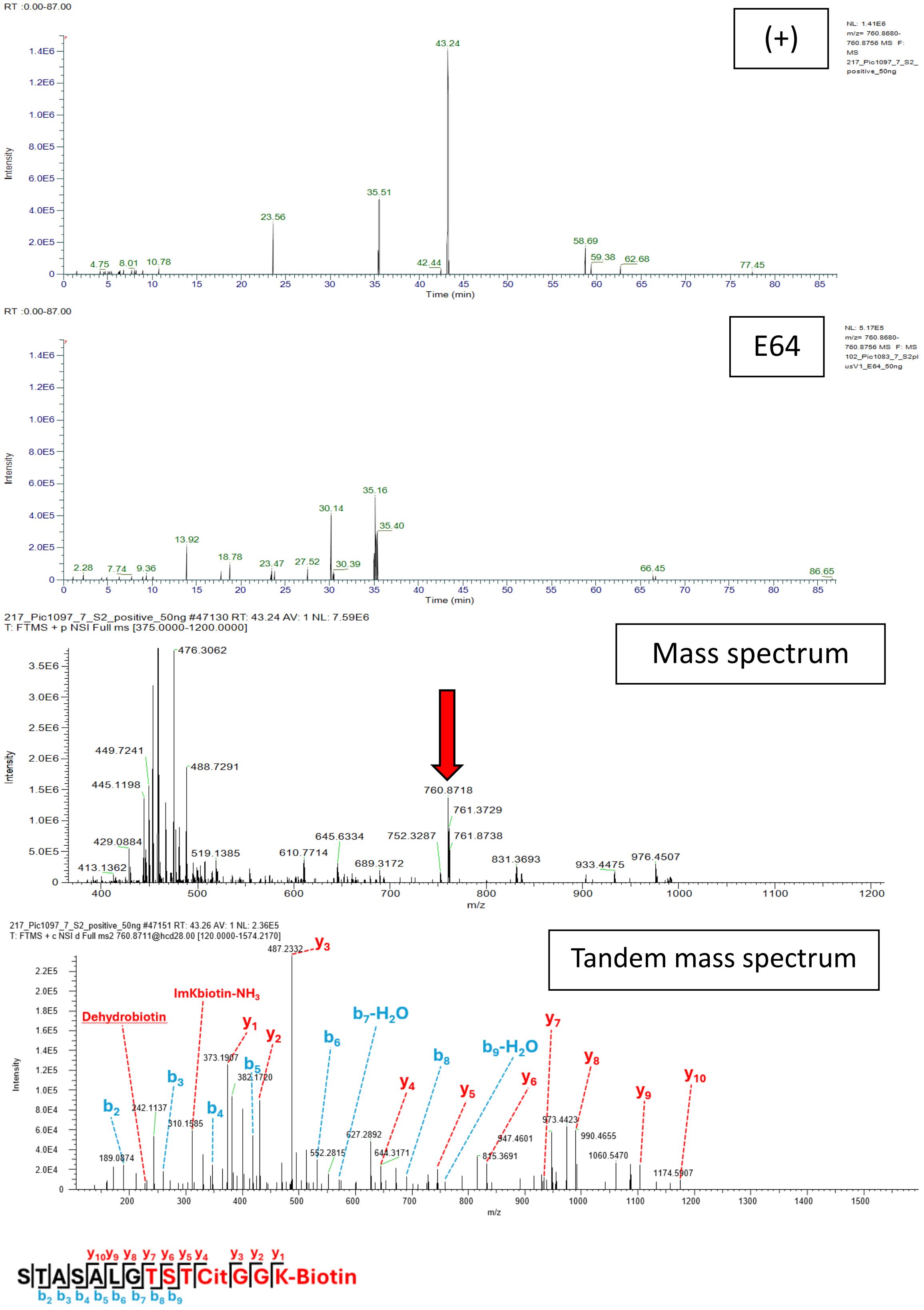


**Figure S18AH. STASALGTSTCitGGK-Biotin; Scan 47151; RT = 43.24; m/z = 760.8718. Spike and V1.**

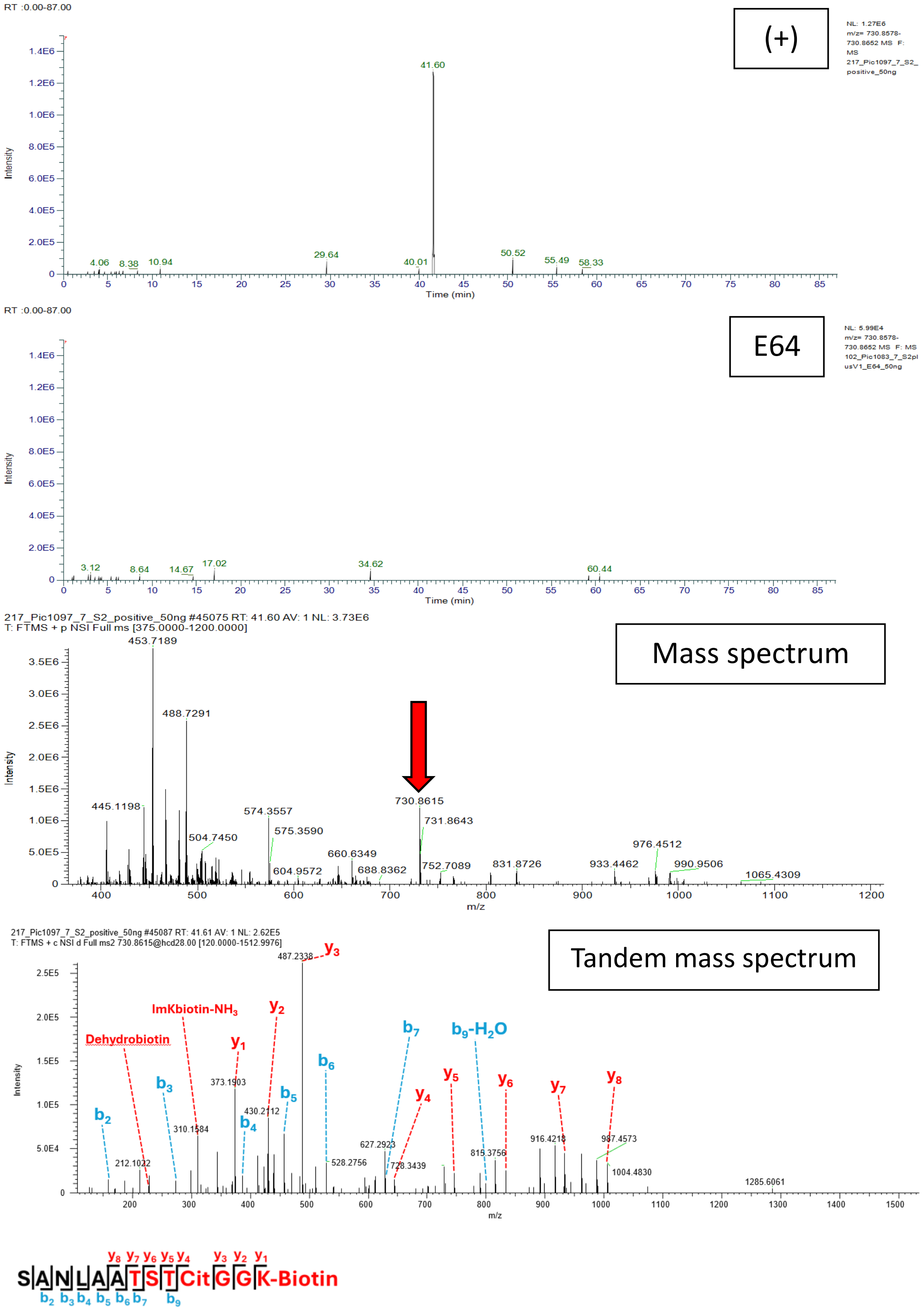


**Figure S18AI. SANLAATSTCitGGK-Biotin; Scan 45087; RT = 41.60; m/z = 730.8615. Spike and V1.**

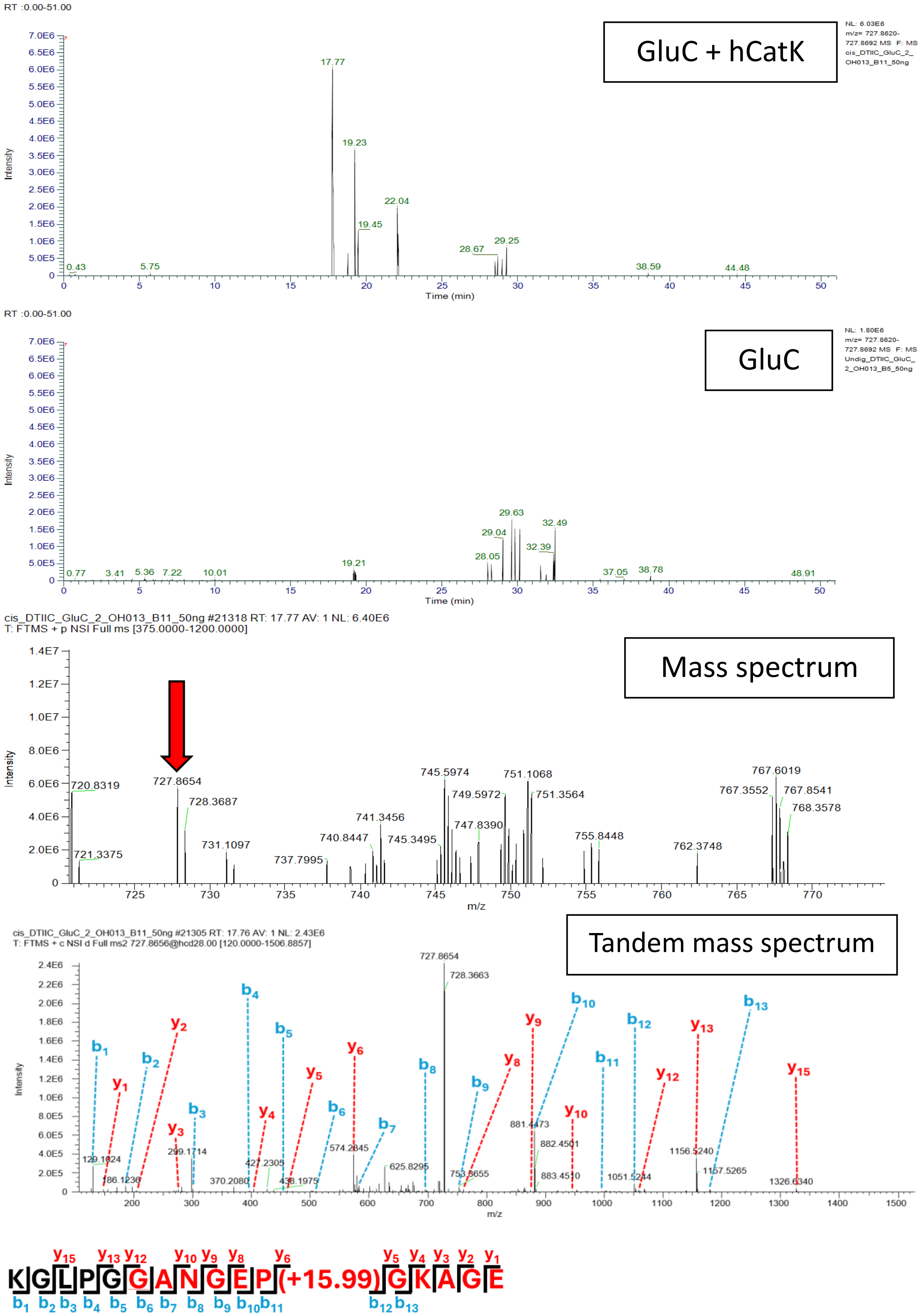


**Figure S19A.** **KGLPGGANGEPGKAGE; Scan 21305; RT = 17.76; m/z = 727.8656. DTIIC *in cis*.**

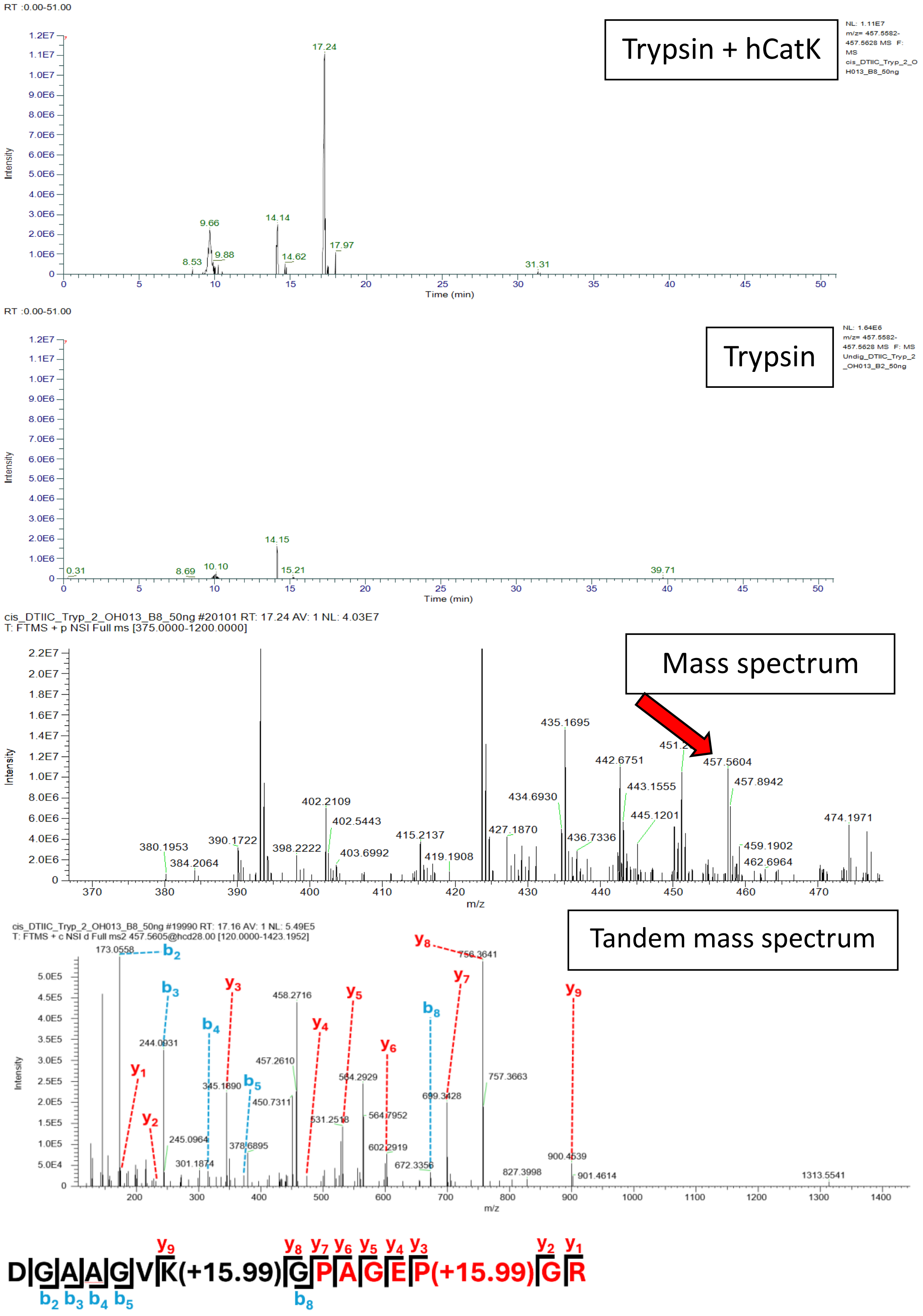


**Figure S19B. DGAAGVKGPAGEPGR; Scan 19990; RT = 17.16; m/z = 457.5605. DTIIC *in cis*.**

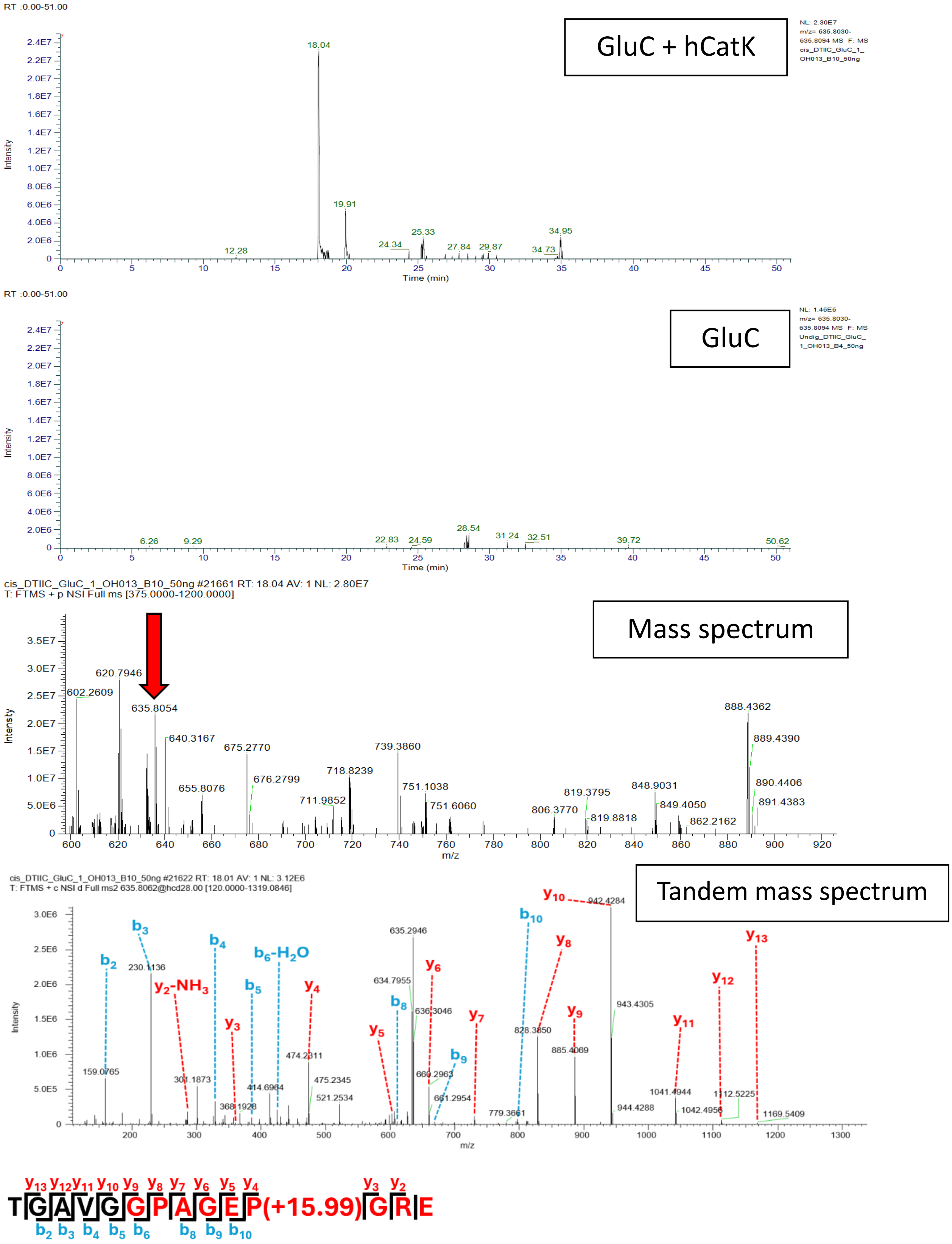


**Figure S19C. TGAVGGPAGEPGRE; Scan 21622; RT = 18.01; m/z = 635.8062. DTIIC *in cis*.**

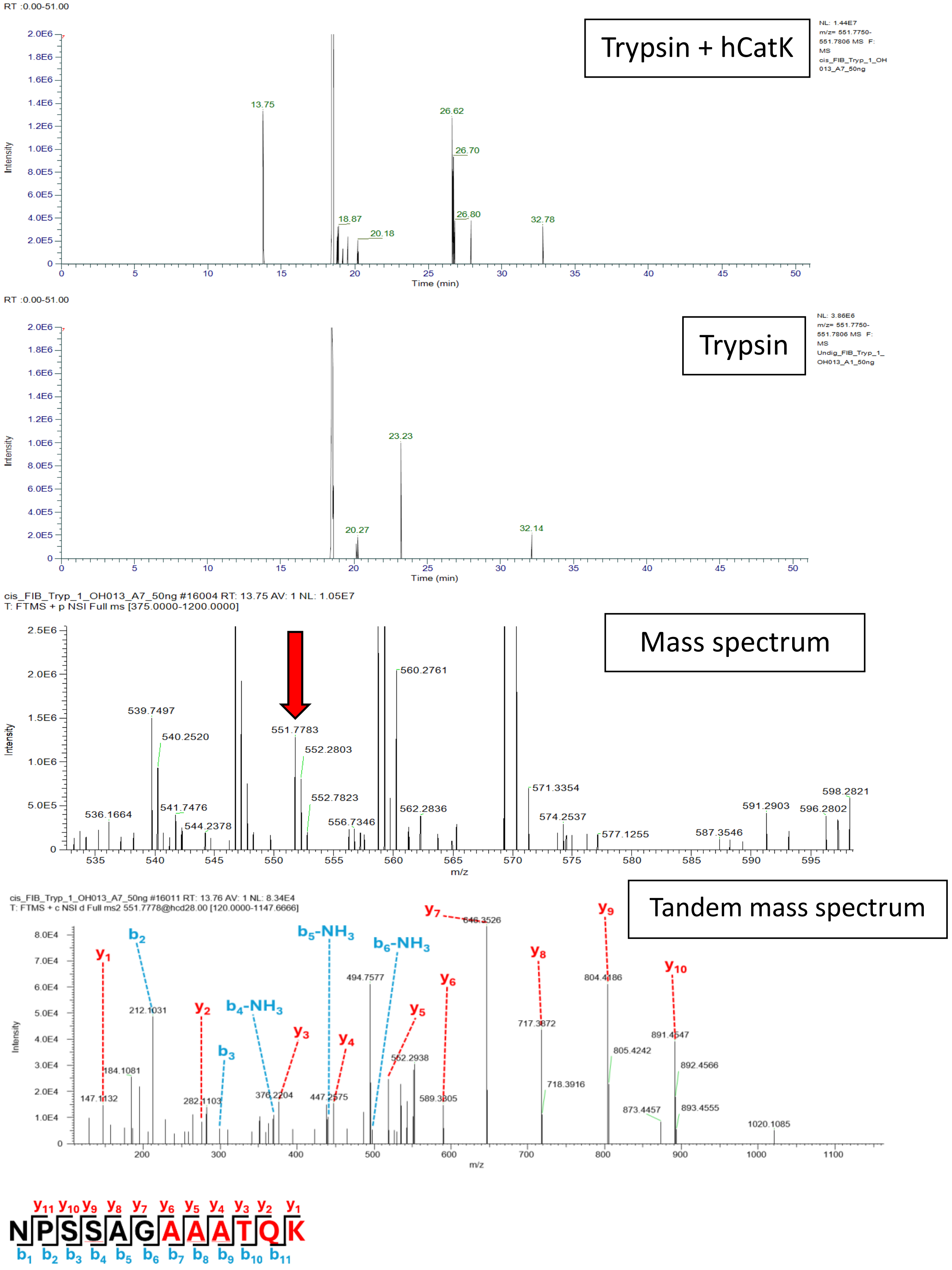


**Figure S19D. NPSSAGAAATQK; Scan 16011; RT = 13.76; m/z = 551.7778. Fibrinogen *in cis*.**

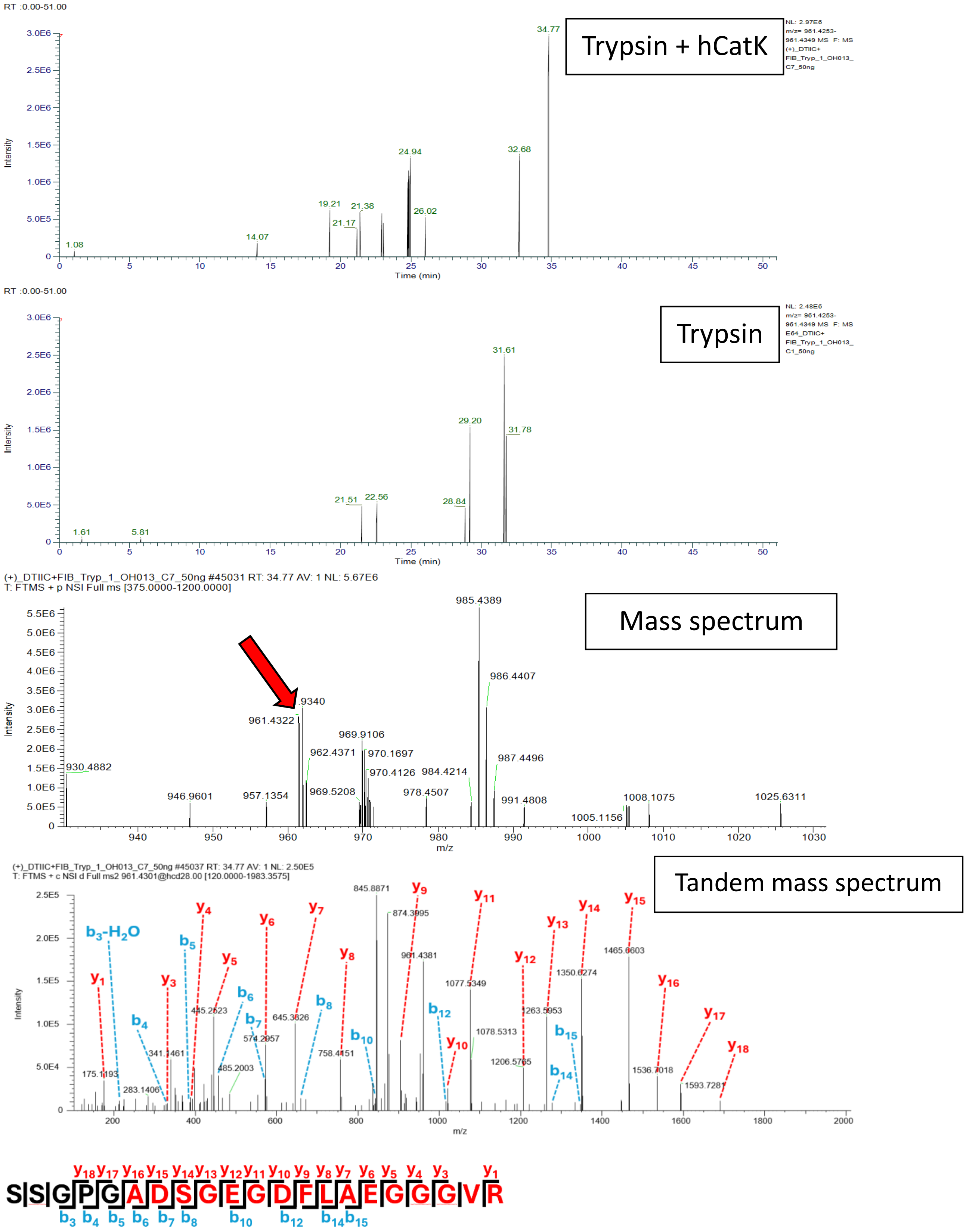


**Figure S19E. SSGPGADSGEGDFLAEGGGVR; Scan 45037; RT = 34.77; m/z = 961.4301. Fibrinogen *in cis*.**

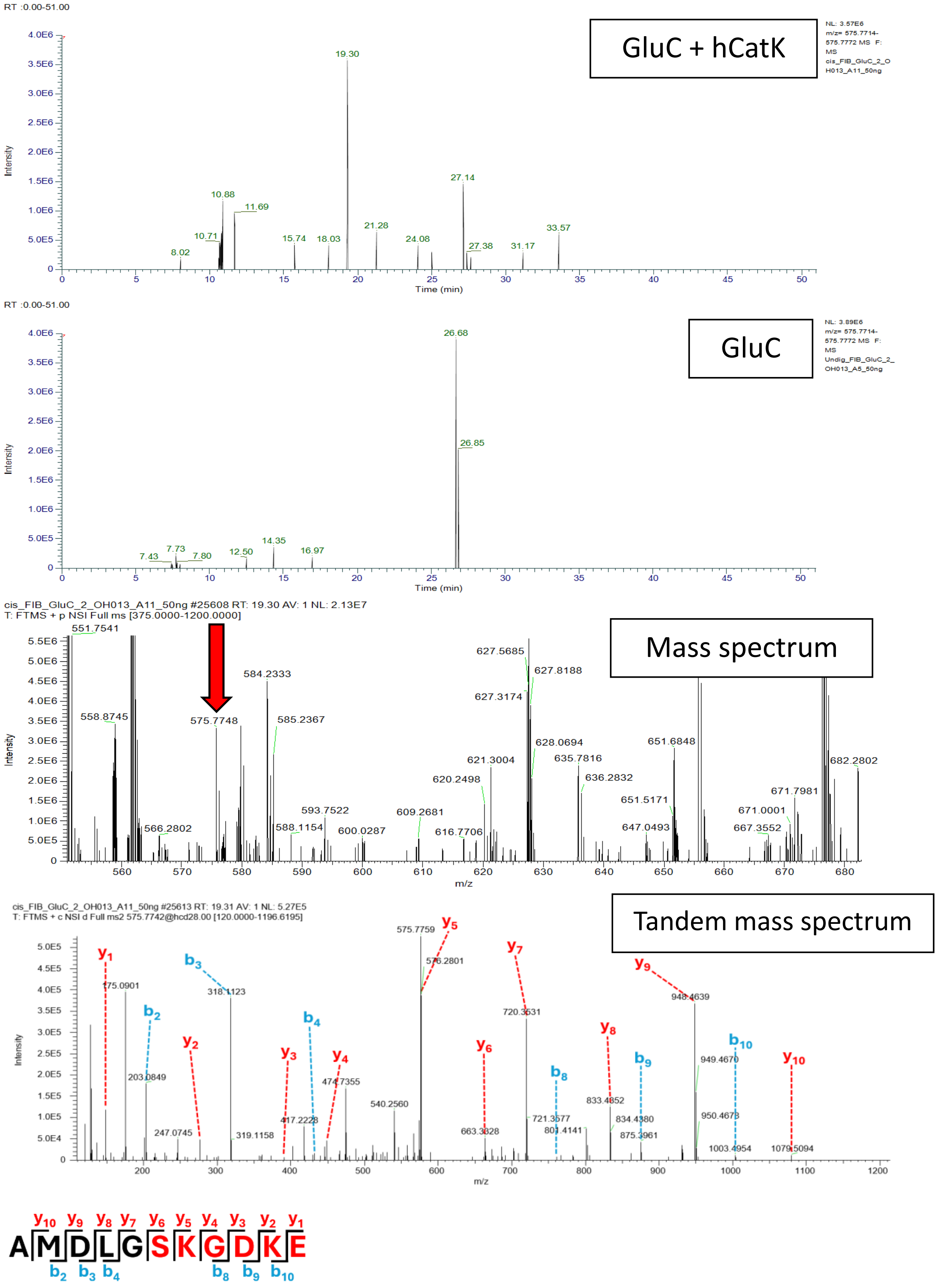


**Figure S19F. AMDLGSKGDKE; Scan 25613; RT = 19.31; m/z = 575.7743. Fibrinogen *in cis*.**

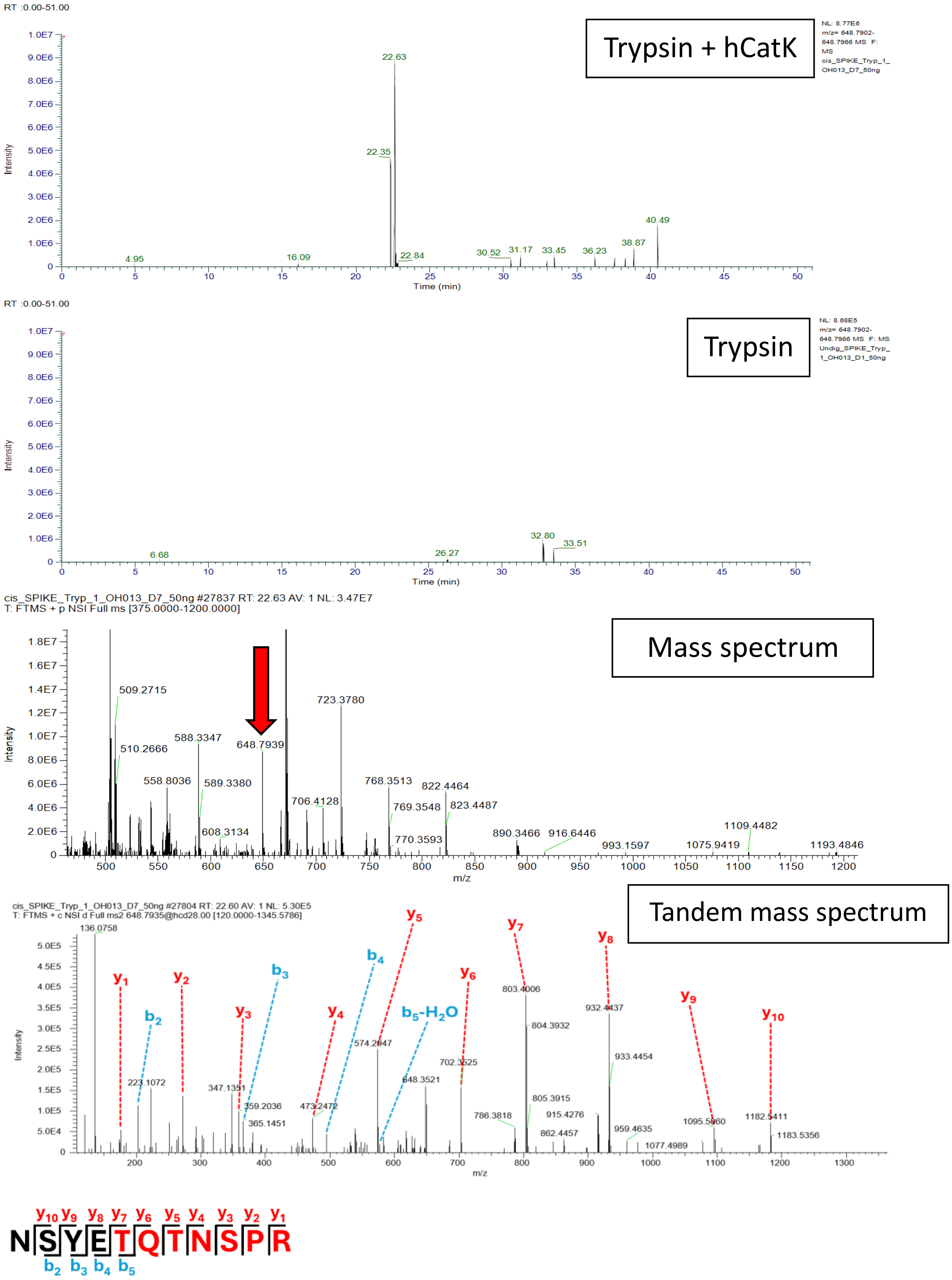


**Figure S19G. NSYETQTNSPR; Scan 27804; RT = 22.60; m/z = 648.7934. Spike *in cis*.**

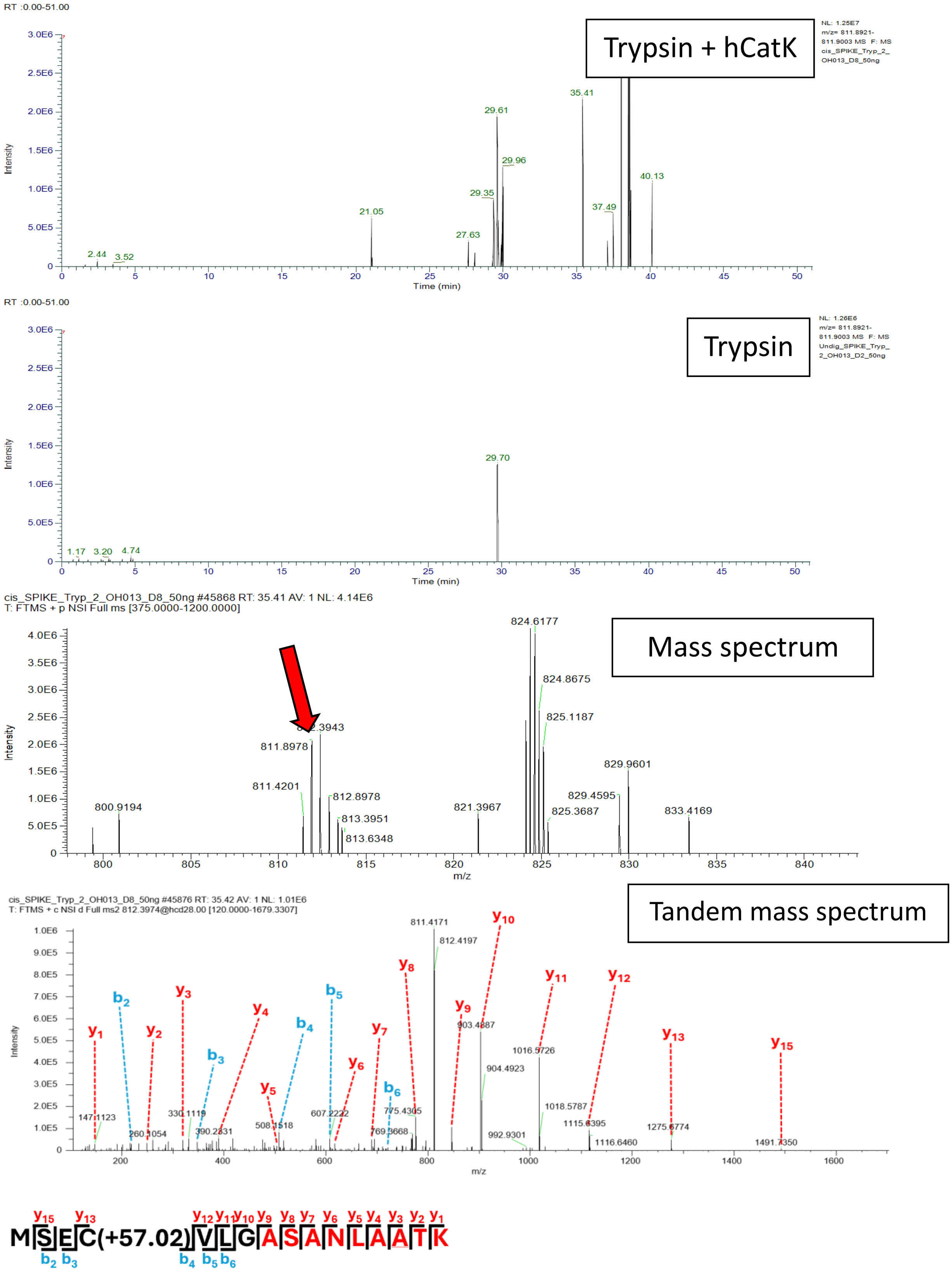


**Figure S19H.** **MSECVLGASANLAATK; Scan 45876; RT = 35.42; m/z = 811.8962. Spike *in cis*.**

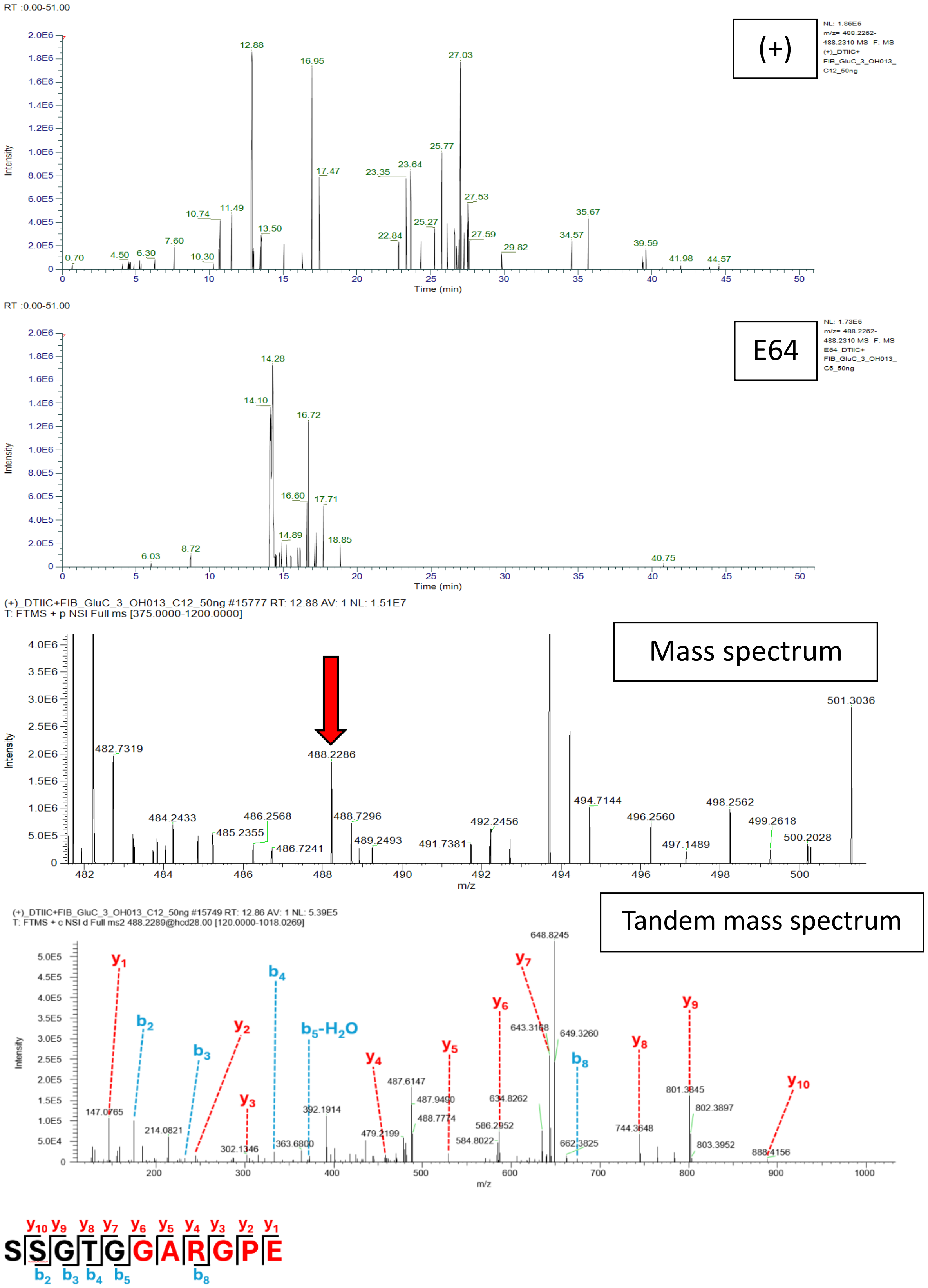


**Figure S20A. SSGTGGARGPE; Scan 15749; RT = 12.86; m/z = 488.2289. FIB and DTIIC *in trans*.**

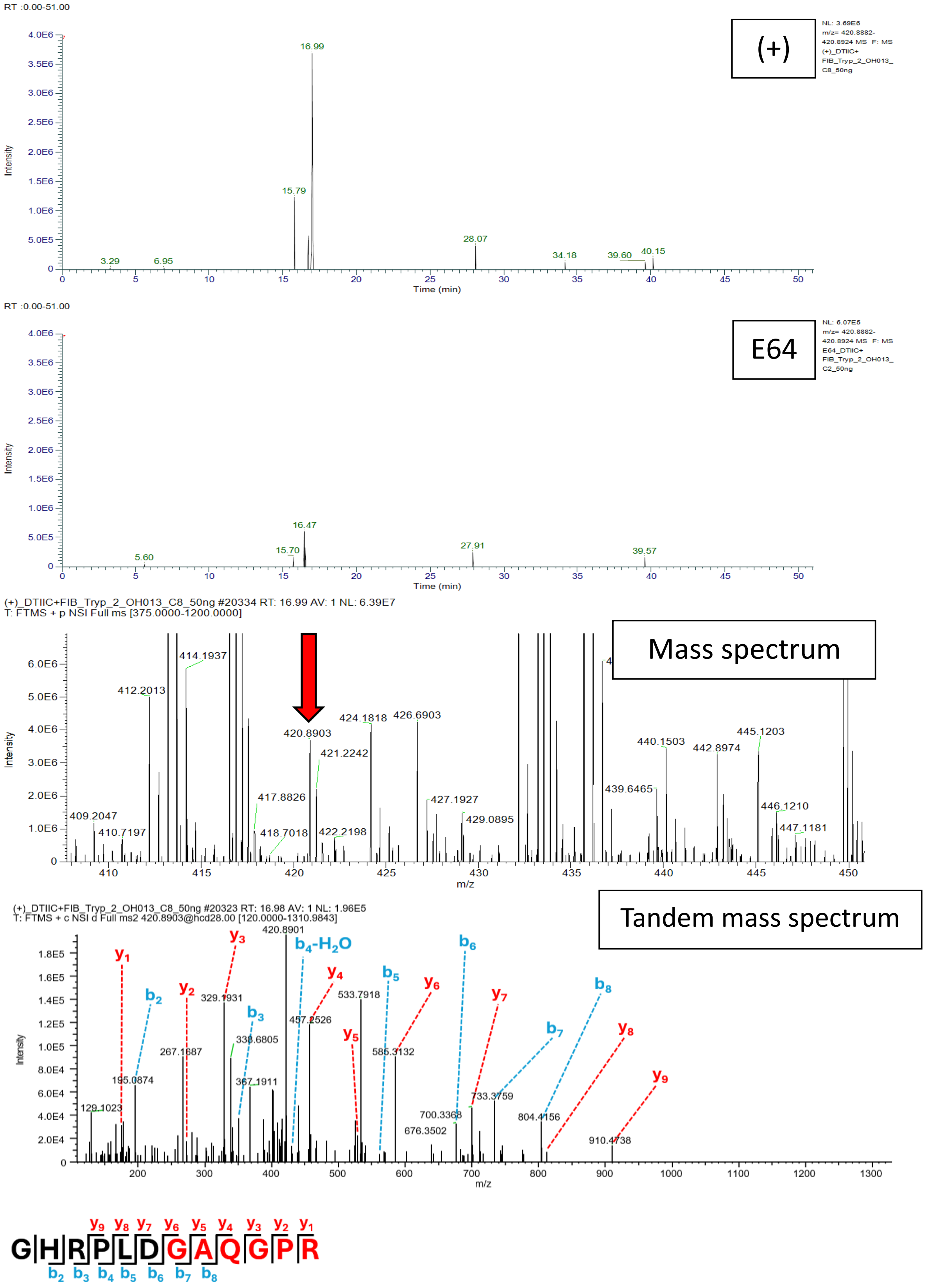


**Figure S20B. GHRPLDGAQGPR; Scan 20323; RT = 16.98; m/z = 420.8903. FIB and DTIIC *in trans*.**

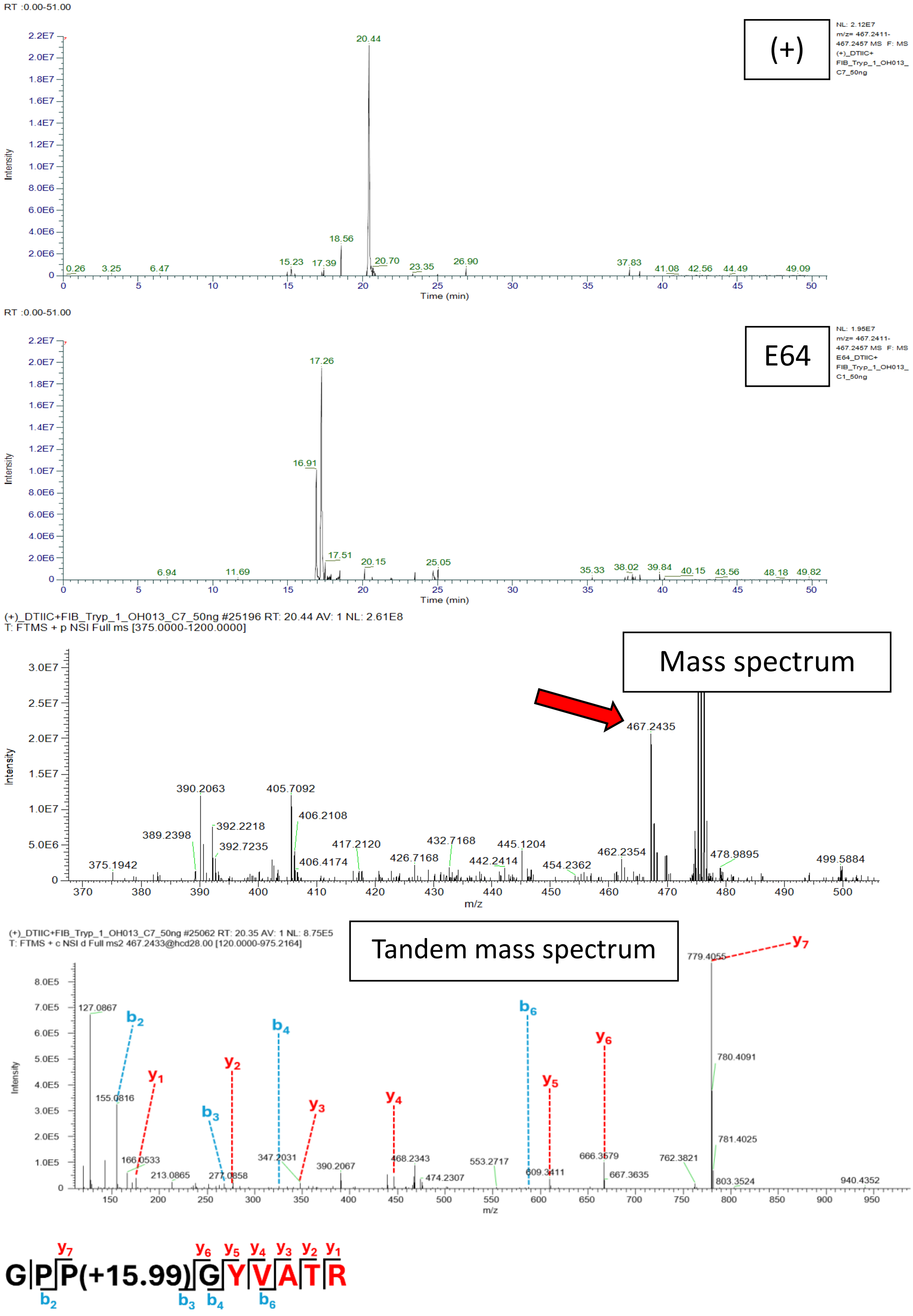


**Figure S20C. GPPGYVATR; Scan 25062; RT = 20.35; m/z = 467.2434. DTIIC and FIB *in trans*.**

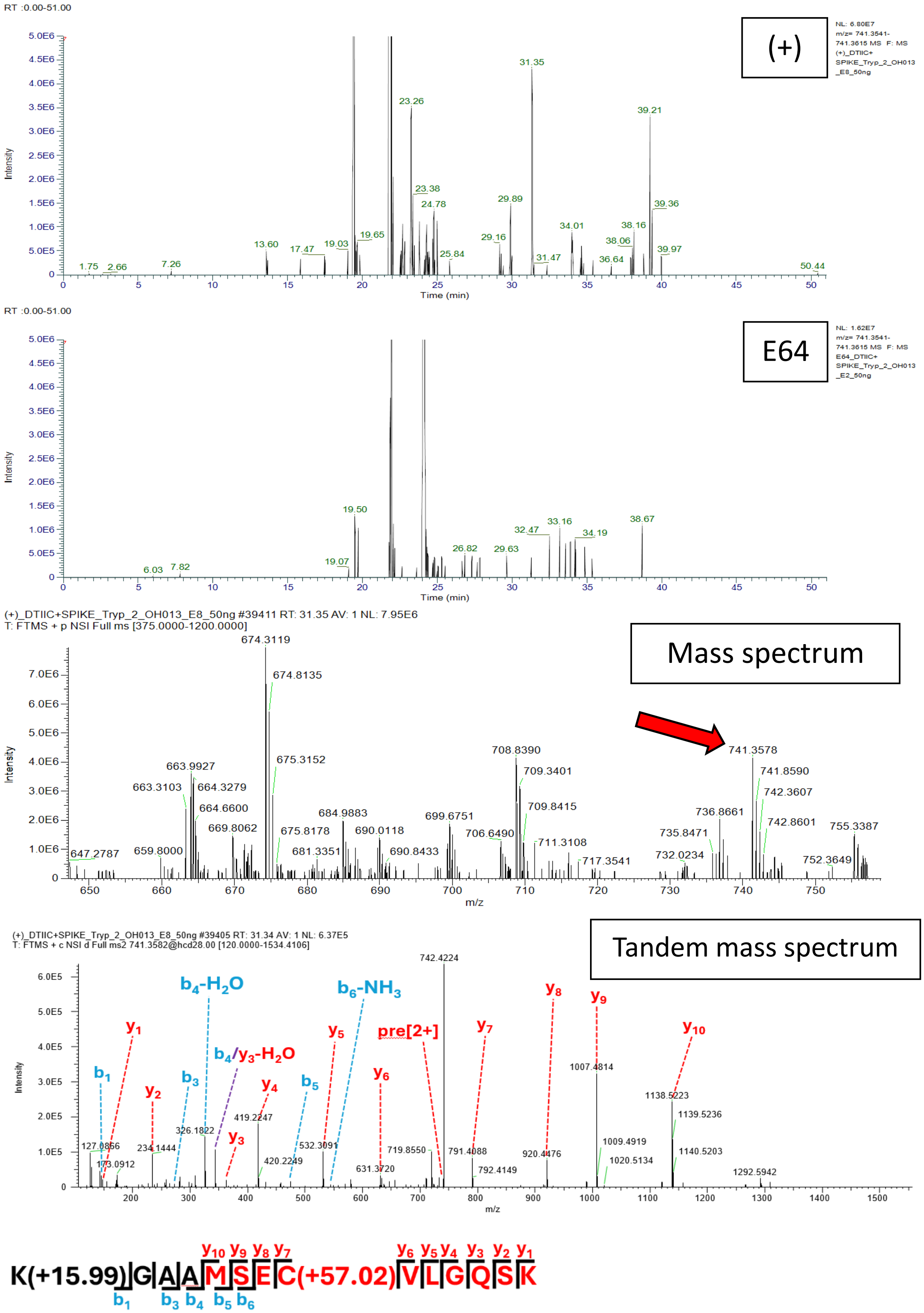


**Figure S20D.** **KGAAMSECVLGQSK; Scan 39405; RT = 31.35; m/z = 741.3578. DTIIC and Spike *in trans*.**

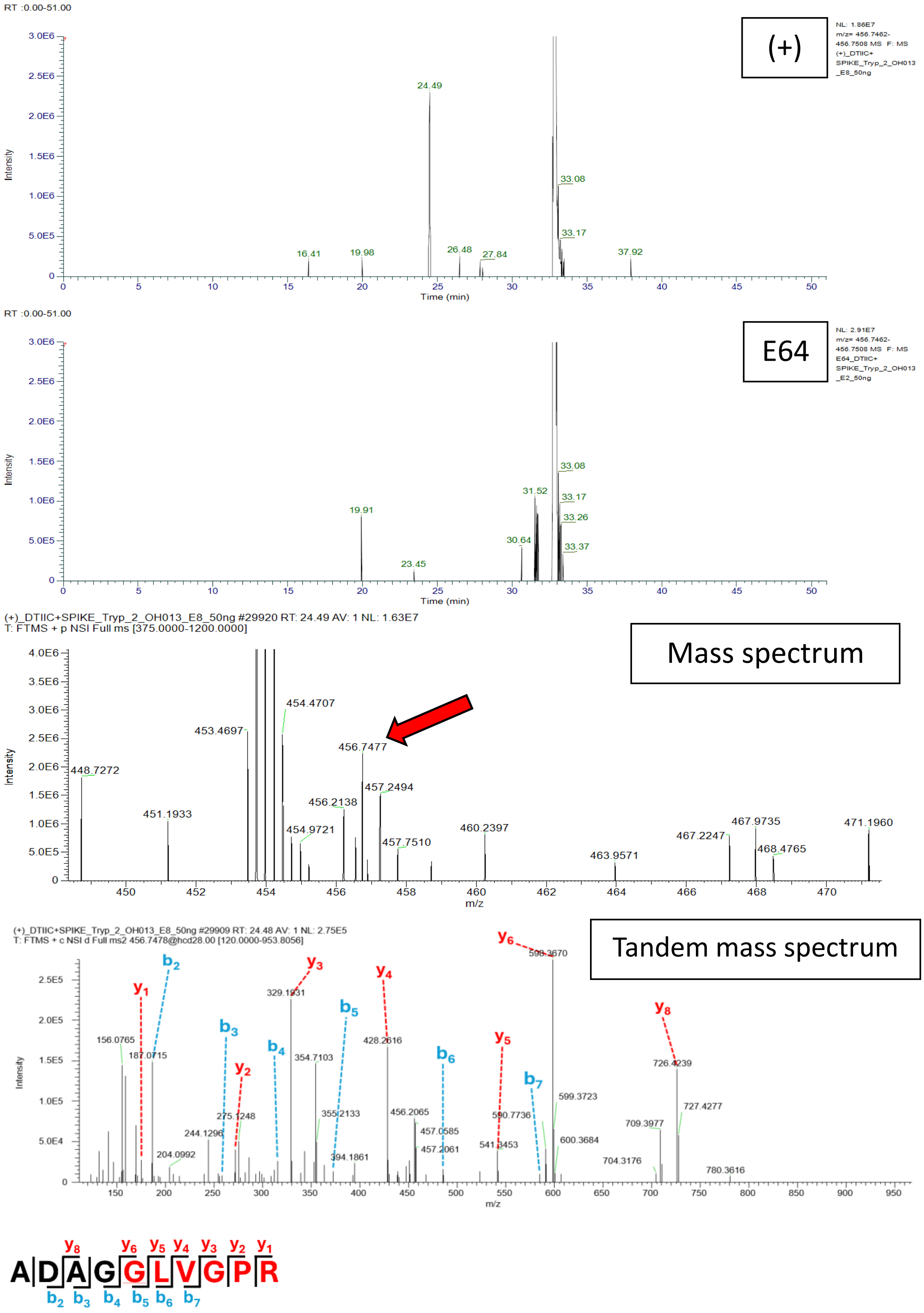


**Figure S20E. ADAGGLVGPR; Scan 29909; RT = 24.47; m/z = 456.7485. Spike and DTIIC *in trans*.**

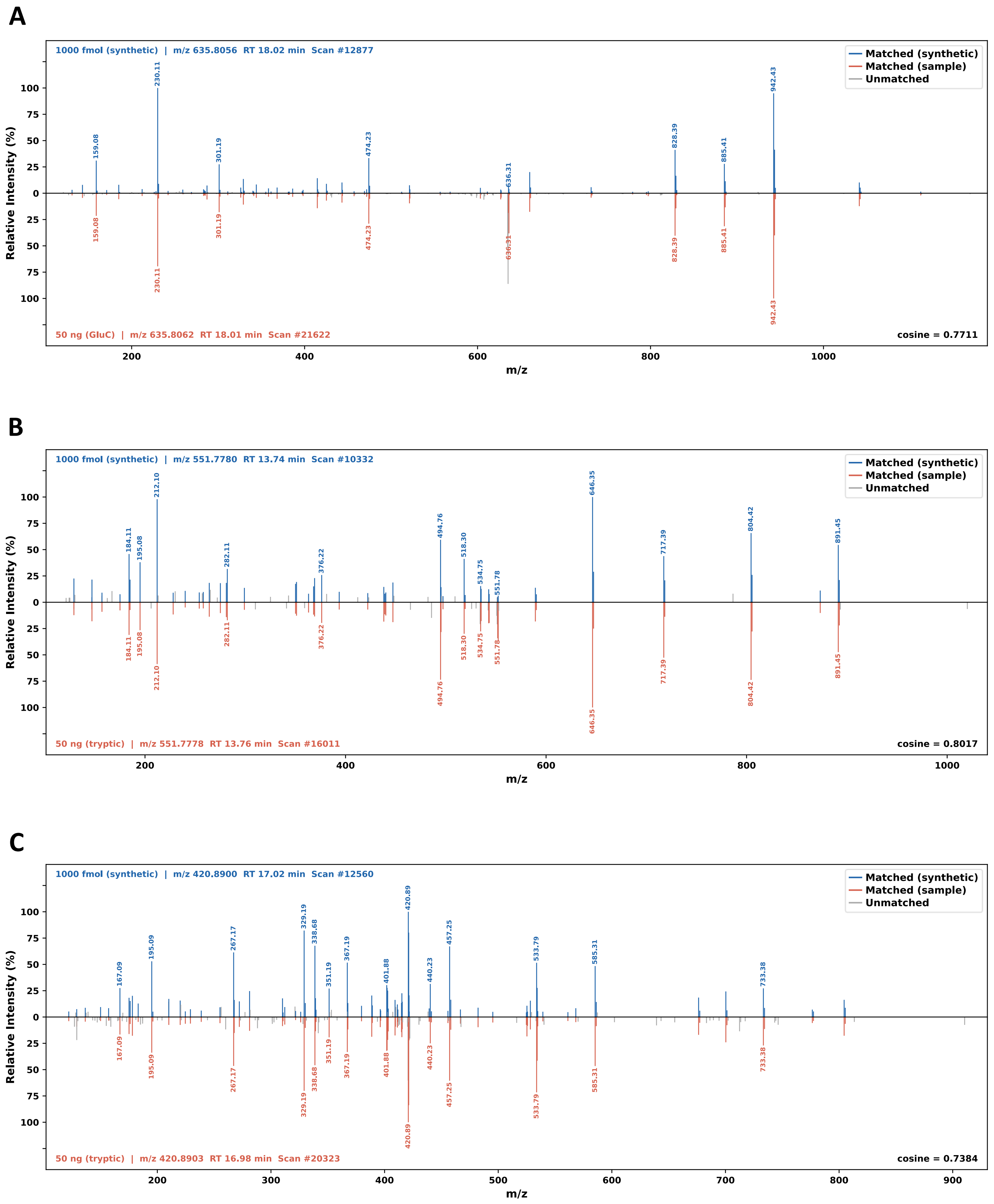


**Figure S21. Comparison between tandem mass spectra identified following hCatK activity on RA-related substrates to synthetic versions of spliced peptides.** Synthetic (top) vs observed following hCatK activity (bottom) tandem mass spectra for **A)** a DTIIC cis-spliced product, **TGAVGGPAGEP(+15.99)GRE**, **B)** a fibrinogen cis-spliced product, **NPSSAGAAATQK**, **C)** and a DTIIC-fibrinogen trans-spliced product, **GHRPLDGAQGPR**.
